# Evolutionary diversification of the biological screw joint in weevils

**DOI:** 10.64898/2026.08.21.746137

**Authors:** Jenny Hein, Julian Katzke, Alexander Riedel, Otto Bell, Alexandre Casadei-Ferreira, Angelica Cecilia, M. Lourdes Chamorro, Alexey Ershov, Tomáš Faragó, Elias Hamann, Chandan Sarkar, Sofiia Syrota, Clément Tavakoli, Nikolay Zagainov, Marcus Zuber, Tilo Baumbach, Michael Heethoff, Thomas van de Kamp

## Abstract

Complex biomechanical innovations are often treated as discrete evolutionary breakthroughs, yet their diversification within large radiations remains poorly understood. Beetle leg joints provide a striking example: some weevils possess screw-like coxa-trochanteral articulations in which rotation and axial displacement are mechanically coupled, resembling engineered screw-and-nut mechanisms. Whether these joints represent isolated mechanical extremes, discrete adaptive types or part of a broader continuum of phenotypic variation has remained unknown.

Here we combine synchrotron X-ray microtomography, landmark-free atlas-based morphometrics, quantitative functional morphology and phylogenetic comparative analyses to examine the mesocoxa-trochanteral joint in 68 species representing seven sampled family-level groups across early-diverging and derived weevil lineages. We show that screw joint evolution combines continuous variation in trochanteral shape with a restricted set of mechanically plausible joint-character combinations, rather than forming sharply separated morphological classes. True screw-and-nut joints are not confined to a distinct region of morphospace, indicating that overall form and mechanical configuration are not necessarily coupled. The occurrence of this configuration in the early diverging Caridae shows that it is not restricted to more derived families. Three-dimensional helix fitting revealed a mosaic geometry: winding angle showed the clearest relationship with the major shape axes (PC1-PC2), whereas axial pitch varied largely independently of shape, size and lineage. Together, these patterns show that screw joint components diversified with different degrees of evolutionary integration.

These results recast the weevil screw joint from a singular biomechanical curiosity into a diversified evolutionary system. These patterns are consistent with diversification through differential modification and retention of components with differing degrees of structural constraint, rather than a single shift from simple to fully specialized designs.

## Introduction

Biomechanical performance can shape the tempo, direction and functional scope of phenotypic diversification [1–4]. Yet we still know little about how the morphological structures that mediate such performance originate and diversify within large evolutionary radiations.

Beetles provide an exceptional system for studying these mechanisms. They are the most species-rich animal order and exhibit extraordinary morphological and ecological diversity [5–9]. A strongly sclerotized, functionally differentiated body plan is a defining feature of beetle morphology [10–12], while repeated ecological innovations have contributed to their exceptional diversification [9,13]. Fossil, molecular and genomic data have greatly advanced our understanding of beetle diversification [7,9]. Nevertheless, the functional implications of beetle evolution remain less well understood, particularly at the level of biomechanical systems [14,15].

Leg joints are central to locomotion, posture and interaction with the substrate. Although the jointed bauplan of the insect leg is broadly conserved, relatively small changes in joint architecture can alter how movements are guided, constrained and stabilized [16,17]. The coxa-trochanteral articulation is particularly relevant in this context because it lies at the proximal base of the leg, where small changes in joint architecture can influence how the distal leg is guided, stabilized and loaded. The insect coxa-trochanteral joint has often been treated as a simple hinge articulation [16,18]. In beetles, however, kinematic and functional-morphological studies have shown that this view is incomplete. The effective axis of rotation can shift during movement [17,18], and such joints include mechanically specialized systems ranging from spiral-ridge articulations in which complementary surfaces slide during rotation [19] to self-locking mechanisms [20].

Screw-like articulations represent one of the most striking examples of this mechanical diversity. Historical kinematic work already interpreted some coxa-trochanteral joints as screw-like mechanical pairs [21], and later studies identified screw-like articular geometries in several beetle groups, including the metacoxa-trochanteral joint of Mordellidae [22]. The most elaborate case described so far occurs in the weevil *Trigonopterus oblongus*, where a helical ridge on the trochanter engages with a corresponding internal guiding structure of the coxa, forming a biological screw-and-nut joint [23]. This articulation couples rotation with axial displacement and has been associated with leg locking and thanatosis in *Trigonopterus* [24]. This mechanically specialized structure provides a starting point for examining whether comparable joints across weevils form isolated biomechanical extremes, discrete adaptive types or a broader continuum of joint variation.

Although weevils (Curculionoidea) are extremely diverse and ecologically prominent (Fig. 1) [9,25], the distribution and morphological diversity of screw joints have not yet been systematically characterized across the superfamily. This limitation reflects the difficulty of resolving the internal three-dimensional geometry of compact, heavily sclerotized thoracic structures across large samples using dissection and light microscopy [14,24,26].

**Figure 1.**
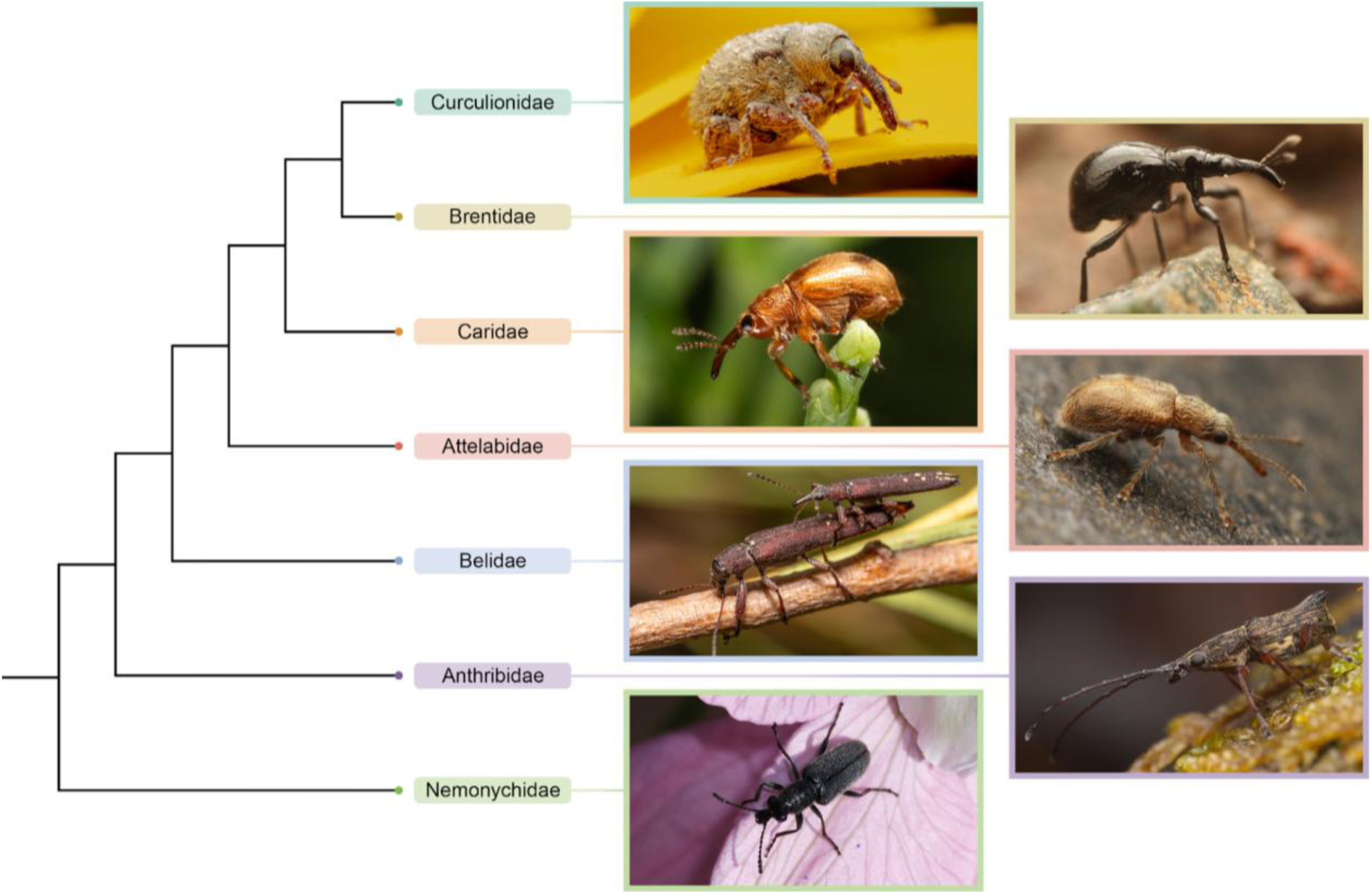
Phylogenetic and morphological diversity of Curculionoidea. Simplified family-level phylogenetic framework of the seven weevil families sampled in this study, shown together with representative live habitus images of Curculionidae (*Cionus olens*), Brentidae (*Alissapion* sp.), Caridae (*Car pini*), Attelabidae (*Metopum bryophagum*), Belidae (*Araiobelus filum*), Anthribidae (*Commista bispina*) and Nemonychidae (*Nemonyx lepturoides*). Tip and image-frame colours correspond to the family colour coding used throughout the manuscript.

X-ray microtomography (micro-CT) enables three-dimensional analysis of insect morphology [24,26–28], while advances in high-throughput imaging [28–30], AI-assisted segmentation [31–33] and shape-based morphometrics [34] now permit comparative quantification of complex internal joints on a broad taxonomic scale. We applied this framework to 68 specimens representing seven sampled family-level groups across early-diverging and derived curculionoid lineages, generating synchrotron micro-CT data and separately segmenting the coxa and trochanter of one mesocoxa-trochanteral joint per specimen. We selected the mesocoxa as a consistent sampling target primarily to exclude specialized locomotor or substrate-related modifications of the fore- or hind legs [16,35].

Landmark-free morphometrics focused on the compact, homologous trochanter, which directly bears the screw-related relief. The coxa was not analysed as a whole-shape object. Instead, we scored its local articular features, including the coxal thread, socket and wall opening, that guide or constrain trochanter movement. This avoids conflating screw joint morphology with broader variation in coxal form associated with thoracic integration and body-plan architecture [35].

Using the trochanter meshes, we performed landmark-free atlas-based diffeomorphic geometric morphometrics and constructed a principal-component morphospace to capture variation in trochanteral form. We used the coxal and trochanteral joint surfaces in parallel to classify recurring joint architectures and quantify screw-related functional geometry, including winding angle, axial span and fitted axial pitch. Finally, we assigned the 68 specimens to 15 taxonomic proxy tips on the available phylogenetic backbone (Supplementary Table 1) and integrated trochanter shape, joint-type classification and screw-geometry measurements with phylogenetic and ecological data. We used this combined dataset to test whether screw-like coxa-trochanteral joints form discrete mechanical categories or diversify across a continuum of shape and geometry, and how this diversity is structured by lineage history, size and broad ecological associations.

## Results and Discussion

### Morphospace structure

Across the first three principal components, trochanter shape occupied a broadly continuous morphospace with extensive overlap among families rather than forming discrete clusters (Fig. 2).

**Figure 2.**
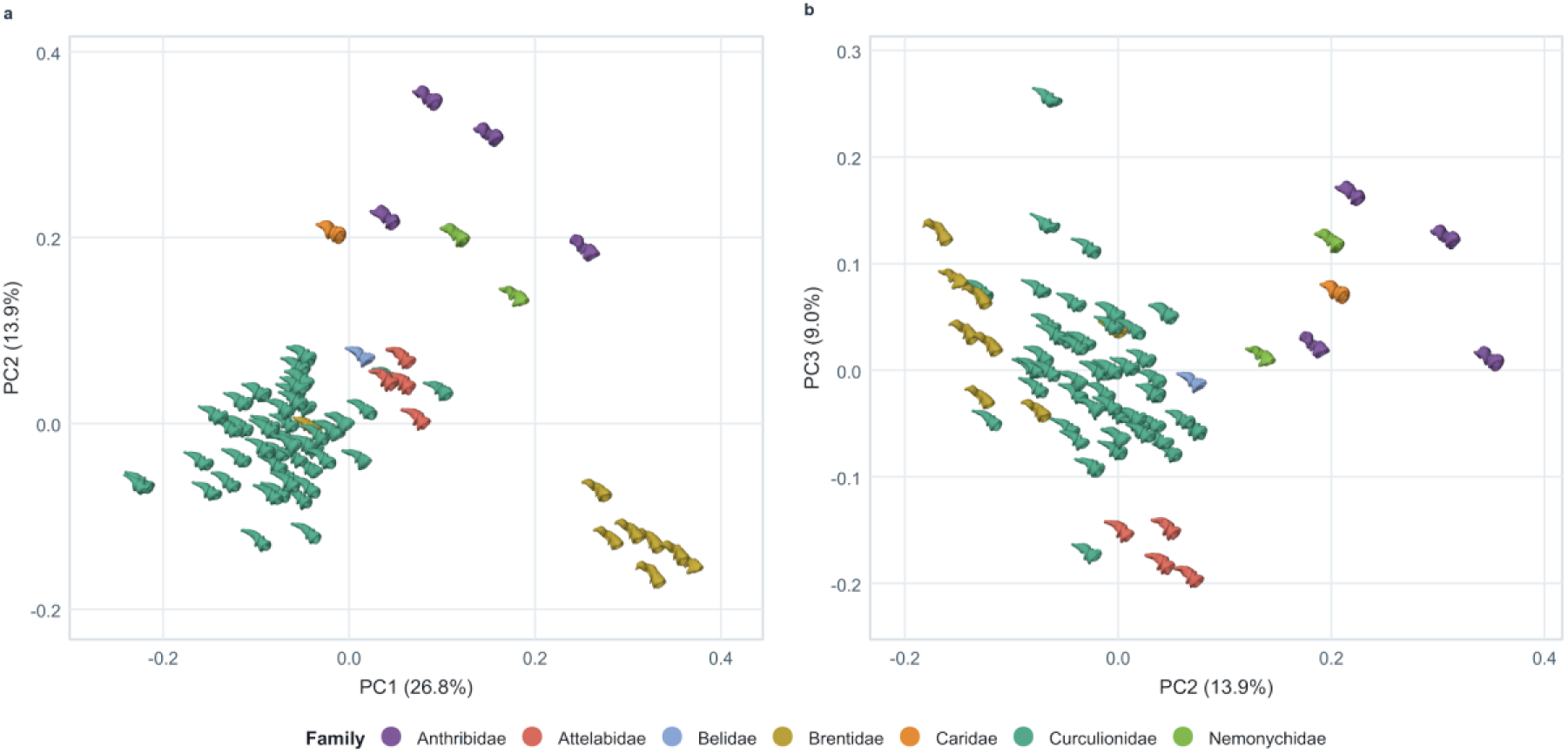
Two-dimensional projections of trochanteral morphospace. Trochanteral morphospace shown in two complementary projections, PC1-PC2 and PC2-PC3, with specimens coloured by family. These panels illustrate broad overlap among families, but also show that some lineages occupy relatively distinct portions of shape space in different principal dimensions. An interactive three-dimensional version of the morphospace is provided as <u>Supplementary Data 1</u>. Specimen-level PCA scores underlying the morphospace are provided in Supplementary Table 2.

We retained PC1-PC5 as a pragmatic major-shape subspace rather than applying a fixed cumulative-variance threshold. This cutoff retained the components that each contributed substantially to the atlas-derived morphospace, with PC5 still explaining approximately 5% of total shape variation (Supplementary Fig. 1; Supplementary Table 3). Together, these axes explained 61.4% of total shape variation and captured the major anatomically interpretable deformation modes, while avoiding overemphasis on increasingly minor higher-order components.

Reconstructions of the positive and negative extremes of PC1-PC5 show that the main axes of variation involve coordinated differences across the trochanter rather than isolated local changes. Along PC1 (Fig. 3), the negative extreme mainly shows a slightly narrower transition between the tapered process and the basal/articular region, whereas the positive extreme has a thicker and more strongly curved proximal process and less pronounced relief around the central articular region. PC2 contrasts an elongated, thinner overall form at the negative extreme with a shorter, more bulbous and compact configuration at the positive extreme. Together, PC1 and PC2 therefore capture changes in curvature and robustness of the proximal process, distal-body proportions and articular-surface relief. PC3 represents subtle changes extending across both the tapered process and the basal/articular body. At the positive extreme, the tip of the proximal process is thicker, slightly kinked and more curved than in the negative extreme. PC4 contrasts a negative extreme with a thicker, more strongly curved proximal process and marked displacement along the inner side of the distal rim against a positive extreme showing subtler deformation around the central articular relief. PC5 mainly captures localized variation around the distal/articular rim. The strongest changes occur in similar regions at both the negative and positive extremes, suggesting that PC5 describes opposite deformations of the same local articular structures rather than broader differences in overall trochanter shape.

**Figure 3.**
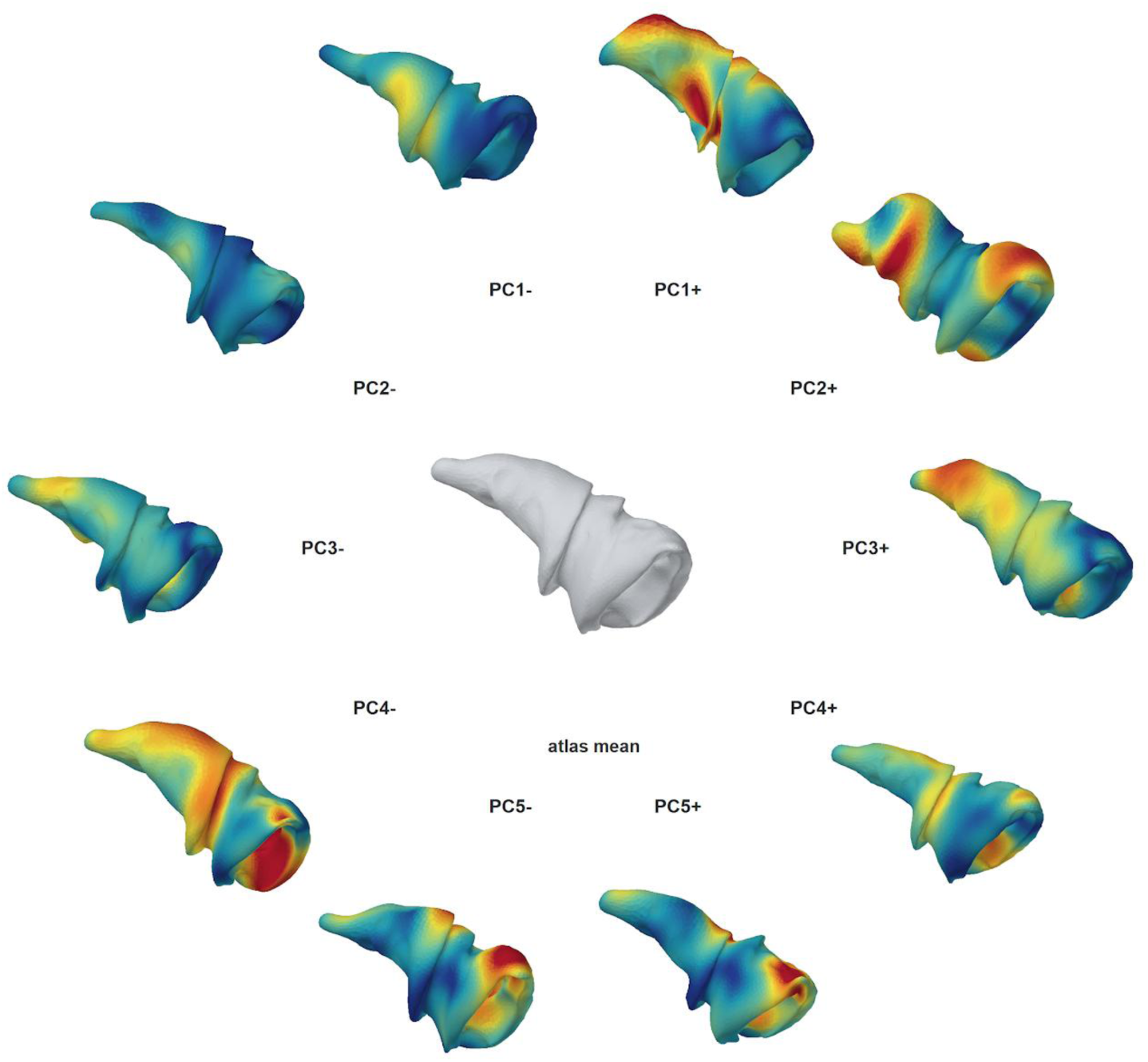
Major axes of trochanteral shape variation. Atlas-based reconstructions of the negative and positive extremes of PC1-PC5 are arranged around the atlas mean shape. Surface colours indicate vertex-wise displacement from the mean shape, with warmer colours marking regions of greater local deviation and cooler colours marking lower deviation. These reconstructions show the main anatomical regions contributing to the first five axes of trochanteral shape variation. Interactive three-dimensional versions of the heatmap reconstructions are provided as <u>Supplementary Data 2</u>, and a morphing animation through the PC extremes and atlas mean is provided as <u>Supplementary Video 1</u>.

With these deformation axes in view, the family-level distribution in morphospace can be interpreted as broad overlap with regional tendencies rather than discrete morphological separation. In the PC1-PC2 projection (Fig. 2a), Curculionidae formed a dense central core, while several other families extended into more peripheral regions. Brentidae were heterogeneous, with sampled specimens occupying both central and peripheral parts of the morphospace. Attelabidae and Belidae plotted close to the curculionid-dominated region, whereas Anthribidae, Nemonychidae and Caridae occupied more peripheral positions. Because both morphospace projections share PC2, the PC2-PC3 view mainly adds information about variation along PC3. In this projection, Attelabidae occupied a more peripheral position, some brentid specimens lay closer to the central curculionid region, and Curculionidae were less tightly clustered along PC3, with a few outlying specimens extending away from the central region. The curculionid outliers indicate substantial within-family shape diversification along PC3, while the broader shifts show that family-level structure is multidimensional rather than confined to the main PC1-PC2 projection. Even so, the families do not resolve into discrete, family-specific clusters.

To test whether the recovered morphospace structure depended on restricting the ordination to linear axes, we repeated the analysis using radial-basis-function kernel principal component analysis (RBF-kPCA) as a sensitivity analysis. With the previously used fixed kernel parameter (γ = 0.25), the first five kernel axes were nearly identical to the ordinary PCs (matched |r| = 0.999-1.000); a data-adaptive median-distance kernel likewise retained the broad family, joint-type and allometric structure (Supplementary Fig. 2; Supplementary Table 4). Thus, the principal biological patterns were not artefacts of the linear ordination, and ordinary PCA was retained as the primary, directly interpretable representation.

Repeated randomized Hopkins evaluations indicated departure from spatial randomness in standardized PC1-PC5 morphospace, but were treated only as an exploratory clusterability diagnostic because the sample of 68 specimens did not permit conventional large-sample conditions for formal Hopkins inference (mean H = 0.906; Supplementary Fig. 3, Supplementary Table 5). Clustering analyses failed to recover stable or well-separated morphological groups, as cluster assignments varied with both method and parameter choice and therefore provided no support for discrete structure in morphospace (Supplementary Fig. 4, Supplementary Table 6). Descriptive estimates based on mean squared Euclidean distances ranked Anthribidae highest, followed by Brentidae and Curculionidae, whereas Attelabidae had the lowest estimate (Extended Data Fig. 1; Supplementary Table 7). A separate bias-adjusted PERMDISP analysis of unsquared Euclidean distances to family centroids supported differences in multivariate dispersion among the four families represented by at least three specimens (F = 6.95, P = 0.0038). Pairwise PERMDISP comparisons adjusted for the false discovery rate (FDR) supported greater dispersion in Anthribidae than in Attelabidae (adjusted P = 0.0006) and Curculionidae (adjusted P = 0.0066). The remaining pairwise differences did not remain significant. Rarefaction of the mean-squared-distance calculation to four specimens per family preserved the same ranking, indicating that the descriptive pattern was not solely driven by unequal sample size, although estimates for the small families remain uncertain (Extended Data Fig. 1; Supplementary Table 7).

Taken together, these results show that trochanteral morphology is distributed continuously across morphospace, but not in a completely unstructured way. This pattern is consistent with broad-scale morphometric studies showing that complex structures often vary along continuous axes shaped by coordinated changes among anatomical features and lineage-specific differences, rather than by changes confined to single isolated features or regions [36–38]. In our dataset, family identity was associated with differences in morphospace position and within-family disparity, but did not yield discrete clusters because families spanned both central and peripheral regions of shape space.

### Joint typology

Although trochanter shape is continuously distributed across morphospace, the structural organization of the coxa-trochanteral articulation can still be grouped into a limited number of recurring mechanical configurations. Across the taxa sampled (with a few exceptions in Anthribidae; see paragraph “Atypical joint-character combinations”), four principal joint types were recognized: a true screw-and-nut joint (Fig. 4a), an unopposed screw configuration (Fig. 4b), a socket-guided rotational joint (Fig. 4c), and an unguided rotational joint (Fig. 4d).

**Figure 4.**
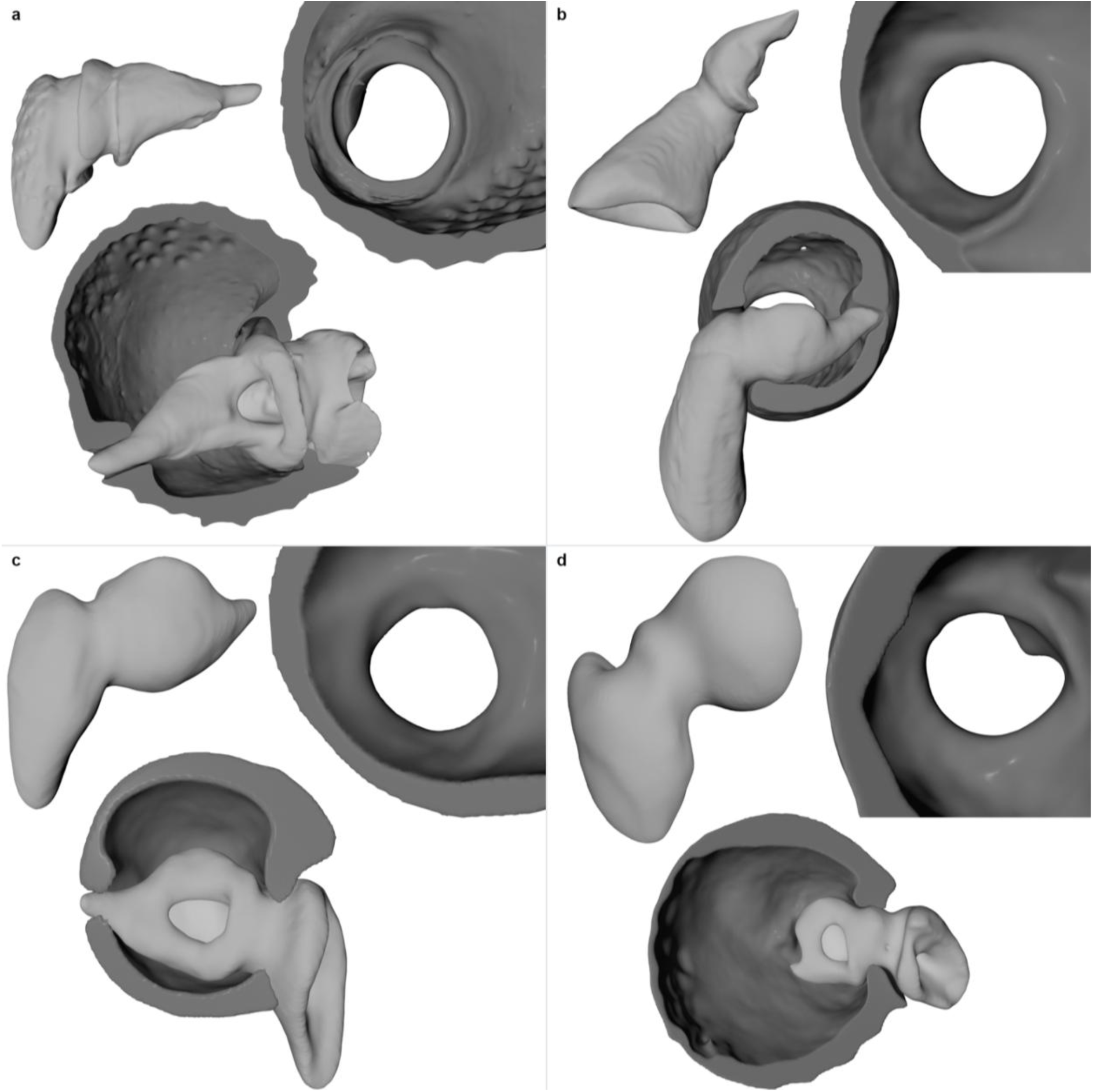
Structural typology of the weevil coxa-trochanteral joint. Representative three-dimensional reconstructions of the four joint configurations recognized in this study: **(a)** true screw-and-nut joint, **(b)** unopposed screw configuration, **(c)** socket-guided rotational joint and **(d)** unguided rotational joint. The panels show *Acalles fallax*, *Nanophyes marmoratus*, *Proterhinus* sp. and *Ormiscus saltator*, respectively. Trochanters are shown in light grey and coxae in dark grey.

In the true screw-and-nut joint (Fig. 4a), a well-developed spiral articulation on the trochanter is matched by a corresponding coxal internal thread. In the unopposed screw configuration (Fig. 4b), trochanteral winding is present, but the coxa lacks a complementary element, leaving the screw articulation incomplete. The socket-guided rotational joint (Fig. 4c) lacks a developed screw geometry, yet trochanter movement is still guided by a socket in the coxal wall. The unguided rotational joint (Fig. 4d), by contrast, lacks both a screw articulation and an internal guiding socket and therefore represents the least internally constrained configuration observed. In the specimens assigned to this joint type, however, movement is not entirely unconstrained, because an external spur-like projection of the coxa may provide an additional mechanical stop. This structure corresponds to the coxal posterior projection described by Chamorro-Lacayo and Konstantinov [19] and likely constrains movement externally rather than through internal coxal guidance or trochanteral winding. Despite these structural differences, the four joint types overlapped broadly in trochanteral morphospace (Supplementary Fig. 5), indicating that overall trochanter shape alone does not distinguish among the four joint architectures.

Across the four binary joint characters scored here, the sampled morphologies occupied only a restricted part of the combinatorial design space, defined by all possible presence-absence combinations of trochanteral helix, coxal guide, socket and coxal wall opening (Fig. 4, Extended Data Fig. 2, Supplementary Table 8). This restriction was structured rather than random. The clearest constraint was the complete absence of coxal winding without a corresponding trochanteral helix. Mechanically, this asymmetry is consistent with the hypothesis that a coxal thread can function as a guide only when a matching helical relief on the trochanter engages with it. In a true screw-and-nut joint, this paired geometry could constrain the trochanter to a more defined rotational-translational path, distribute contact across complementary surfaces and increase resistance to lateral displacement under load. A trochanteral helix without a complementary coxal thread forms an unopposed screw configuration and cannot impose the same internally guided rotational-translational path. In all sampled unopposed configurations, however, the coxal socket still receives the coxa-facing end of the trochanter and could reduce lateral play, help preserve rotational alignment and resist withdrawal or dislocation under load. In an unopposed screw, the raised trochanteral helix is positioned on the inner side of the rim of the apical coxal opening. Under axial loading, the helix may press against this rim and thereby help retain the trochanter within the coxa. This geometry could give the joint greater pull-out resistance than an unguided rotational joint, even without the continuous form-locked guidance of a true screw-and-nut joint, but that prediction was not tested here. An unopposed screw therefore retains greater skeletal constraint than an unguided rotational joint, which lacks both a complementary coxal thread and an internal guiding socket. Several other alternative configurations were likewise not observed, consistent with the hypothesis that screw joint diversification is channelled by mechanical interdependence among paired articular surfaces rather than producing all theoretically possible character combinations. However, as shown in other densely sampled morphometric systems, broader sampling may reveal rare or transitional forms that alter the apparent discreteness of complex morphological variation [39,40].

We defined the coxal wall opening as a discrete perforation through the wall of the coxal socket adjacent to the coxa-facing trochanteral apex, distinct from the socket cavity itself and from artefactual breaks in the cuticle. Because its presence did not define a distinct mechanical architecture, we treated it as an accessory structural character rather than as an additional joint type, resulting in seven observed character combinations (Extended Data Fig. 2, Supplementary Table 8). In other cucujiform beetles, comparable openings occur at the termination of the coxal suture and accommodate or expose this part of the trochanter. In some early-diverging weevils, the external procoxal suture is strongly reduced or obliterated while the opening remains, indicating that the opening can persist as a distinct component of the coxa-trochanter interface even after the external suture is no longer visible [16,41]. These comparative descriptions primarily concern the procoxa, so we regard the mesocoxal opening observed here as morphologically comparable without assuming strict homology across thoracic segments. Because a wall perforation may occur preferentially in generally thin-walled coxae, and because very thin cuticle is especially susceptible to apparent discontinuities during imaging and segmentation, we tested whether opening-bearing coxae had lower whole-volume three-dimensional wall thickness after accounting for overall coxa size. Wall thickness increased strongly with overall coxa size (R² = 0.534, P < 0.001), whereas coxa size itself did not predict opening state (P = 0.803). In the primary whole-volume analysis, opening-bearing coxae had 11.5% lower median wall thickness than coxae without an opening, but this contrast was not statistically significant (P = 0.116); the lower-tail metric showed the same direction (P = 0.158; Supplementary Fig. 6, Supplementary Tables 9-11). The opening may accommodate an elongated trochanteral tip and extend its guidance through the coxal wall, thereby constraining lateral displacement. In *Trigonopterus*, passage of the trochanteral tip through a comparable opening stabilizes the rotational axis and protects the joint against jamming [23]. The opening showed no consistent association with the four joint types. We therefore retained it as an accessory structural character and interpreted the phylogenetic distribution and associated trait contrasts descriptively (Supplementary Figs. 7 and 8), rather than as evidence for a separate functional joint category.

The distribution of joint types across families was similarly uneven (Fig. 4, Extended Data Fig. 2, Supplementary Fig. 5c, Supplementary Table 12). Screw-dominated configurations occurred mainly in derived curculionoid lineages, whereas socket-guided or purely rotational configurations were more common among several early-diverging families. At the same time, joint-type composition was not strictly family-specific, because some families included more than one configuration and the same joint type occurred in phylogenetically distant parts of the tree. Taxonomic grouping did not map directly onto joint type, indicating that repeated modification or differential retention of joint components has shaped the diversification of the coxa-trochanteral articulation.

The single representative of Caridae is particularly notable because it lies outside the curculionid-dominated region of morphospace, close to several early-diverging families, yet possesses a true screw-and-nut joint type. Given the phylogenetic position of Caridae between early-diverging and more derived curculionoid lineages [42–44], this observation shows that a complete screw-and-nut architecture is not restricted to the derived curculionid shape cluster. It is compatible with either an older origin of screw-like articulation followed by subsequent modification or loss, or an independent origin of a comparable mechanism in Caridae. Distinguishing between these alternatives cannot be achieved from shape-based ancestral-state reconstructions alone, because overall trochanter shape and joint type are not tightly coupled in our dataset. Testing the origin of this configuration would require denser sampling of Caridae and other early-diverging families, explicit ancestral-state reconstructions of the individual joint characters, and, where preservation permits, evidence from fossil joints. These alternative histories need to be interpreted within a broader macroevolutionary setting. Beetles underwent extensive diversification during the Cretaceous terrestrial revolution [45], providing a broad historical context for considering the gain, modification, loss or recombination of joint components. The Caridae specimen extends the observed distribution of the complete screw-and-nut architecture, but does not resolve the number, timing or direction of evolutionary transitions.

### Atypical joint-character combinations

Two of four anthribid specimens in the dataset showed joint-character combinations that did not fit into the four recurrent structural categories. In *Anthribus nebulosus*, the trochanter showed only a weak and ambiguous screw-like surface relief, much less developed than in clearly screw-bearing specimens and insufficient to form a fully expressed screw articulation. *Anthribus nebulosus* and *Urodon rufipes* both lacked a true coxal guiding socket. Instead, the coxal wall carried a unilateral internal edge or stop-like structure (Supplementary Fig. 9). Mechanically, this structure differs from a socket because it provides no continuous enclosure or guidance of the trochanteral tip. At most, it could act as a unilateral contact surface if the trochanter moves towards the coxal wall at extreme joint positions, but its position along only one side of the coxal wall does not indicate a screw-like guiding function. In the absence of strong skeletal guidance, movement and positional stability in these joints may therefore depend more strongly on soft-tissue constraints, including muscles, tendons and joint membranes. The occurrence of fully developed guiding sockets in both earlier-diverging and more derived lineages argues against interpreting these unilateral stops as intermediate stages. We therefore treated the two specimens as atypical variants outside the main typological framework and excluded them from group-wise joint-type statistics, but retained them as descriptive observations because they reveal structural variation around this framework.

### Screw joint geometry

Screw joint geometry (Fig. 5) varied substantially across taxa. Four of the 68 specimens lacked a discernible helical ridge and therefore provided no trajectory for fitting. Robust circular three-dimensional helix fits converged for all 64 traced trajectories. One trajectory exceeded the relative-RMS threshold used to define the main dataset, leaving 63 specimens (Supplementary Table 13). Other diagnostic warnings were retained for quality control but did not define an exclusion subset, because point count and angular coverage partly reflect the biological extent of short helices rather than fit failure.

**Figure 5.**
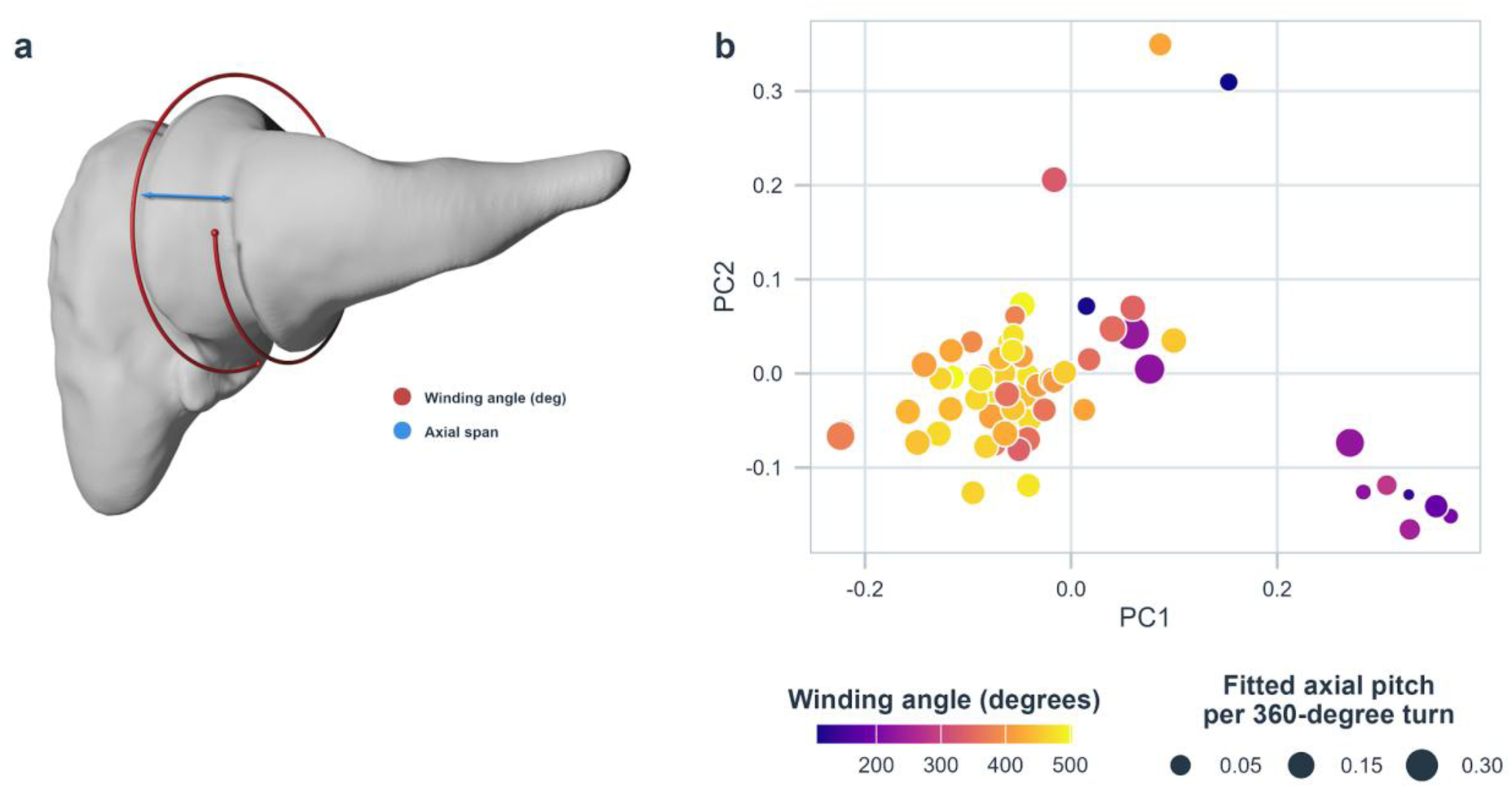
Quantification of screw joint geometry. (a) Three-dimensional trochanter reconstruction illustrating the principal geometric descriptors of the screw-like articulation. The red curve schematically represents the helical trajectory used to quantify absolute winding angle, and the blue arrow indicates axial span along the screw axis. (b) PC1-PC2 trochanteral morphospace for the 63 trajectories in the main dataset. Points represent individual specimens; colour indicates absolute fitted winding angle and point size indicates fitted axial pitch per 360° turn. Linear measurements are expressed in centroid-size-normalized coordinates. Conditional-bootstrap results used to assess measurement sensitivity are reported in Supplementary Tables 14-15.

In the main dataset, absolute winding angle covaried strongly with PC1-PC2 shape (R² = 0.512, F2,60 = 31.46, P = 4.52 × 10⁻¹⁰), driven by PC1 (P = 6.43 × 10⁻¹¹), and all 200 conditional-bootstrap analyses remained significant. Fitted axial pitch showed only marginal support (R² = 0.097, P = 0.046) and was significant in only 55.5% of bootstrap draws. Screw geometry is therefore heterogeneous: winding angle covaries strongly with the major shape axes (PC1-PC2), which together account for 40.6% of atlas-derived shape variation, whereas evidence for a shape-pitch relationship is weak and sensitive to conditional measurement uncertainty. The corresponding diagnostics are shown in Supplementary Fig. 10.

Axial span also covaried strongly with PC1-PC2 shape in the main dataset (n = 63, R² = 0.476, F2,60 = 27.23, P = 3.86 × 10⁻⁹), driven by PC1 (P = 6.25 × 10⁻¹⁰), and all 200 conditional-bootstrap fits were significant. Because axial span is geometrically coupled to winding angle and the length of the traced helical relief, this model is not independent evidence for a separate functional association. Winding angle and axial span were strongly related in the main dataset (n = 63, R² = 0.504, F1,61 = 61.95, P = 7.34 × 10⁻¹¹; Supplementary Fig. 11, Supplementary Table 15). Fitted pitch showed no robust shape association. At proxy-tip level under the Grafen primary tree, the PC1-span slope was nominally significant (n = 14, raw P = 0.039, FDR-adjusted P = 0.177; Supplementary Fig. 12 and Supplementary Table 16), but its significance changed across tree and leave-one-out analyses. Thus, coordination among axial extent, winding and shape is clear at specimen level, whereas the corresponding proxy-level phylogenetic association is sensitive to tree specification and individual tips. These results do not establish mechanically or evolutionarily independent modules.

Comparisons between structural joint types were limited by highly unequal sample sizes (Supplementary Fig. 13; Supplementary Table 17). The main dataset contained 57 true screw-and-nut joints but only three unopposed screw configurations. Winding angle differed nominally (Kruskal-Wallis P = 0.0202) but not after correction across the three univariate tests (adjusted P = 0.0605); fitted pitch (P = 0.387), axial span (P = 0.317) and combined geometry (permutational multivariate analysis of variance (PERMANOVA), P = 0.052; conditional-bootstrap median P = 0.070) were not supported. These descriptors characterize the traced joint geometry rather than realized motion, and none represents the in-vivo range of motion, which also depends on soft tissues. Joint-type patterns are therefore descriptive hypotheses for kinematic or biomechanical testing, not general group effects.

### Allometry

Allometric scaling explained a statistically detectable but limited component of trochanter shape variation (Extended Data Fig. 3, Supplementary Tables 18-22). Because atlas shapes were normalized for scale before PCA, isometry predicts no systematic association between shape and log centroid size. A multivariate regression based on randomization of residuals in a permutation procedure (RRPP) using all 67 non-zero atlas PCs rejected this null, but size explained only 7.0% of total atlas-derived shape variation (R² = 0.070, F = 4.98, Z = 3.52, P = 0.0001). A PC1-PC5 sensitivity analysis yielded R² = 0.106, but this estimate applies only to that retained subspace. Across individual axes, PC2 was the only component that remained significant after Holm correction across all 67 axes (R² = 0.279, adjusted P < 0.001; Supplementary Table 18). PC1 had a weaker raw association (R² = 0.087, P = 0.015) that did not survive correction. PC1-PC5 nevertheless accounted for 92.6% of the fitted allometric shape change, so these axes retain anatomical interpretability without defining the scope of the full-shape test. Larger trochanters tended towards lower PC2 values, corresponding to a more elongate and less bulbous overall shape (Fig. 3), whereas most localized higher-order variation was largely size-independent. The concentration of size effects along only one major axis is consistent with allometry acting as a direction-specific component of shape variation rather than as a uniform scaling response [46,47].

After accounting for shared ancestry, direct multivariate phylogenetic RRPP across all 67 non-zero PCs did not detect a supported shape-size association at the 15 aggregated taxonomic proxy tips (R² = 0.099, F = 1.42, Z = 0.98, P = 0.157; Extended Data Fig. 3a, Supplementary Table 22). Axis-specific phylogenetic generalized least-squares (PGLS) relationships likewise did not remain significant after FDR correction. At specimen level, absolute winding angle increased with centroid size in the main dataset (n = 63, R² = 0.150, raw P = 0.00169, Holm-adjusted P = 0.0305; Supplementary Fig. 14). Axial span showed a nominal association with size (raw P = 0.044), but this did not survive Holm correction (adjusted P = 0.620), and fitted pitch showed no evidence of an association (raw P = 0.611). No geometry-size PGLS association survived correction. Shape allometry was therefore clear at specimen level but unsupported in the phylogenetic multivariate test, whereas winding-angle allometry was modest and likewise lacked corrected phylogenetic support.

### Phylogenetic analyses

Shape-only comparative analyses used 15 aggregated taxonomic proxy tips, whereas exploratory geometry analyses used 14 tips from the main dataset. To keep trait sampling identical, the matched geometry analyses recalculated PC1 and PC2 tip means from the same geometry-bearing specimens before aggregation. They are therefore distinct from shape-only signal analyses using all 15 tips. PC1 and PC2 retained phylogenetic signal in the 15-tip shape-only dataset. Geometry signal estimates and matched shape-geometry analyses used the 14 tips with adequate geometry data. Absolute winding angle also showed signal (Blomberg’s K = 0.90, FDR-adjusted P = 0.024), whereas axial span did not (Extended Data Fig. 4). Fitted axial pitch likewise showed no corrected signal (Supplementary Table 23). This proxy-level correspondence of winding angle remains conditional on sparse sampling and broader taxonomic assignments and cannot be generalized to species- or genus-level signal. Ancestral-state reconstructions on the proxy tree should not be read as direct reconstructions for the sampled genera or Curculionoidea as a whole (Extended Data Fig. 5). All reported continuous reconstructions are conditional on a Brownian-motion (BM) model used for estimation; this is a reconstruction assumption, not a model-selection result. In particular, OU was top-ranked for axial span in the exploratory univariate comparison on the primary tree, but rankings were tree-dependent and did not distinguish a general evolutionary process. Under the BM reconstruction on the Grafen primary tree, root estimates in the 14-tip main geometry dataset were an absolute winding angle of 299.0° (95% CI, 179.6–418.3°) and axial span of 0.159 (95% CI, 0.048–0.270). Broad intervals, sparse sampling and broader proxy assignments preclude precise anatomical inference from either the shape or geometry reconstructions. Raw-scale node estimates and intervals for all displayed traits are provided in Supplementary Table 24; root summaries across 15 tree representations are provided in Supplementary Table 33.

The PC1-PC2 proxy-tip phylomorphospace showed separation among several mapped groups. Proxy tips representing Brentidae and Curculionidae extended towards opposite ends of PC1, while those used for Nemonychidae and Anthribidae occupied the positive PC1-PC2 region and the Attelabidae and Belidae proxies remained closer to the reconstructed root (Supplementary Fig. 15). The positive region corresponds to comparatively robust and curved proximal processes and compact overall forms, whereas separation along PC1 primarily involves proximal-process curvature and robustness and central articular relief.

PC3-PC5 provided no consistent proxy-level pattern. At the 14 tips represented in the main dataset, winding angle showed significant phylogenetic signal, whereas axial span and pitch did not (Extended Data Figs. 4-5). Within the limitations of proxy assignment, sparse tips and within-tip averaging, broad shape corresponded more consistently to the Grafen primary tree than did most robust screw-geometry estimates.

Univariate evolutionary-model rankings were conditional on tree choice. In the main dataset, Brownian motion had the highest small-sample-corrected Akaike information criterion (AICc) weight for winding angle (0.56) and fitted pitch (0.47), whereas Ornstein-Uhlenbeck was top-ranked for axial span (0.78). Rankings changed across tree variants. These small-sample comparisons therefore do not distinguish a general evolutionary process and are reported as exploratory summaries (Supplementary Table 25).

Multivariate evolutionary-model fits for trochanteral shape were not used for biological inference. The main-dataset comparison used 15 shape-only proxy tips. For PC1-PC5, Brownian motion and early burst produced reliable fits and Brownian motion had lower AIC (ΔAIC = 0 versus 2.79). The OU optimizer converged and its raw AIC was lower than Brownian motion, but mvMORPH reported an unreliable Hessian solution, so OU was excluded from the reliable-fit ranking. For PC1-PC4, Brownian motion and early burst were reliable and Brownian motion again had lower AIC (ΔAIC = 0 versus 3.24), whereas the OU fit failed. Brownian motion was therefore top-ranked among reliable fits for both shape representations, but these incomplete small-sample comparisons do not distinguish a general evolutionary process (Extended Data Fig. 6; Supplementary Table 26). We therefore do not infer a clade-wide mode of trochanteral-shape evolution from these fits.

Given the small proxy-level sample, key comparative analyses were repeated across 15 topology and branch-length representations. For PC1 versus winding angle, slopes were negative on all 15 trees in the main dataset, but only one test was nominally significant and none survived FDR correction. PC1 versus axial span was negative on all trees but was significant only on some representations and changed under leave-one-tip-out refitting. Pitch relationships were similarly inconsistent. The phylogenetic multivariate-allometry result changed from nominally significant on the historical 223 Ma tree to unsupported on the Grafen primary tree, and BM/OU/EB rankings likewise depended on branch-length representation. Shape-axis signal was more stable than geometry signal. These checks cannot resolve uncertainty introduced by broader taxonomic proxy assignment, averaging multiple specimens within tips or uneven family sampling. Phylogenetic geometry results are therefore treated as exploratory trends rather than positive tests. Associated robustness summaries include PGLS, phylogenetic signal, evolutionary-model rankings and continuous root-state estimates (Supplementary Figs. 16-19; Supplementary Tables 27-34).

One point that deserves brief discussion is the placement of *Cimberis* in our dataset. Because our comparative framework follows the backbone of McKenna et al. [9], we treated *Cimberis* as part of Nemonychidae. More recent phylogenomic work, however, recovers Cimberididae as a distinct monophyletic family [48]. In our morphospace, *Cimberis* also shows a slight offset from the remaining nemonychid sample, which may indicate that this revised family-level placement is reflected in trochanter morphology. Given the limited sampling, this observation requires confirmation with broader representation of both Cimberididae and Nemonychidae.

### Evolutionary and ecological implications

The continuous shape variation and restricted set of realized character combinations suggest that joint diversification involved differential modification of interdependent elements rather than repeated shifts between sharply discrete architectures. Comparable patterns occur in other complex biomechanical systems, where integration, mechanical sensitivity and functional dependencies can channel evolutionary variation [3,4,36–38,49,50]. In the main dataset, winding angle covaried with the major shape axes (PC1-PC2), axial span and size. Axial span, fitted pitch and angular extent describe related aspects of the same traced geometry, but axial span was obtained directly from the trajectory endpoints rather than calculated from pitch and winding angle. These statistical associations do not by themselves demonstrate mechanical integration; direct tests require kinematics, loading experiments or simulations.

Family-level disparity provides tentative ecological context for this variation. Anthribidae showed the highest estimated disparity and also span an unusually broad ecological repertoire, including predation and associations with dead-wood, fungal, seed, fruit, pollen and other plant tissues [51–55], with movement, feeding, hunting and oviposition across these substrates imposing varied postural and stabilizing demands on the middle legs that may have contributed to mesocoxal diversification. Brentidae also showed high disparity, but specimen-level scores indicated that this primarily reflected a pronounced separation in morphospace between centrally positioned *Brentus* and peripheral Apioninae, rather than uniformly high variation throughout the family (Supplementary Table 2). This separation parallels phylogenetic and systematic evidence that Apioninae represent a distinct lineage within the broader brentid assemblage [56–58]. Attelabidae combined comparatively low trochanteral disparity with a highly coordinated maternal plant-manipulation system. Females weaken leaf tissue with the rostrum and mouthparts and subsequently use their legs to pinch, fold, roll and tighten the leaf into a brood structure [59]. Detailed observations of *Euops chinensis* further show a serrated foreleg assisting leaf cutting and the fore- and middle legs participating in the manipulation sequence [60]. These behaviours repeatedly require the legs to grip the leaf, brace the body and stabilize the developing roll. The comparatively narrow trochanteral morphospace of the sampled Attelabidae may therefore reflect conservation around a mechanically effective mesocoxal configuration associated with this highly coordinated behaviour. However, available behavioural studies rarely distinguish the precise contribution of each leg pair during rolling, leaving this proposed functional connection to be tested directly. The single belid specimen precludes a comparable family-level interpretation.

These family-level correspondences motivated an exploratory proxy-tip test of broad ecological associations. The two gymnosperm-coded proxy tips had smaller, more compact trochanters than the ten angiosperm-coded proxy tips, with the contrast expressed primarily along PC2. PC2 differed in PGLS after multiple-testing correction (FDR-adjusted P = 0.014), but the corresponding phylogenetic analysis of variance (ANOVA) did not survive correction (raw P = 0.006, FDR-adjusted P = 0.108; Supplementary Fig. 20; Supplementary Tables 35 and 36). Centroid size likewise differed in PGLS (FDR-adjusted P = 0.014) but not in the corrected phylogenetic ANOVA (Supplementary Fig. 21; Supplementary Tables 35 and 36). The elongation represented by PC2 occurs primarily at the trochanteral tip projecting into the coxa and could plausibly affect coxal guidance, but engagement and stability were not measured. Because the comparison contains only two gymnosperm-coded proxies, largely represents earlier-diverging groups, and aggregates heterogeneous sampled genera into broader tips, ecology, phylogenetic history and proxy assignment cannot be separated. The observed contrast is therefore hypothesis-generating rather than evidence for a general host-lineage effect.

Other ecological predictors showed weak or inconsistent associations after correction (Supplementary Figs. 22-24; Supplementary Tables 37-38). Winding angle was associated with fungal coding in the main-dataset PGLS (9 versus 2 proxy tips; FDR-adjusted P = 0.037), but this contrast was unsupported by phylogenetic ANOVA. The severe imbalance and method dependence therefore preclude a general fungal-association inference. Size-associated shape variation was similarly limited, as centroid size explained 7.0% of the complete atlas-derived shape space at specimen level, although PC1-PC5 captured 92.6% of the fitted allometric deformation and the clearest axis-specific effect remained concentrated on PC2 (Extended Data Fig. 3; Supplementary Tables 18-22). Ecology and size were therefore associated with particular dimensions without explaining the overall organization of trochanteral morphospace or screw joint geometry. This selective and analysis-sensitive ecological signal is compatible with the broader importance of host associations and life-history shifts in insect diversification generally [61] and in weevils specifically [42,62,63], but should be treated as hypothesis-generating.

These exploratory ecological contrasts motivate, but do not test, a functional hypothesis linking joint architecture to oviposition and substrate-manipulation behaviour. Females that excavate oviposition sites with the rostrum must brace the body against resistant plant tissue, transmitting reaction forces through the legs. A true screw-and-nut joint could help maintain controlled engagement between the trochanter and coxa and resist unwanted lateral displacement under these loads, consistent with the contribution of screw-and-nut joints to stable leg postures in *Trigonopterus* [24,64]. In our sample, Caridae, Brentidae and Curculionidae, in which rostral excavation of oviposition sites is widespread, predominantly possessed true screw-and-nut joints. Carids drill into closed female cones, whereas angiosperm-associated brentids and curculionids excavate oviposition sites in diverse living tissues [25,42,62].

Other lineages provide contrasting mechanical contexts. Nemonychids deposit eggs among exposed conifer microsporangia, while most anthribids develop in fungus-colonized dead wood and use an abdominal ovipositor rather than the rostrum to insert their eggs into the substrate [25]. *Anthribus nebulosus* deposits eggs within scale-insect brood chambers [51], and *Urodon rufipes* develops in seeds of *Reseda* [65]. These sampled lineages generally lacked complete screw-and-nut joints. The sampled attelabids retained a trochanteral screw component but did not consistently possess a complementary coxal thread. Their cutting, folding and rolling of plant tissues relies extensively on the legs [59], suggesting that the trochanteral screw may support stability during repeated substrate manipulation rather than rostral drilling. This hypothesis predicts that independent transitions towards mechanically demanding drilling or leg-mediated plant manipulation should be accompanied by convergent changes in joint phenotype, a relationship that can be tested through denser within-lineage sampling and direct measurements of joint kinematics and loading.

## Conclusion

Our results recast the weevil coxa-trochanteral screw joint from a singular mechanical curiosity as a diversified biomechanical system. Across the sampled Curculionoidea, trochanter shape varies continuously and overlaps among joint types, whereas the articulation occupies only a restricted subset of possible structural combinations. Coxal winding was never observed without a corresponding trochanteral helix. Robust helix fitting showed substantial geometric heterogeneity: winding angle covaried with shape and size in the main dataset, whereas joint-type comparisons were limited by severe group imbalance and proxy-level relationships varied across tree representations. Fitted axial pitch showed no robust association with shape, size or proxy phylogeny. The true screw-and-nut configuration in Caridae shows that this architecture is not confined to Brentidae and Curculionidae in our dataset, but sparse taxon sampling and the proxy phylogeny do not establish its time or direction of origin. Direct biomechanical experiments, kinematic analyses and repeated tracing will be needed to connect structural diversity with movement, loading and performance.

By integrating synchrotron micro-CT, AI-assisted segmentation, landmark-free atlas-based morphometrics and phylogenetic comparative analyses, we resolved a concealed internal articulation and quantified an evolutionary phenotype. More broadly, our approach shows how concealed internal joint systems in small-bodied organisms can be brought into a comparative evolutionary framework, providing a basis for testing whether other hidden biomechanical traits show comparable patterns of diversification.

## Materials and Methods

### Taxon sampling

We analysed the coxa-trochanteral joint across a broad sample of Curculionoidea including both early-diverging and derived lineages. We selected specimens from lineages with documented or suspected screw-like articulations, as well as those without such reports. The dataset includes representatives of Nemonychidae (*Basilogeus dacrycarpi*, *Cimberis attelaboides*), Anthribidae (*Anthribus nebulosus*, *Euparius paganus*, *Ormiscus saltator*, *Urodon rufipes*), Belidae (*Proterhinus sp*.), Attelabidae (*Coenorhinus pauxillus*, *Merhynchites sp*., *Paramechoris hirtus*, *Rhynchites cupreus*), Caridae (*Car pini*), Brentidae (*Apion frumentarium*, *Apion haematodes*, *Aspidapion validum*, *Brentus* sp., *Nanophyes marmoratus*, *Oxystoma* sp., *Perapion curtirostre*, *Protapion* sp.) and Curculionidae (**Baridinae:** *Limnobaris* sp.; **Ceutorhynchinae:** *Coeliodes cinctus, Mononychus punctumalbum, Nedyus quadrimaculatus, Parethelcus pollinarius*; **Conoderinae:** *Phaenomerus foveipennis, Zygops sp.*; **Cossoninae**: *Brachytemnus porcatus, Scobinoides dentatus*; **Cryptorhynchinae:** *Acalles fallax, Echinodera micros, Echinodera hypocrita, Trigonopterus plicicollis, Trigonopterus pseudonasutus*; **Curculioninae:** *Archarius pyrrhoceras, Cionus tuberculosus, Cleopus solani, Curculio glandium, Dorytomus taeniatus, Orchestes testaceus, Rhinusa tetra, Sibinia pellucens*; **Dryophthorinae:** *Dryophthorus corticalis;* **Entiminae:** *Barypeithes araneiformis, Brachysomus hirtus, Metapocyrtus adspersus, Otiorhynchus singularis, Phyllobius* sp.*, Polydrusus marginatus, Sitona* sp.*, Sitona hispidulus, Strophosoma capitatum, Strophosoma melanogrammum;* **Cyclominae:** *Listronotus* sp.; **Erirhininae:** *Lissorhoptrus oryzophilus, Neochetina* sp.; **Hyperinae:** *Hypera postica*; **Lixinae:** *Larinus turbinatus*; **Mesoptiliinae:** *Magdalis* sp.; **Molytinae:** *Adexius scrobipennis, Leiosoma deflexum*; **Platypodinae:** *Euplatypus parallelus*; **Scolytinae:** *Hylesinus crenatus, Hylurgops palliatus, Leperisinus fraxini, Scolytus carpini, Xylocleptes bispinus, Xylosandrus germanus*). Specimen identifiers, taxonomic assignments and tree-tip assignments are listed in Supplementary Table 1.

For phylogenetic analyses, the 68 specimens were assigned to 15 taxonomic proxy tips on the available backbone (Supplementary Table 1). Five assignments were exact sampled-genus matches; 63 used a broader taxonomic proxy. Multiple specimens assigned to the same proxy were summarized at tip level. These analyses therefore characterize the chosen proxy-tip dataset and are not genus-level comparative analyses of all sampled taxa.

### Data acquisition, segmentation and mesh processing

The specimens were scanned at the UFO-I high-throughput setup at the IMAGE beamline [66] at the KIT Light Source. A polychromatic X-ray beam, produced by a superconducting wiggler (corresponding to a magnetic field B = 1 T), was filtered with 10 mm pyrolytic graphite sheets, resulting in a spectrum peaking at approximately 16.5 keV with a full width at half maximum of approximately 11 keV. The setup included a rapid indirect detector system with a scintillator, visible-light optics, a white-beam microscope (Optique Peter, Lentilly, France) [67] and a 12-bit pco.dimax high-speed camera (Excelitas PCO GmbH, Kelheim, Germany) with 2016 × 2016 pixels, each 11 µm in size. A nominal 10× optical setting yielded a calibrated effective pixel size of 1.22 µm at the specimen plane. Each scan consisted of 100 dark-field images, 200 flat-field images and 3000 equiangularly distributed radiographic projections over 180° at 60 fps. The control system “concert” [68] facilitated automated data acquisition. The UFO framework [69] was employed for data processing, including dark- and flat-field correction and phase retrieval. Final three-dimensional tomographic reconstruction was performed with tofu [70] and included ring removal, 8-bit conversion and blending of the phase and absorption reconstructions.

Internal joint morphology was documented using synchrotron X-ray microtomography. Tomographic TIFF stacks were imported into Amira 2022.2 (Thermo Fisher Scientific, Waltham, MA, USA) [71] and cropped to isolate a single coxa-trochanteral joint per specimen. Cropping isolated the focal joint from the surrounding tomogram but retained the complete coxa in every specimen; no coxa mask was intersected by the boundary of the original image stack. Relevant structures were segmented in napari 0.6.1 [72] using the nnInteractive plugin [31], and resulting binary segmentations were returned to Amira, where surface meshes were generated and exported as OBJ files.

Meshes were batch-processed in Cinema 4D 2026 (Maxon Computer GmbH, Bad Homburg, Germany) using a study-specific Python script prior to morphometric analysis. This script is retained in the project workspace but is not included in the public code repository. Because coxal morphology is strongly influenced by internal muscle volume and other factors, only trochanteral meshes were used for landmark-free shape analysis. Where necessary, meshes were mirrored along the global X-axis to ensure consistent orientation across specimens. Each mesh was centred at the world origin, quad-remeshed to an absolute target of 10,000 polygons with automatic hard-edge handling and adaptive remeshing disabled, and then triangulated. Meshes were optimized with a tolerance of 0.01 and smoothed using three iterations of uniform Laplacian smoothing (α = 0.40). The automated cleanup retained the largest connected component, removed degenerate and duplicate triangles, performed a final zero-tolerance optimization and corrected outward normal orientation before OBJ and PLY export. Hole closing was disabled for the production batch. The processed coxa and trochanter surface meshes are deposited separately on RADAR4KIT rather than in GitHub; the original Checkpoint landmark files, aligned VTK meshes and initial atlas template are not redistributed. Prior to atlas construction, all trochanter meshes were aligned using generalized Procrustes analysis (GPA) based on three homologous landmarks placed in Checkpoint 25.06.02 (Stratovan Corporation, Davis, CA, USA): one landmark at the tip of the proximal trochanteral process projecting into the coxa, and two landmarks at opposing extremities of the distal articulation with the femur. Together, these landmarks formed a triangular configuration defining the orientation and scale of each specimen. Landmark configurations were centred, scaled to unit centroid size and rotated into a common canonical frame, and the resulting transformations were applied to the corresponding surface meshes. These landmarks were used only for alignment and subsequent shape variation was quantified from the complete trochanteral surfaces.

### Landmark-free quantification of trochanter shape

Trochanteral shape was quantified using a landmark-free morphometric framework implemented in Deformetrica v4.3.0 [34]. Processed trochanter meshes were used for deterministic atlas construction under a large deformation diffeomorphic metric mapping (LDDMM) framework, following the general Deformetrica workflow adapted to the present dataset. Atlas construction was performed in three dimensions using SurfaceMesh objects. During workflow development, kernel width 0.1 was first selected by visual comparison of atlas reconstructions across widths from 0.025 to 0.2, using the 67 specimens available at that stage (Supplementary Fig. 25). We subsequently audited the archived development-stage runs quantitatively. Because the meshes were GPA-aligned and scale-standardized, kernel widths are expressed in normalized coordinate units. Pairwise specimen-distance structure placed width 0.1 within a stable intermediate region: full-space Spearman’s ρ was 0.986 relative to width 0.075 and 0.973 relative to width 0.15, and the corresponding PC1-PC5 Procrustes similarities were 0.982 and 0.958. By contrast, width 0.025 showed substantially lower concordance with width 0.1 (ρ = 0.355), indicating greater sensitivity of the ordination at the narrowest tested scale. Among the concordant intermediate settings, width 0.1 retained 900 control points, compared with 294 at width 0.15 and 125 at width 0.2. The retrospective audit supported 0.1 as a central intermediate setting that preserved relatively fine deformation support while avoiding the morphospace sensitivity observed at 0.025; it did not identify a unique quantitative optimum. The estimated atlas template had bounding-box dimensions of 0.909 × 0.946 × 0.799 and an enclosed volume of 0.0856 normalized units³. The selected kernel width corresponded to approximately 10.6% of the maximum template span and 6.5% of its bounding-box diagonal. Width 0.1 was then fixed without further tuning for the final atlas, which comprised 68 trochanter meshes after the addition of *Rhynchites cupreus*. In the final atlas run, both the surface-attachment and deformation kernels used a KeOps kernel width of 0.1. The surface-attachment noise standard deviation was set to 0.05, corresponding to approximately 5.3% of the maximum template span and allowing modest residual surface discrepancies after alignment. Deformation trajectories were discretized into 30 time points to provide a sufficiently fine numerical representation of the diffeomorphic flow while maintaining computational tractability. Optimization was performed using gradient ascent with a conservative initial step size of 0.01 and adaptive line search. A maximum of 1,000 iterations was specified as an upper limit, as recorded in the released optimization configuration; neither a convergence tolerance nor the completed iteration count is available in the release-verifiable records. Atlas construction yielded an estimated mean shape together with subject-specific momenta and control points describing major regions of shape variability.

Atlas-derived shape variation was summarized by ordinary principal component analysis (PCA) of the vectorized subject-specific momenta. For each specimen, momenta across all control points and spatial dimensions were concatenated into a single feature vector, centred across specimens and analysed without feature scaling. This provided a linear tangent-space representation of the LDDMM deformation parameters. Because nonlinear kernel embeddings are also used for momenta-based morphometric data, we repeated the ordination using radial-basis-function kernel PCA as a sensitivity analysis. Both the parameter used in the original workflow (γ = 0.25) and a data-adaptive median-distance kernel preserved the broad morphospace structure and the principal family, joint-type and allometric results (Supplementary Fig. 2; Supplementary Table 4).

PC1-PC5 were retained because each explained at least approximately 5% of total shape variation. Together, they accounted for 61.4%. This per-axis threshold retained the dominant anatomically interpretable modes while excluding higher-order components with individually small contributions. PC1-PC5 were used for disparity, clusterability and clustering, phylogenetic signal, ancestral-state reconstruction and evolutionary model fitting. Ecological analyses were restricted to PC1 and PC2, the axes used for the reported ecological interpretation. The primary multivariate allometry analysis used the complete ordinary-PCA representation of all 68 specimens, comprising all 67 non-zero PCs and therefore 100% of the atlas-derived shape variance. PC1-PC5 were retained for axis-specific interpretation and sensitivity analyses.

### Quantification of screw joint geometry

Geometric descriptors of the coxa-trochanteral interface were derived from manually placed ordered three-dimensional semilandmarks tracing the centre of the screw-like surface relief (Supplementary Fig. 26). Semilandmarks were placed only for the 64 specimens with discernible helical surface relief; the four specimens lacking such a ridge had no trajectory to measure. Coordinates were exported as CSV files. The released Cinema 4D/Python axis-and-circle implementation was first reproduced outside Cinema 4D as a numerical audit. Primary measurements were then obtained with a standalone Python/SciPy robust circular three-dimensional helix fit. The model continuously optimized axis orientation and position, radius, angular phase and axial rise from all trajectory points, jointly minimizing radial and axial residuals with a soft-L1 loss and multiple starting solutions. Fit diagnostics included radial, axial and three-dimensional helix RMS, axial-angular linearity, backtracking and near-optimal-solution stability. The scale-free relative-RMS threshold defining the main dataset was fixed before downstream biological associations were inspected. Other diagnostic warnings were retained for transparent quality control but were not used as exclusion criteria, because point count and angular coverage partly reflect the biological extent of the helix. Semilandmark placement followed a standardized single-observer protocol and was performed once per specimen; placement repeatability was not quantified. Both implementations are released. The original ordered per-specimen semilandmark CSV files are not included in the public repository; their observed point names and XYZ coordinates are retained in the released point-residual table so that the trajectories can be reconstructed without reproducing the original input-file packaging.

Signed and absolute winding angles were calculated from the unwrapped angular coordinates of the fitted helix, with turn number defined as winding angle divided by 360. Fitted axial pitch was calculated as 2π times the absolute axial rise per radian. Axial span was defined as the absolute displacement between the first and last traced semilandmarks projected onto the fitted helix axis. The model-derived axial rise over the traced angular extent, calculated as fitted pitch multiplied by absolute winding angle divided by 360°, was retained separately as an audit variable but was not used in downstream analyses. Start-to-end distance was likewise retained as an audit variable. Linear descriptors are reported in the unit-centroid-size coordinate system and winding angle in degrees. All 64 traced trajectories converged. The main dataset required helix RMS/fitted radius ≤ 0.10, limiting the average three-dimensional residual to no more than 10% of the fitted helix radius. Dryophthorus corticalis (helix RMS/fitted radius = 0.180) exceeded this threshold, leaving 63 trajectories in the main dataset. Additional diagnostic flags were recorded for fewer than 20 trajectory points (with fewer than 10 flagged separately), angular coverage below 180° (with below 30° flagged separately), fitted radius divided by start-to-end trajectory span above 5, axial-angular R² below 0.80, angular backtracking fraction above 0.10, a near-optimal winding-angle range above 20°, or a near-optimal axis separation above 15°. These flags were not used to exclude trajectories: in particular, low point counts and limited angular coverage can result from biologically short helices. For quality control, 200 moving-block residual-bootstrap fits were generated for each of all 64 traced trajectories, yielding 12,800 successful draws. Downstream uncertainty propagation for the main dataset used the corresponding 12,600 draws from its 63 retained trajectories. This bootstrap does not include manual semilandmark-placement repeatability. Fit and trajectory diagnostics are shown in Supplementary Fig. 27.

### Ecological data

Ecological information was compiled from published literature for the genera represented in the comparative dataset and coded at the tree-tip level rather than for individual imaged specimens. Where ecological information within a lineage was heterogeneous or incomplete, coding was based on the best-supported broad ecological state represented by the sampled lineage, and ambiguous cases were treated conservatively.

To avoid overparameterization, detailed ecological observations were collapsed into four broad predictors. Host lineage was coded as angiosperm-associated, gymnosperm-associated or mixed, whereas wood association, larval lifestyle and fungal association were coded as binary variables. Mixed lineages were retained in descriptive summaries but excluded from pairwise tests of angiosperm-versus gymnosperm-associated taxa. These codings should be understood as broad comparative categories rather than fine-scale ecological reconstructions. The resulting ecology matrix is provided in Supplementary Table 39.

### Phylogenetic framework and tip assignment

The phylogenetic framework was based on the Curculionoidea amino-acid supermatrix of McKenna et al. [9]. Fourteen retained sequences comprising 1,909,716 aligned amino-acid sites were analysed with IQ-TREE multicore 2.4.0 [73] using the source partition file, ModelFinder with partition merging (MFP+MERGE), 1,000 ultrafast-bootstrap replicates, 1,000 SH-aLRT replicates and automatic thread selection. The recorded random seed was 783104. The resulting inferred tree and downstream working variants are archived in the GitHub repository; the source alignment, partition files and source-study IQ-TREE command, log and report are not redistributed and must be obtained or recreated from McKenna et al. [9]. The resulting maximum-likelihood tree was rooted on Nemonychidae, represented by Rhynchitomacerinus, following published Curculionoidea phylogenies that consistently recover nemonychids among the earliest-diverging weevil lineages [9,42–44].

Because Caridae was not represented in the published molecular alignment, it could not be placed directly by phylogenetic inference. To include this lineage, Caridae was inserted manually in the position supported by current phylogenetic literature, as sister to the clade comprising Brentidae and Curculionidae. In practice, Caridae was inserted as a new tip at the most recent common ancestor of Brentidae and Curculionidae using a minimal positive branch length, which was subsequently rescaled during time calibration.

After insertion of Caridae, the primary working phylogeny was assigned topology-based Grafen branch lengths using ape::compute.brlen [74]. These branch lengths encode relative node depth for comparative analysis and are not interpreted as geological time; this study does not estimate divergence times. Fossil-calibrated chronos trees were retained only as sensitivity representations. Fossil minima followed McKenna et al. [9] supplementary table S5: 157.3 Ma for crown Curculionoidea based on Nemonychidae fossils and 113.0 Ma for crown Curculionidae based on Arariperhinus monnei from the Crato Formation. The 223 Ma ceiling in the archived historical tree did not originate from McKenna et al.; it followed Letsch et al. [75], who used the age of Leehermania prorova, then interpreted as the oldest polyphagan beetle. Because Leehermania was subsequently reassigned to Myxophaga [76], we retained that tree only as a historical sensitivity case. Additional calibrated sensitivity trees used a fixed 195 Ma root, following the lower margin of the estimated origin of Phytophaga used as a hard maximum in recent Belidae dating [77], or root intervals of 157.3-170, 157.3-195 and 157.3-223 Ma, with Curculionidae constrained to 113-151 Ma where applicable. Calibrated solutions depended on the chosen constraints and optimizer start values, so none was selected as the primary phylogeny (Supplementary Data 3-4).

We used the McKenna et al. [9] Curculionoidea phylogeny as a taxonomic backbone rather than as an exact tree of the imaged taxa. Because most sampled genera were absent, each of the 68 specimens was assigned to the closest defensible available taxonomic proxy tip based on taxonomic affinity (Supplementary Table 1). This yielded 15 proxy tips: five specimen assignments were exact sampled-genus matches and 63 were broader taxonomic proxies. Tree labels were standardized and assignments were checked manually. Where several specimens mapped to one proxy, continuous measurements were averaged; discrete states were retained only when contributors agreed and otherwise treated as ambiguous or missing. The complete mapping, including within-tip sample sizes, is provided in taxonomic_proxy_mapping.csv. Comparative results are conditional on these assignments and are treated as exploratory.

### Morphospace structure, clustering and disparity

Morphospace structure was assessed from ordinary-PCA scores derived from vectorized momenta. Specimens were visualized primarily in PC1-PC2 and PC2-PC3 space, and variance explained by each component was calculated from the PCA eigenvalues. PC1-PC5 were retained as the main representation of the atlas-derived shape space because each explained at least approximately 5% of total variation. The same family, joint-type and allometric comparisons were repeated on matched RBF-kPCA axes as an ordination-sensitivity analysis (Supplementary Fig. 2; Supplementary Table 4). The kPCA family-dispersion calculation was a parallel exploratory diagnostic, not an exact rerun of the primary bias-adjusted four-family PERMDISP.

Before assigning specimens to discrete groups, global clusterability was assessed descriptively from standardized PC1-PC5 scores using 100 randomized Hopkins evaluations, each sampling six observed points and six uniformly generated reference points (n = 68, m = 6, seed = 1; Supplementary Fig. 3; Supplementary Table 5). Because the sample did not permit conventional large-sample conditions for formal Hopkins inference, the result was interpreted only as an exploratory indication of departure from spatial randomness. To test whether this qualitative indication depended on m = 6, we additionally performed 1,000 randomized evaluations at each m from 4 to 10 using documented independent seeds. Mean H ranged from 0.885 to 0.931 and 85.2-93.8% of evaluations exceeded 0.75, so the qualitative indication was not unique to m = 6; these repetitions remained descriptive and did not add independent specimens. Complementary clustering approaches were then applied to the same standardized PC1-PC5 scores: k-means, hierarchical clustering using Ward.D2, and model-based clustering implemented in mclust [78]. For k-means and hierarchical clustering, candidate values of k = 2-10 were evaluated using average silhouette width, and gap statistics were additionally calculated for k-means (Supplementary Fig. 4). For model-based clustering, the number of groups and covariance structure were selected using Bayesian information criterion.

Family-level disparity was summarized descriptively as the mean squared Euclidean distance of specimens from their family centroid in PC1-PC5 space [79,80]. Only families represented by at least three specimens were included, and uncertainty in these squared-distance summaries was quantified with 5,000 bootstrap resamples. These summaries and their rarefied counterparts were not the responses used for PERMDISP. Global differences in multivariate dispersion were tested separately on the same PC1-PC5 data using bias-adjusted unsquared Euclidean distances to family centroids and 9,999 PERMDISP permutations. Pairwise PERMDISP comparisons were adjusted with the Benjamini-Hochberg procedure. To evaluate sensitivity of the descriptive disparity ranking to unequal sample sizes, the mean-squared-distance calculation was additionally rarefied to four specimens per family across 5,000 random subsamples.

### Shape-geometry relationships, allometry and coxal wall thickness

Relationships between trochanter shape and screw joint geometry were assessed by exact specimen-identifier matching of principal-component scores and robust helix fits. The main analysis used 63 trajectories with helix RMS divided by fitted radius ≤ 0.10. Linear models related absolute winding angle, axial span and fitted axial pitch to PC1 and PC2, with PC1-PC5 and regional models reported as supplementary analyses. The complete models were propagated through all 200 conditional-bootstrap draws per specimen, so point estimates from the main dataset were evaluated together with their conditional measurement-sensitivity distributions.

Allometric effects were assessed by relating centroid size to trochanter shape and robust screw joint geometry at specimen level. Centroid size was calculated from the raw, unscaled Checkpoint landmark coordinates before GPA centring and scaling, as the square root of the summed squared distances of the three landmarks from their centroid, and was log-transformed. Because atlas shapes were normalized for scale before PCA, the isometric null predicts no systematic multivariate relationship between shape and log centroid size. We tested this null primarily with RRPP::lm.rrpp [81] and, as a complementary implementation, geomorph::procD.lm [82], using all 67 non-zero atlas PCs and 9,999 residual permutations. A PC1-PC5 regression was retained as a sensitivity analysis. Axis-specific models were fitted for all non-zero PCs with Holm correction, and each PC’s contribution to the fitted allometric deformation was quantified from the multivariate regression vector. Geometry-size models were fitted to the 63-specimen main dataset; response variables were absolute winding angle, axial span and fitted axial pitch. Shape-only phylogenetic allometry used all 15 taxonomic proxy tips, whereas geometry PGLS models used the 14 proxy tips obtained by matching the 63 main-dataset specimens before aggregation. For both datasets, centroid size was averaged within each proxy tip and the resulting mean was natural-log transformed; Pagel’s λ was estimated by maximum likelihood within [0, 1] without a fixed-λ fallback. A one-dimensional allometric projection score was calculated only for visualization in Extended Data Fig. 3 and not used for inference.

Coxal wall thickness was quantified from the complete three-dimensional binary coxa masks. Each stack was cropped to the foreground extent, and the largest six-connected component was retained to remove isolated segmentation fragments. Local thickness was calculated at every coxa voxel as the diameter of the largest sphere fully contained within the binary mask and containing that voxel. The median of the resulting voxel-wise distribution was used as the primary whole-volume wall-thickness estimate for each specimen, with the 10th percentile evaluated as a lower-tail sensitivity metric. To reduce memory requirements, masks were analysed at 0.75 isotropic scale. In a five-specimen validation subset spanning lower, middle and upper portions of the observed coxa-size and wall-thickness distributions, 0.75-scale median estimates differed from full-resolution calculations by 0.46-5.21%; the specimen-level comparison is archived with Supplementary Table 9 source data. Thickness was converted to micrometres using the 1.22-µm voxel size. Coxa size was estimated as the diagonal of the three-dimensional OBJ bounding box. Associations were tested on log10-transformed size and thickness, including coxal-wall-opening state and its interaction with size. The whole-volume thickness dataset and associated model summaries are provided in Supplementary Tables 9-11.

### Joint typology and ecology

To describe recurring structural configurations of the coxa-trochanteral articulation, specimens were coded for four discrete morphological features: a helical ridge on the trochanter, a corresponding guiding track in the coxa, a stabilizing coxal socket and a coxal wall opening. Their combinations yielded four recurrent structural categories: true screw-and-nut joints, unopposed screw configurations, socket-guided rotational joints and unguided rotational joints. The wall opening was retained as an accessory character rather than defining an additional joint type. Joint assignments were merged with shape and robust geometry data. In the main dataset, Kruskal-Wallis tests compared absolute winding angle, fitted axial pitch and axial span between 57 true screw-and-nut joints and only three unopposed screw configurations. Their three P values formed one correction family and were adjusted together using the Benjamini-Hochberg procedure. A fixed-seed Euclidean PERMANOVA (999 permutations) tested combined standardized geometry, with betadisper used separately to assess dispersion; these multivariate diagnostics were not included in the three-test univariate correction family. All joint-type comparisons were treated as exploratory because of the severe imbalance.

Using the ecology matrix described above, ecological effects on shape were analysed for PC1 and PC2 at the 15 taxonomic proxy tips. Geometry-specific ecological models used the 14 matched proxy tips from the main dataset. Descriptive summaries and non-phylogenetic Wilcoxon tests were accompanied by phylogenetic ANOVA and PGLS. Within each analytical framework, P values for all tested trait-predictor combinations were adjusted together using the Benjamini-Hochberg procedure [83]. One geometry-ecology association survived this correction in PGLS (absolute winding angle versus fungal association; nine non-associated versus two associated proxy tips; FDR-adjusted P = 0.037), but not in the companion phylogenetic ANOVA (raw and FDR-adjusted P = 0.853). Because this contrast was strongly imbalanced and method-dependent, it was treated as exploratory rather than general evidence of fungal-associated geometry. Host-lineage tests for geometry were not estimable because only one gymnosperm-coded tip retained geometry data. These analyses cannot separate ecology from phylogeny, proxy assignment or within-tip heterogeneity.

### Phylogenetic comparative analyses

Phylogenetic comparative analyses were used to describe trait patterns conditional on the selected taxonomic proxy tree [84,85]. Shape-only analyses used all 15 proxy tips. Geometry analyses matched specimens exactly to the 63-trajectory main dataset before aggregation, yielding 14 proxy tips. The nemonychid proxy tip was absent from the geometry dataset because neither sampled nemonychid had a measurable helical ridge. Continuous variables were averaged across contributing specimens; discrete variables were retained only when contributors agreed and otherwise treated as ambiguous or missing. These proxy-level analyses should not be interpreted as exact genus-level comparative tests.

Phylogenetic signal was evaluated using Blomberg’s K and Pagel’s lambda. Within each analysis dataset, Benjamini-Hochberg false-discovery-rate correction was applied jointly to all trait-by-method P values from these two signal tests. Alternative univariate evolutionary models were compared by fitting Brownian motion, Ornstein-Uhlenbeck and early-burst models with geiger::fitContinuous [86] and comparing AICc support. Multivariate Brownian-motion, Ornstein-Uhlenbeck and early-burst shape models at 15 shape-only proxy tips were fitted with mvMORPH. Optimizer convergence and Hessian reliability were recorded separately; raw AIC values were retained for transparency, but ΔAIC rankings and Akaike weights were calculated only among fits with both successful optimizer convergence and a reliable Hessian diagnostic. The PC1-PC5 OU optimizer converged but its Hessian solution was unreliable, and the PC1-PC4 OU fit failed. Brownian-motion and early-burst fits were reliable for both shape representations. Continuous trait-trait and allometry PGLS analyses used nlme::gls [87] with a Pagel’s λ correlation structure implemented in ape::corPagel [88]. The categorical ecology PGLS analyses used caper::pgls, evaluating a maximum-likelihood λ fit together with fixed boundary candidates λ = 0 and λ = 1 and retaining the valid fit with the highest likelihood. Within each analysis dataset, continuous-model coefficient P values and ecology-factor P values were corrected in their respective predefined test families. Tree pruning, label standardization and specimen-to-tip aggregation preceded all geometry models. Phylogenetic ANOVA with 1,000 simulations was used for selected categorical contrasts where group sizes were adequate. Continuous ancestral-state reconstruction used phytools::fastAnc [89] under a Brownian-motion (BM) model for PC1-PC5 and centroid size on the 15-tip shape-only dataset, and for absolute winding angle and axial span on the 14-tip main geometry dataset. BM was specified as a common reconstruction model and was not selected from the exploratory BM/OU/EB model comparisons. Numerical node estimates, variances and 95% intervals were exported on the raw trait scale. Extended Data Fig. 5 maps used phytools::contMap under the same BM model with the same trait-specific tip sets; values were standardized within each trait only for the shared colour scale. All numerical estimates and maps are therefore BM-conditional. Coxal socket presence and coxal wall opening were both screened for stochastic character mapping with phytools::make.simmap under the equal-rates (ER) model. Coxal socket presence had only one state among non-missing proxy tips and was therefore not mapped; only the variable wall-opening character was reconstructed with 50 simulations. Geometry reconstructions were additionally interpreted as conditional summaries because of sparse proxy sampling, within-tip averaging and broader taxonomic assignments. Discrete ancestral-state summaries are provided in Supplementary Table 40. Sensitivity of comparative results to phylogenetic specification was assessed by repeating key analyses across 15 topology and branch-length representations: the Grafen primary working tree plus 14 alternatives, including trees with and without manually inserted Caridae, alternative Grafen transformations, unrooted and ultrametric representations, the historical 223 Ma calibration, a fixed 195 Ma calibration and three interval-calibrated trees (Supplementary Data 4). Phylogenetic signal, evolutionary-model, PGLS, phylogenetic-allometry, ecological and continuous ancestral-state analyses were repeated across this set. Selected PGLS models were also refitted after sequential removal of each proxy tip. These checks evaluate dependence on the available tree representation and calibration, but cannot resolve error introduced by broader taxonomic proxy assignment or sparse and uneven lineage sampling.

### Software and reproducibility

Image processing, segmentation, mesh preparation, morphometric analyses and statistical workflows used Amira 2022.2, napari 0.6.1 with nnInteractive, Checkpoint 25.06.02, Cinema 4D 2026, Deformetrica 4.3.0, Python and R. Release checks were performed with R 4.5.2 and Python 3.12.8; the current downstream audit and full-resolution coxa validation were repeated with Python 3.12.13, NumPy 2.3.5 and SciPy 1.18.0. R-package and downstream Python package lists are archived in the repository; only explicitly recorded component versions are treated as exact release metadata. The public scripts cover three-landmark GPA alignment, the legacy Cinema 4D/Python axis-and-circle workflow, robust three-dimensional helix fitting, uncertainty propagation, data integration, comparative analyses and figure generation; R visualizations used ggplot2 [90]. The parameters of the study-specific Cinema 4D batch remeshing, smoothing and mesh-cleanup step are summarized above, but Obj_Processing.py is not included in the public repository. The repository supports the downstream statistical analyses and derived tables and figures. It includes robust point estimates, point-level residuals containing the observed semilandmark XYZ coordinates, all 12,800 conditional-bootstrap draws, analysis-set definitions, audit data and processed outputs. It does not include the original per-specimen semilandmark CSV files, Checkpoint landmarks, aligned VTK meshes, initial atlas template, binary coxa masks, source phylogenetic alignment and partition files, or the source-study IQ-TREE command, log and report.

### Use of artificial intelligence tools

Generative artificial intelligence tools, including OpenAI ChatGPT/Codex, were used during manuscript preparation for editorial and programming support, including language polishing, troubleshooting analysis and visualization scripts, and organizing code and supplementary materials. All AI-assisted text, code and outputs were critically reviewed, edited and verified by the authors. No generative AI tool generated primary data, altered micro-CT data or segmentations, or drew scientific conclusions independently of the authors. nnInteractive was used separately as the interactive AI-assisted segmentation tool described above; its outputs were reviewed and corrected by the researchers.

All statistical analyses, morphometric measurements, figure outputs and scientific interpretations reported in the manuscript remain the responsibility of the authors.

## Data availability

The processed tomographic datasets and the surface meshes of coxae and trochanters are publicly accessible via RADAR4KIT under https://doi.org/10.35097/9p77hjk7wa656d6k. Derived analysis tables, supplementary source data, quality-control outputs and analysis-ready tabular inputs needed to rerun the downstream statistical analyses and scripted quantitative figures are archived in the public GitHub repository at https://github.com/heinjenny95/weevil-coxa-trochanter-joint-evolution. The repository is not a complete archive of all raw and intermediate inputs; the non-redistributed items are identified in the Software and reproducibility section.

## Code availability

Scripts for three-landmark generalized Procrustes alignment, robust three-dimensional helix fitting, downstream geometry analyses, morphometric analyses, phylogenetic comparative methods and publication-figure rendering are available in release v1.3.2 at https://github.com/heinjenny95/weevil-coxa-trochanter-joint-evolution/releases/tag/v1.3.2.

The study-specific Cinema 4D Obj_Processing.py script used for the earlier batch-remeshing step is not included.

## Supporting information

Interactive 3D Morphospace

Interactive 3D Heatmaps

Phylogenetic Tree

Tree Sensitivity analysis

## Acknowledgements

This work was supported by the Deutsche Forschungsgemeinschaft (DFG, German Research Foundation) - Project number 518779777. We thank the KIT Light Source for access to beamline facilities and the Institute for Beam Physics and Technology (IBPT) for operating the storage ring KARA. We thank Bruno Bell and Szatmári Ferenc for kindly providing photographs of live weevils used in Figure 1. Mention of trade names or commercial products in this publication is solely for the purpose of providing specific information and does not imply recommendation or endorsement by the USDA. The USDA is an equal opportunity employer and provider.

## Author contributions

T.v.d.K. conceived the study. J.H. and J.K. designed the analyses. J.H. processed the data, performed the morphometric, geometric and comparative analyses, prepared the figures and wrote the initial manuscript draft. T.v.d.K. supervised the project and contributed to study design, data interpretation and manuscript development. J.H., T.v.d.K., A.C., E.H. and M.Z. acquired the micro-CT data. A.R., O.B. and M.L.C. contributed specimens, taxonomic expertise and/or ecological information. J.K. and A.C.-F. contributed to the adaptation and implementation of the Deformetrica workflow. A.E., T.F., C.S., S.S., C.T. and N.Z. contributed to image reconstruction, segmentation workflows, data processing and computational infrastructure. All authors discussed the results, contributed to manuscript revision and approved the final version.

## Extended Data

**Extended Data Fig. 1.**
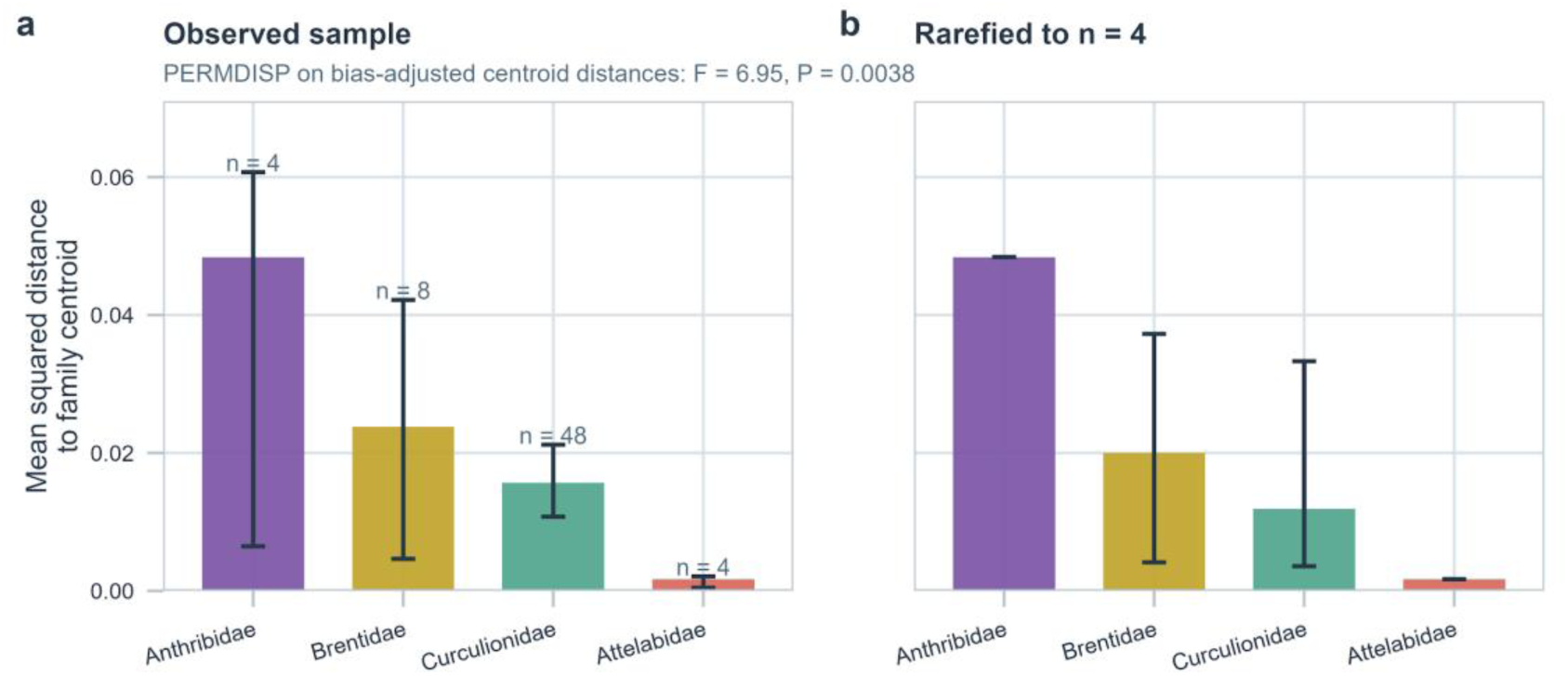
Within-family disparity in trochanteral morphospace. (a) Bars show observed family-level mean squared Euclidean distances of specimens from their family centroid in PC1-PC5 morphospace; error bars show bootstrap 95% confidence intervals and labels denote family sample size. These bars are descriptive. The F and P values above panel a come from a related but separate bias-adjusted PERMDISP on unsquared Euclidean distances to family centroids (F = 6.95, P = 0.0038). Pairwise PERMDISP indicated greater dispersion in Anthribidae than in Attelabidae (FDR-adjusted P = 0.0006) and Curculionidae (FDR-adjusted P = 0.0066); other pairwise contrasts were not significant after correction. (b) Rarefied mean-squared-distance estimates after subsampling each family to n = 4 specimens across 5,000 replicates. The preserved rank order indicates that the descriptive pattern was not solely driven by unequal family sample sizes.

**Extended Data Fig. 2.**
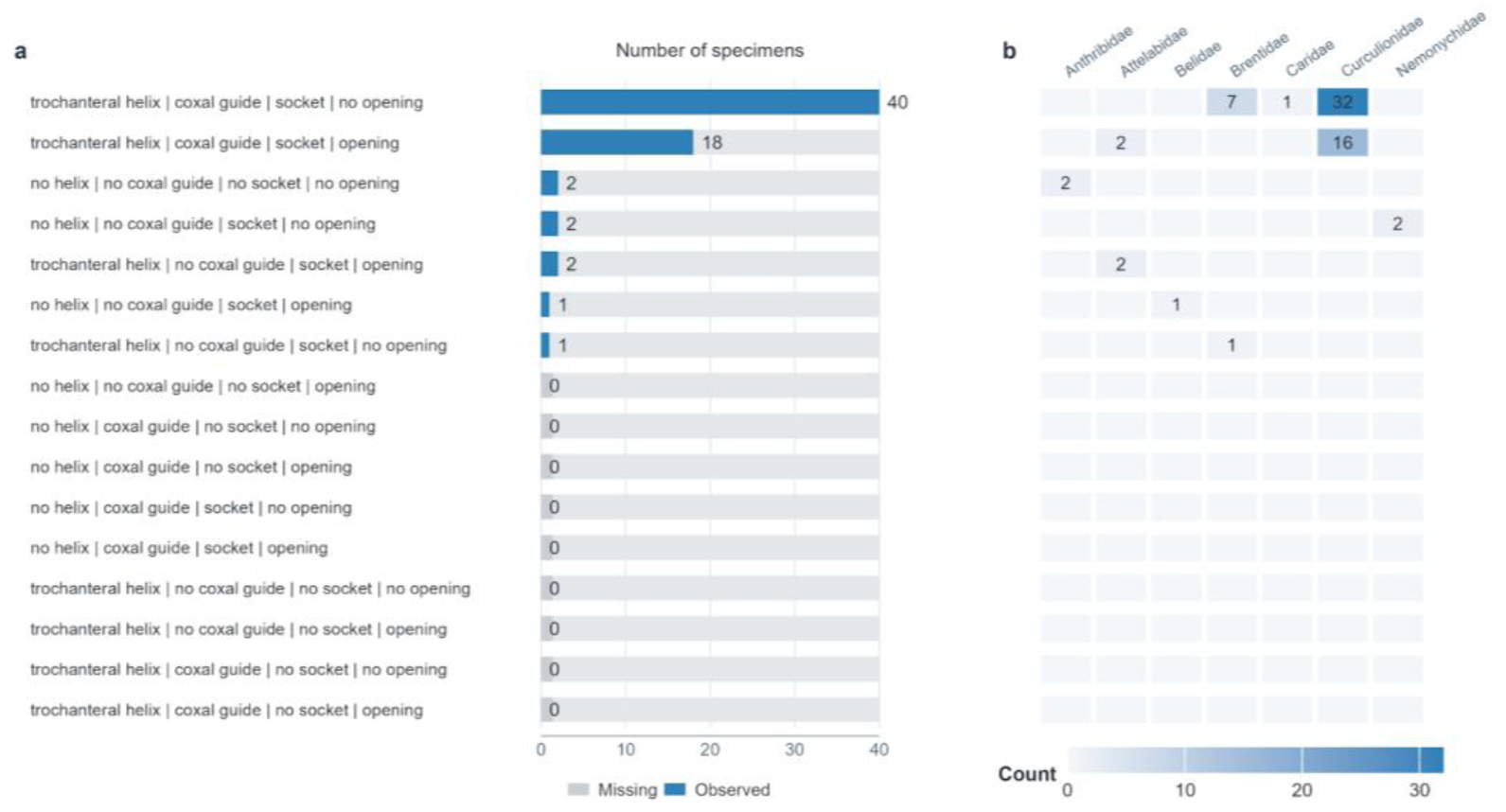
Observed and missing joint-character combinations across the sampled Curculionoidea. **(a)** Frequency of theoretically possible combinations of four discrete joint characters used in the typology analysis: presence or absence of a trochanteral helix, coxal guide, socket and coxal opening. Blue bars indicate combinations observed in the dataset, whereas grey bars indicate mechanically conceivable but unrealized combinations. **(b)** Family-level distribution of the observed combinations. Cell values indicate the number of specimens assigned to each family and joint-character combination. Together, the panels show that only a restricted subset of the theoretically possible design space is occupied, and that the dominant combination is broadly represented across sampled Curculionoidea rather than confined to a single family.

**Extended Data Fig. 3.**
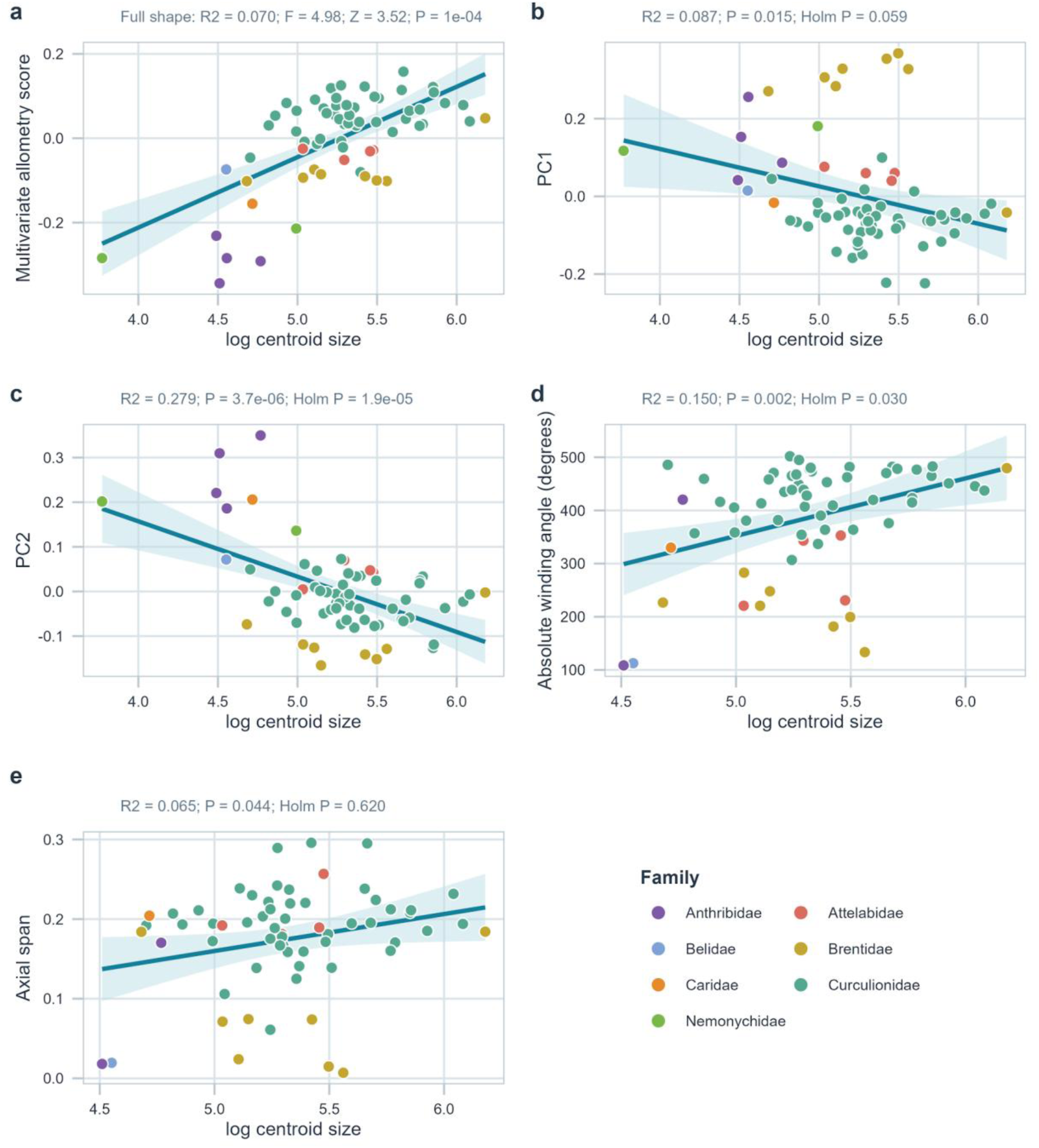
Multivariate and univariate allometry of trochanteral shape and robust screw joint geometry. (a) Full-shape multivariate allometry score versus log centroid size. The primary RRPP analysis used all 67 non-zero PCs and detected a small but significant effect (R² = 0.070, F = 4.98, Z = 3.52, P = 0.0001). (b,c) Specimen-level relationships of PC1 and PC2 with log centroid size; PC2 remained significant after Holm correction. (d,e) Absolute fitted winding angle and axial span versus log centroid size in the 63-specimen main dataset. Winding angle retained a Holm-corrected size association (R² = 0.150, adjusted P = 0.0305), whereas axial span did not (R² = 0.065, adjusted P = 0.620).

**Extended Data Fig. 4.**
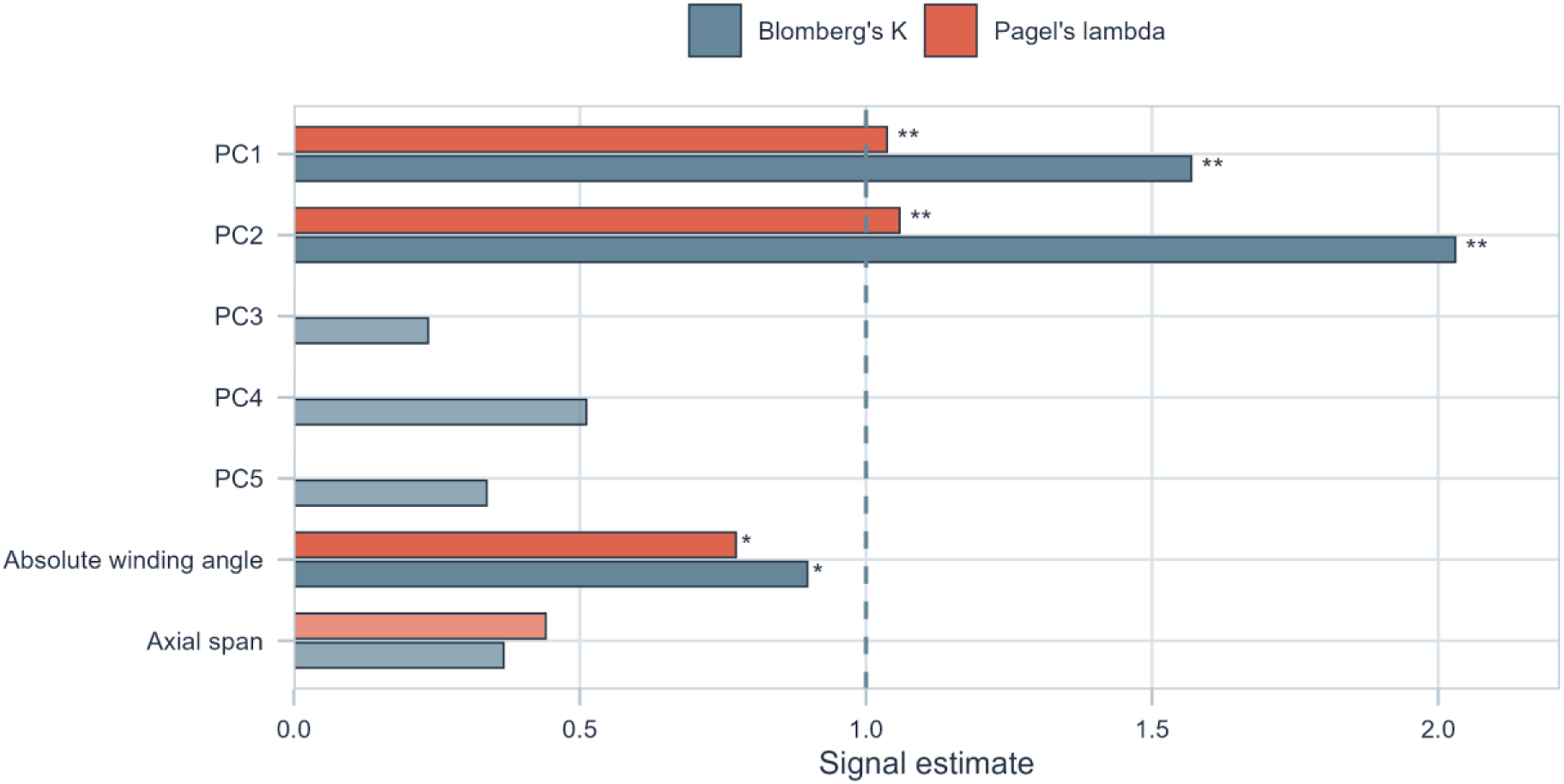
Phylogenetic signal in trochanteral shape and screw joint geometry. **Bars** show Blomberg’s K and Pagel’s λ under the Grafen primary tree. PC1-PC5 use all 15 shape-only proxy tips; winding angle and axial span use the 14 geometry-bearing tips in the main dataset. Darker bars denote FDR-significant estimates. PC1, PC2 and winding angle showed signal, whereas axial span did not.

**Extended Data Fig. 5.**
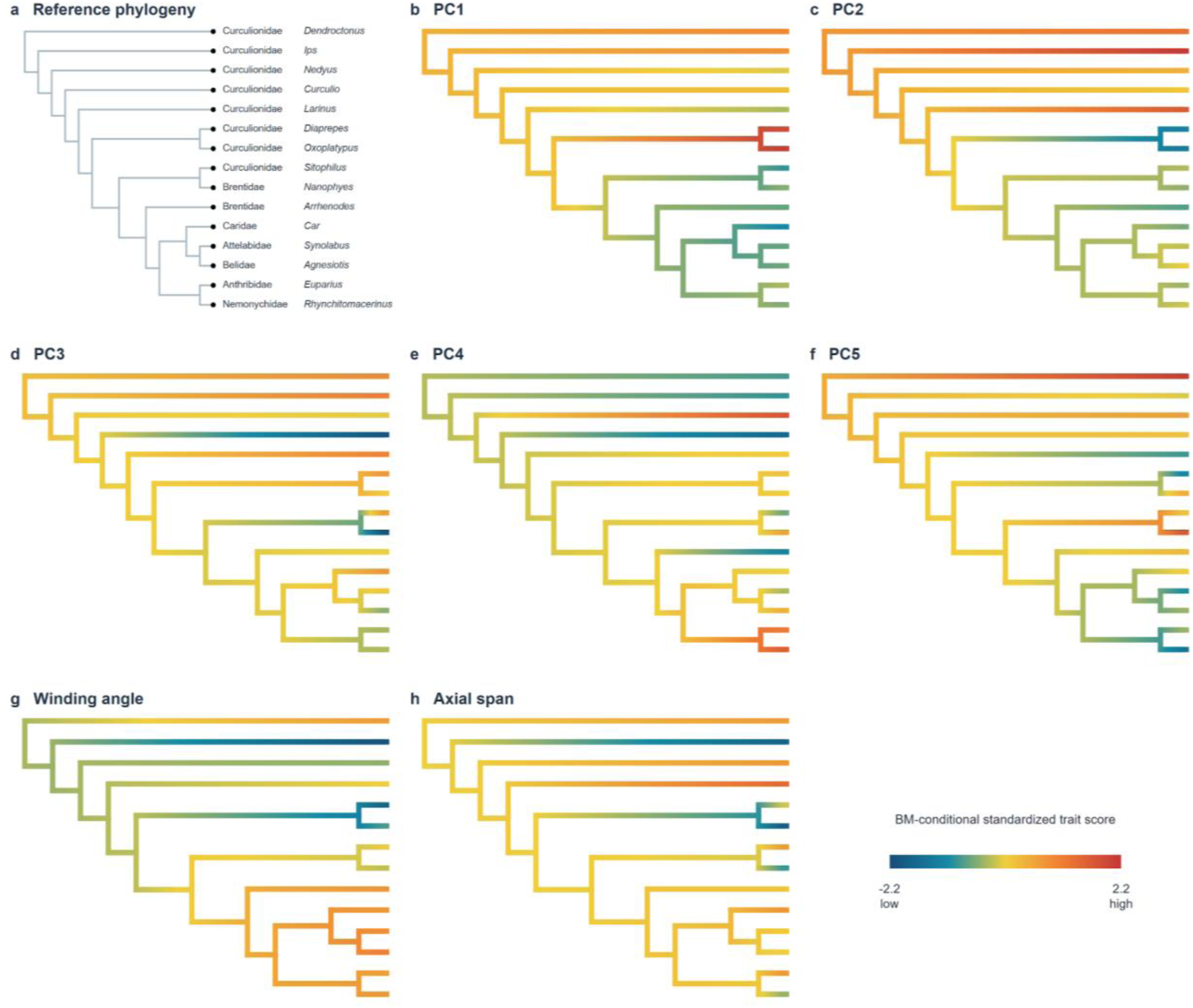
Brownian-motion-conditional ancestral-state reconstructions of major trochanter shape and screw geometry variables. (a) Fifteen-tip shape-only proxy phylogeny under Grafen branch lengths. (b-f) Brownian-motion (BM)-conditional continuous ancestral-state reconstructions for PC1-PC5 on the same 15-tip tree. (g,h) BM-conditional reconstructions for absolute winding angle and axial span on the 14-tip main geometry dataset, omitting the Nemonychidae proxy tip for which geometry values are unavailable. Branch colours indicate standardized trait scores; raw-scale node estimates and 95% intervals for every displayed trait are reported in Supplementary Table 24. Every map assumes BM for reconstruction; this does not imply that BM was top-ranked in the exploratory evolutionary-model comparisons. Because the maps use sparse proxy sampling, within-tip averaging and broader taxonomic assignments, they are conditional visualizations rather than robust genus-, family- or clade-level reconstructions.

**Extended Data Fig. 6.**
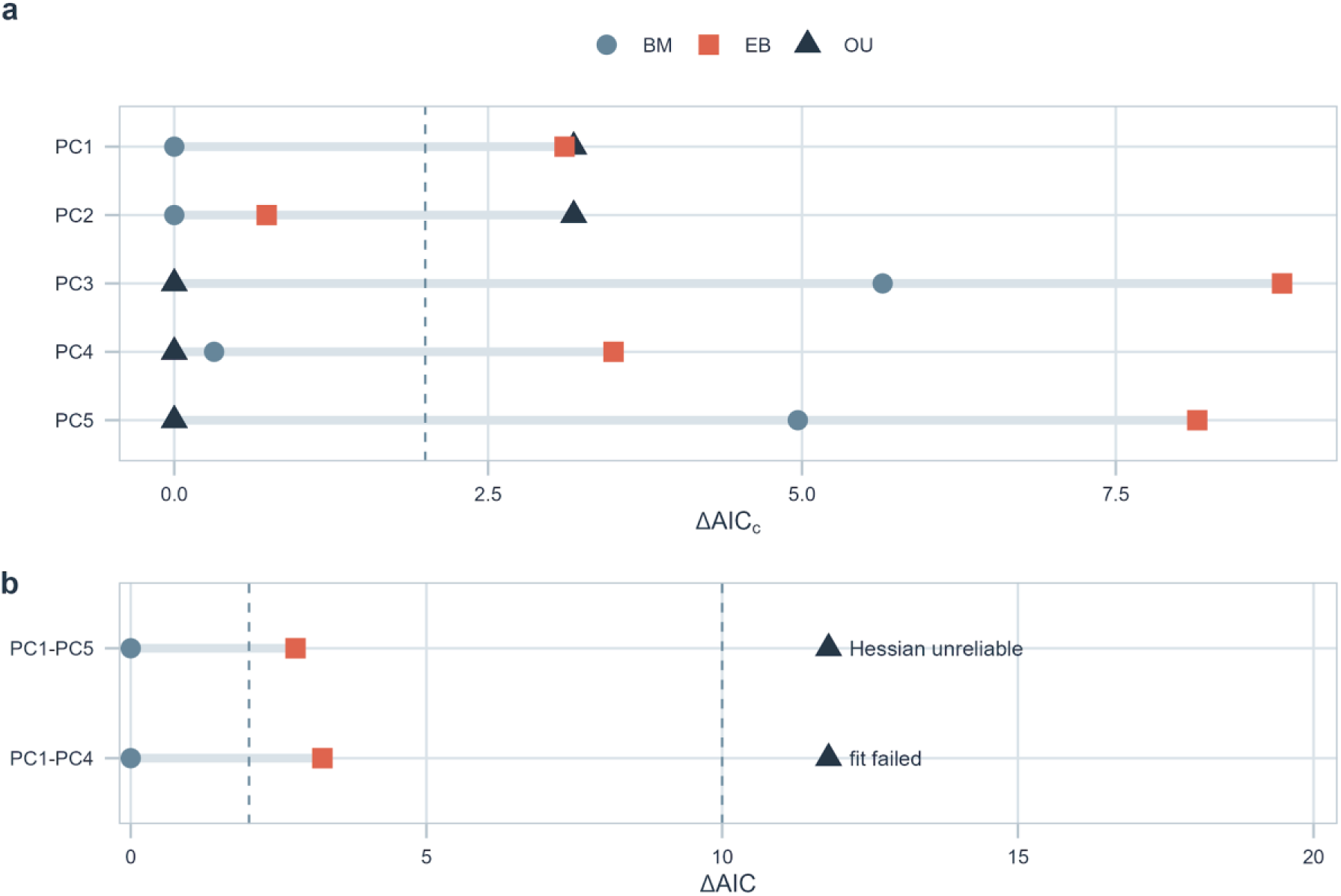
Evolutionary model support for trochanteral shape under the Grafen primary tree. **(a)** ΔAICc among Brownian motion, Ornstein-Uhlenbeck and early-burst univariate models fitted separately to PC1-PC5 at 15 shape-only proxy tips. Brownian motion was top-ranked for PC1 and PC2, whereas Ornstein-Uhlenbeck was top-ranked for PC3-PC5; support for Brownian motion and Ornstein-Uhlenbeck was similar for PC4. (b) ΔAIC among Hessian-reliable multivariate fits at the same 15 shape-only proxy tips. Brownian motion was top-ranked for PC1-PC5 and PC1-PC4; the corresponding early-burst ΔAIC values were 2.79 and 3.24. The PC1-PC5 OU optimizer converged and had the lowest raw AIC, but its Hessian solution was unreliable; the PC1-PC4 OU fit failed. Both OU outcomes are marked separately from the reliable-fit ΔAIC scale, and raw AIC values and diagnostic statuses are retained in Supplementary Table 26. These small-sample comparisons are exploratory and branch-length-dependent.

## References

[1] Arnold, S. J. Morphology, performance and fitness. Am. Zool. 23, 347–361 (1983).

[2] Wainwright, P. C., Alfaro, M. E., Bolnick, D. I. & Hulsey, C. D. Many-to-one mapping of form to function: a general principle in organismal design? Integr. Comp. Biol. 45, 256–262 (2005).

[3] Anderson, P. S. L. & Patek, S. N. Mechanical sensitivity reveals evolutionary dynamics of mechanical systems. Proc. R. Soc. B 282, 20143088 (2015).

[4] Muñoz, M. M., Hu, Y., Anderson, P. S. L. & Patek, S. N. Strong biomechanical relationships bias the tempo and mode of morphological evolution. eLife 7, e37621 (2018).

[5] Gaston, K. J. The magnitude of global insect species richness. Conserv. Biol. 5, 283–296 (1991).

[6] Grimaldi, D. & Engel, M. S. Evolution of the Insects (Cambridge Univ. Press, 2005).

[7] Hunt, T. et al. A comprehensive phylogeny of beetles reveals the evolutionary origins of a superradiation. Science 318, 1913–1916 (2007).

[8] Bouchard, P. et al. Biodiversity of Coleoptera. In Insect Biodiversity: Science and Society (eds Foottit, R. G. & Adler, P. H.) 337–417 (Wiley-Blackwell, 2017).

[9] McKenna, D. D., et al. The evolution and genomic basis of beetle diversity. Proc. Natl Acad. Sci. USA 116, 24729–24737 (2019).

[10] Crowson, R. A. The Biology of the Coleoptera (Academic Press, 1981).

[11] Beutel, R. G. & Haas, F. Phylogenetic relationships of the suborders of Coleoptera (Insecta). Cladistics 16, 103–141 (2000).

[12] Lawrence, J. F. & Slipinski, A. Australian Beetles, Volume 1: Morphology, Classification and Keys (CSIRO Publishing, 2013).

[13] Farrell, B. D. “Inordinate fondness” explained: why are there so many beetles? Science 281, 555–559 (1998).

[14] Wipfler, B., Pohl, H., Yavorskaya, M. I. & Beutel, R. G. A review of methods for analysing insect structures: the role of morphology in the age of phylogenomics. Curr. Opin. Insect Sci. 18, 60–68 (2016).

[15] Oh, J. K., Behmer, S. T., Marquess, R., Scholar, E. A. & Akbulut, M. Investigation of mechanical properties of tibia and femur articulations of insect joints with different joint functions. MRS Commun. 9, 900–903 (2019).

[16] Snodgrass, R. E. Principles of Insect Morphology (McGraw-Hill, 1935).

[17] Frantsevich, L. & Wang, W. Gimbals in the insect leg. Arthropod Struct. Dev. 38, 16–30 (2009).

[18] Beutel, R. G. & Lawrence, J. F. Coleoptera, morphology. In Handbook of Zoology Vol. IV: Arthropoda: Insecta, Coleoptera Vol. I (eds Beutel, R. G. & Leschen, R. A. B.) 23–28 (De Gruyter, 2005).

[19] Chamorro-Lacayo, M. L. & Konstantinov, A. S. Morphology of the prothorax and procoxa in the New World Cryptocephalini (Coleoptera: Chrysomelidae: Cryptocephalinae). Zootaxa 676, 1–46 (2004).

[20] Zong, L. et al. A self-locking mechanism of the frog-legged beetle Sagra femorata. Insect Sci. 31, 1864–1875 (2024).

[21] Reuleaux, F. Die praktischen Beziehungen der Kinematik zu Geometrie und Mechanik. Lehrbuch der Kinematik, Vol. 2 (Friedrich Vieweg und Sohn, 1900).

[22] Reuter, M. Funktionsmorphologische Studien zum Sprung der Stachelkäfer (Coleoptera, Mordellidae). Acta Biol. Benrodis 7, 99–133 (1995).

[23] van de Kamp, T. et al. A biological screw in a beetle’s leg. Science 333, 52 (2011).

[24] van de Kamp, T. et al. Three-dimensional reconstructions come to life: interactive 3D PDF animations in functional morphology. PLoS ONE 9, e102355 (2014).

[25] Oberprieler, R. G., Marvaldi, A. E. & Anderson, R. S. Weevils, weevils, weevils everywhere. Zootaxa 1668, 491–520 (2007).

[26] Betz, O. et al. Imaging applications of synchrotron X-ray phase-contrast microtomography in biological morphology and biomaterials science. I. General aspects of the technique and its advantages in the analysis of millimetre-sized arthropod structure. J. Microsc. 227, 51–71 (2007).

[27] van de Kamp, T. et al. Parasitoid biology preserved in mineralized fossils. Nat. Commun. 9, 3325 (2018).

[28] van de Kamp, T. & Economo, E. P. Large-scale library of 3D ant images captured with synchrotron X-ray microtomography. Nat. Methods 23, 505–506 (2026).

[29] Rühr, P. T. et al. Juvenile ecology drives adult morphology in two insect orders. Proc. R. Soc. B 288, 20210616 (2021).

[30] Katzke, J. et al. High-throughput phenomics of global ant biodiversity. Nat. Methods 23, 663–672 (2026).

[31] Isensee, F., et al. nnInteractive: redefining 3D promptable segmentation. Preprint at https://arxiv.org/abs/2503.08373 (2025).

[32] Lösel, P. D. et al. Natural variability in bee brain size and symmetry revealed by micro-CT imaging and deep learning. PLoS Comput. Biol. 19, e1011529 (2023).

[33] Toulkeridou, E., Gutierrez, C. E., Baum, D., Doya, K. & Economo, E. P. Automated segmentation of insect anatomy from micro-CT images using deep learning. Natural Sciences 3, e20230010 (2023).

[34] Durrleman, S. et al. Morphometry of anatomical shape complexes with dense deformations and sparse parameters. NeuroImage 101, 35–49 (2014).

[35] Larsén, O. On the morphology and function of the locomotor organs of the Gyrinidae and other Coleoptera. Opusc. Entomol. Suppl. 30, 1–242 (1966).

[36] Klingenberg, C. P. Morphological integration and developmental modularity. Annu. Rev. Ecol. Evol. Syst. 39, 115–132 (2008).

[37] Larouche, O., Zelditch, M. L. & Cloutier, R. Modularity promotes morphological divergence in ray-finned fishes. Sci. Rep. 8, 7278 (2018).

[38] Bardua, C. et al. Morphological evolution and modularity of the caecilian skull. BMC Evol. Biol. 19, 30 (2019).

[39] Booher, D. B. et al. Functional innovation promotes diversification of form in the evolution of an ultrafast trap-jaw mechanism in ants. PLoS Biol. 19, e3001031 (2021).

[40] Coombs, E. J. et al. The tempo of cetacean cranial evolution. Curr. Biol. 32, 2233–2247.e4 (2022).

[41] Marvaldi, A. E., Oberprieler, R. G., Lyal, C. H. C., Bradbury, T. & Anderson, R. S. Phylogeny of the Oxycoryninae sensu lato (Coleoptera: Belidae) and evolution of host-plant associations. Invertebr. Syst. 20, 447–476 (2006).

[42] Marvaldi, A. E., Sequeira, A. S., O’Brien, C. W. & Farrell, B. D. Molecular and morphological phylogenetics of weevils (Coleoptera, Curculionoidea): do niche shifts accompany diversification? Syst. Biol. 51, 761–785 (2002).

[43] Shin, S. et al. Phylogenomic data yield new and robust insights into the phylogeny and evolution of weevils. Mol. Biol. Evol. 35, 823–836 (2018).

[44] Li, Y.-D., Engel, M. S., Tihelka, E. & Cai, C. Phylogenomics of weevils revisited: data curation and modelling compositional heterogeneity. Biol. Lett. 19, 20230307 (2023).

[45] McKenna, D. D. et al. The beetle tree of life reveals that Coleoptera survived end-Permian mass extinction to diversify during the Cretaceous terrestrial revolution. Syst. Entomol. 40, 835–880 (2015).

[46] Klingenberg, C. P. Size, shape, and form: concepts of allometry in geometric morphometrics. Dev. Genes Evol. 226, 113–137 (2016).

[47] Pélabon, C. et al. Evolution of morphological allometry. Ann. N.Y. Acad. Sci. 1320, 58–75 (2014).

[48] McKenna, D. D., Farrell, B. D., Marvaldi, A. E., Oberprieler, R. G. & Li, X. Phylogenomics of palynophagous pine cone weevils (Coleoptera: Cimberididae) recovers the monophyly of Cimberidini and Doydirhynchini and reveals the paraphyly of Cimberis. Eur. J. Entomol. 121, 435–442 (2024).

[49] Wagner, G. P. & Altenberg, L. Complex adaptations and the evolution of evolvability. Evolution 50, 967–976 (1996).

[50] Muñoz, M. M., Anderson, P. S. L. & Patek, S. N. Mechanical sensitivity and the dynamics of evolutionary rate shifts in biomechanical systems. Proc. R. Soc. B 284, 20162325 (2017).

[51] Hoebeke, E. R. & Wheeler, A. G. Anthribus nebulosus, a Eurasian scale predator in the eastern United States (Coleoptera: Anthribidae): notes on biology, recognition, and establishment. Proc. Entomol. Soc. Wash. 93, 45–50 (1991).

[52] Schultz, P. B. & Muhleman, C. E. Recovery and distribution of Anthribus nebulosus, a scale predator introduced into Virginia in 1981. Banisteria 47, 14–17 (2016).

[53] Janicki, J. & Young, D. K. Nemonychidae and Anthribidae of Wisconsin (Coleoptera: Curculionoidea). Insecta Mundi 0579, 1–36 (2017).

[54] Alba-Alejandre, I., Alba-Tercedor, J. & Vega, F. E. Micro-CT to document the coffee bean weevil, Araecerus fasciculatus (Coleoptera: Anthribidae), inside field-collected coffee berries (Coffea canephora). Insects 9, 100 (2018).

[55] Machado, A. Some additions and corrections to the Coleoptera fauna of the Canary Islands. Graellsia 70, e005 (2014).

[56] Ptaszyńska, A. A., Łętowski, J., Gnat, S. & Małek, W. Application of COI sequences in studies of phylogenetic relationships among 40 Apionidae species. J. Insect Sci. 12, 16 (2012).

[57] Hundsdoerfer, A. K., Rheinheimer, J. & Wink, M. Towards the phylogeny of the Curculionoidea (Coleoptera): reconstructions from mitochondrial and nuclear ribosomal DNA sequences. Zool. Anz. 248, 9–31 (2009).

[58] Winter, S., Friedman, A. L. L., Astrin, J. J., Gottsberger, B. & Letsch, H. Timing and host plant associations in the evolution of the weevil tribe Apionini (Apioninae, Brentidae, Curculionoidea, Coleoptera) indicate an ancient co-diversification pattern of beetles and flowering plants. Mol. Phylogenet. Evol. 107, 179–190 (2017).

[59] Kobayashi, C., Okuyama, Y., Kawazoe, K. & Kato, M. The evolutionary history of maternal plant-manipulation and larval feeding behaviours in attelabid weevils (Coleoptera; Curculionoidea). Mol. Phylogenet. Evol. 64, 318–330 (2012).

[60] Wang, L. et al. Farming of a defensive fungal mutualist by an attelabid weevil. ISME J. 9, 1793–1801 (2015).

[61] Schowalter, T. D. Insect Ecology: An Ecosystem Approach 4th edn (Academic Press, 2016).

[62] McKenna, D. D., Sequeira, A. S., Marvaldi, A. E. & Farrell, B. D. Temporal lags and overlap in the diversification of weevils and flowering plants. Proc. Natl Acad. Sci. USA 106, 7083–7088 (2009).

[63] Haran, J. et al. Phylogenomics illuminates the phylogeny of flower weevils (Curculioninae) and reveals ten independent origins of brood-site pollination mutualism in true weevils. Proc. R. Soc. B 290, 20230889 (2023).

[64] van de Kamp, T. et al. Comparative thorax morphology of death-feigning flightless cryptorhynchine weevils (Coleoptera: Curculionidae) based on 3D reconstructions. Arthropod Struct. Dev. 44, 509–523 (2015).

[65] Yunakov, N., Nazarenko, V., Filimonov, R. & Volovnik, S. A survey of the weevils of Ukraine (Coleoptera: Curculionoidea). Zootaxa 4404, 1–494 (2018).

[66] Cecilia, A. et al. The IMAGE beamline at the KIT Light Source. J. Synchrotron Radiat. 32, 1036–1051 (2025).

[67] Douissard, P.-A. et al. A versatile indirect detector design for hard X-ray microimaging. J. Instrum. 7, P09016 (2012).

[68] Vogelgesang, M. et al. Real-time image-content-based beamline control for smart 4D X-ray imaging. J. Synchrotron Radiat. 23, 1254–1263 (2016).

[69] Vogelgesang, M. et al. UFO: A scalable GPU-based image processing framework for on-line monitoring. In 2012 IEEE 14th International Conference on High Performance Computing and Communication & 2012 IEEE 9th International Conference on Embedded Software and Systems 824-829 (IEEE, 2012).

[70] Faragó, T. et al. Tofu: a fast, versatile and user-friendly image processing toolkit for computed tomography. J. Synchrotron Radiat. 29, 916–927 (2022).

[71] Stalling, D., Westerhoff, M. & Hege, H.-C. Amira: a highly interactive system for visual data analysis. In The Visualization Handbook (eds Hansen, C. D. & Johnson, C. R.) 749–767 (Elsevier, 2005).

[72] napari contributors. napari: a multi-dimensional image viewer for Python. Zenodo 10.5281/zenodo.3555620 (2019).

[73] Minh, B. Q. et al. IQ-TREE 2: new models and efficient methods for phylogenetic inference in the genomic era. Mol. Biol. Evol. 37, 1530–1534 (2020).

[74] Paradis, E. & Schliep, K. ape 5.0: an environment for modern phylogenetics and evolutionary analyses in R. Bioinformatics 35, 526–528 (2019).

[75] Letsch, H., Balke, M., Toussaint, E. F. A. & Riedel, A. Historical biogeography of the hyperdiverse hidden snout weevils (Coleoptera, Curculionidae, Cryptorhynchinae). Syst. Entomol. 45, 312–326 (2020).

[76] Fikáček, M. et al. Reliable placement of beetle fossils via phylogenetic analyses: Triassic *Leehermania* as a case study (Staphylinidae or Myxophaga?). Syst. Entomol. 45, 175–187 (2020).

[77] Li, X. et al. The evolutionary history of the ancient weevil family Belidae (Coleoptera: Curculionoidea) reveals the marks of Gondwana breakup and major floristic turnovers, including the rise of angiosperms. eLife 13, RP97552 (2024).

[78] Scrucca, L., Fop, M., Murphy, T. B. & Raftery, A. E. mclust 5: clustering, classification and density estimation using Gaussian finite mixture models. R J. 8, 289–317 (2016).

[79] Foote, M. Contributions of individual taxa to overall morphological disparity. Paleobiology 19, 403–419 (1993).

[80] Guillerme, T. et al. Disparities in the analysis of morphological disparity. Biol. Lett. 16, 20200199 (2020).

[81] Collyer, M. L. & Adams, D. C. RRPP: an R package for fitting linear models to high-dimensional data using residual randomization. Methods Ecol. Evol. 9, 1772–1779 (2018).

[82] Adams, D. C. & Otárola-Castillo, E. geomorph: an R package for the collection and analysis of geometric morphometric shape data. Methods Ecol. Evol. 4, 393–399 (2013).

[83] Benjamini, Y. & Hochberg, Y. Controlling the false discovery rate: a practical and powerful approach to multiple testing. J. R. Stat. Soc. B 57, 289–300 (1995).

[84] Adams, D. C. & Collyer, M. L. Multivariate phylogenetic comparative methods: evaluations, comparisons and recommendations. Syst. Biol. 67, 14–31 (2018).

[85] Garamszegi, L. Z. (ed.) Modern Phylogenetic Comparative Methods and Their Application in Evolutionary Biology: Concepts and Practice (Springer, 2014).

[86] Pennell, M. W. et al. geiger v2.0: an expanded suite of methods for fitting macroevolutionary models to phylogenetic trees. Bioinformatics 30, 2216–2218 (2014).

[87] Pinheiro, J. C. & Bates, D. M. Mixed-Effects Models in S and S-PLUS (Springer, 2000).

[88] Paradis, E., Claude, J. & Strimmer, K. APE: analyses of phylogenetics and evolution in R language. Bioinformatics 20, 289–290 (2004).

[89] Revell, L. J. phytools 2.0: an updated R ecosystem for phylogenetic comparative methods (and other things). PeerJ 12, e16505 (2024).

[90] Wickham, H. ggplot2: Elegant Graphics for Data Analysis (Springer, 2016).

