## Supplementary Material for "Evolutionary diversification of the biological screw joint in weevils"

### Supplements

#### Supplementary Figures

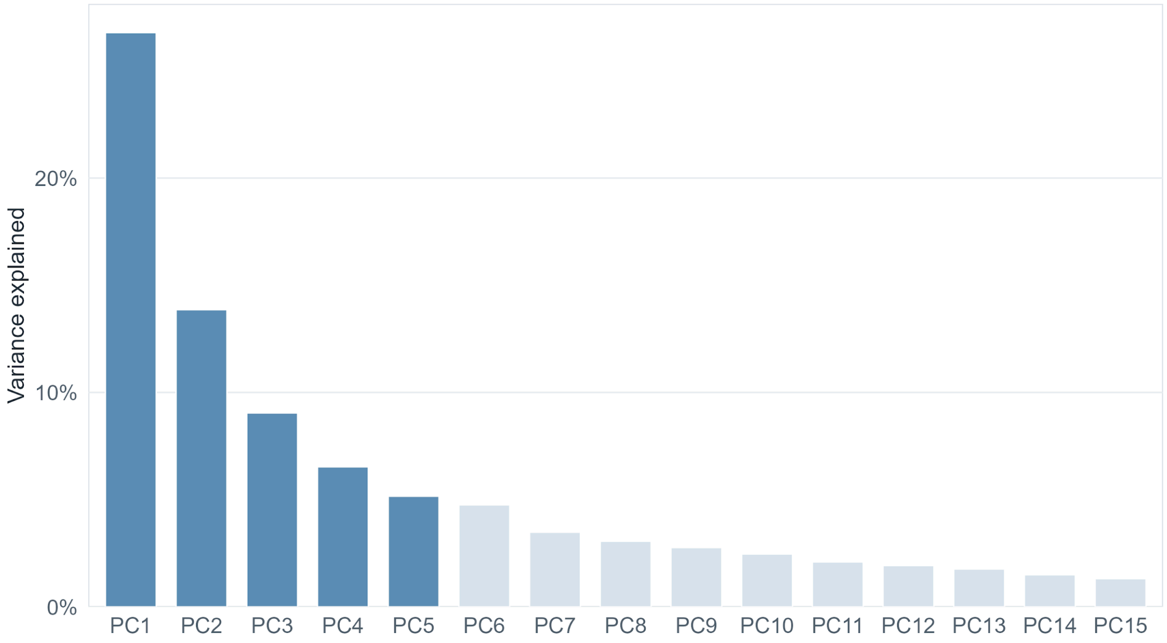


**Supplementary Figure 1. Variance explained by principal components.**

Bar plot showing the percentage of total trochanteral shape variation explained by the first 15 of 67 non-zero principal components. Dark blue bars indicate components above the approximately 5% per-axis retention threshold and correspond to PC1-PC5; light blue bars show subsequent lower-variance components.

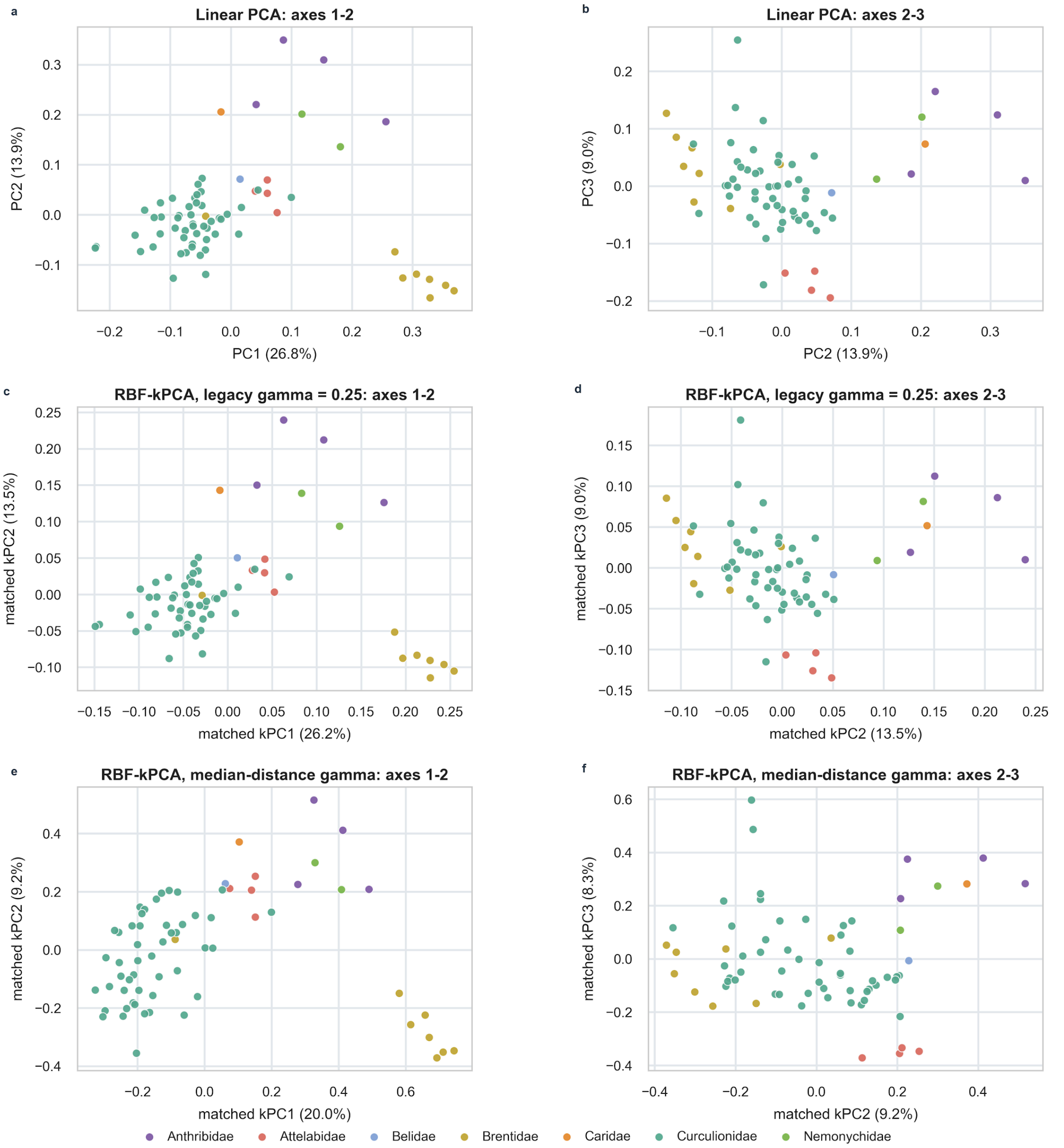


**Supplementary Figure 2. Sensitivity of the atlas-derived morphospace to linear and nonlinear ordination.**
**(a,b)** Ordinary PCA of centred, vectorized subject-specific momenta. **(c,d)** RBF-kPCA using γ = 0.25, retained from the original workflow. **(e,f)** RBF-kPCA using a data-adaptive median-distance kernel. Panels show the first two and second and third matched axes. Points represent specimens and are coloured by family. Both kPCA parameterizations retained the principal family-level structure observed with ordinary PCA. Quantitative comparisons of axis correspondence, interspecimen distances and downstream tests are provided in Supplementary Table 4.

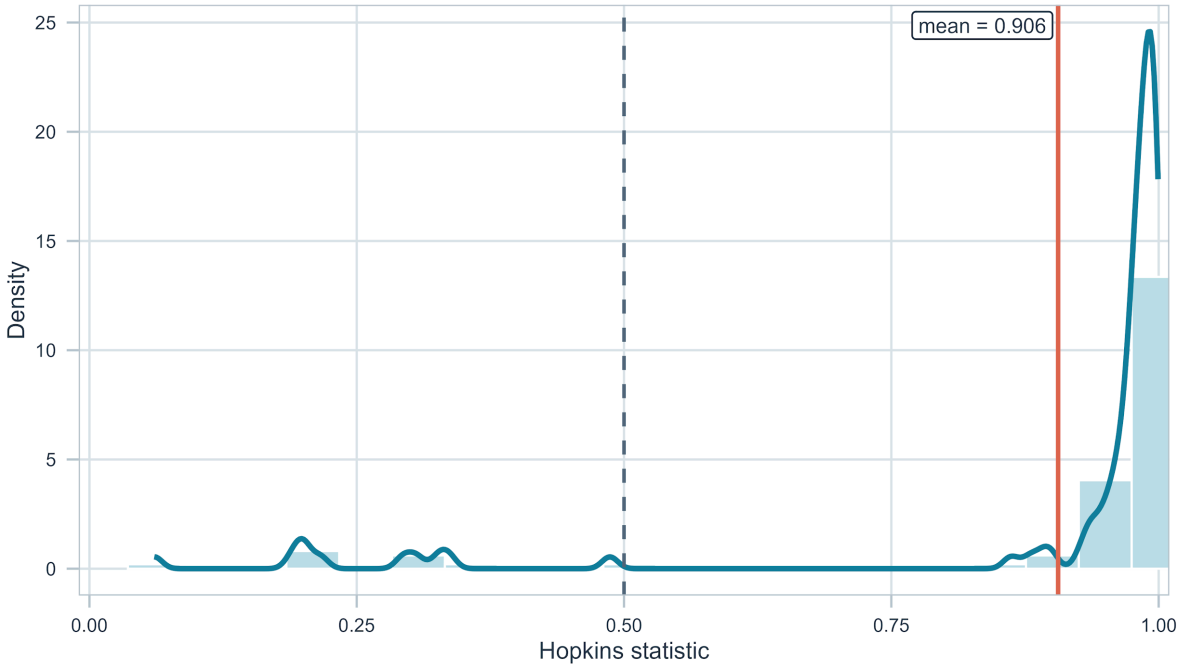


**Supplementary Figure 3. Clusterability of the trochanteral morphospace.**

Distribution of 100 randomized Hopkins evaluations calculated from standardized PC1-PC5 scores (n = 68, m = 6, seed = 1). Values close to 1 indicate stronger spatial clustering than expected under spatial randomness. The orange line marks the mean (0.906), and the dashed line marks the 0.5 reference value. Because the sample did not permit conventional large-sample conditions for formal Hopkins inference, the distribution is interpreted only as an exploratory clusterability diagnostic.

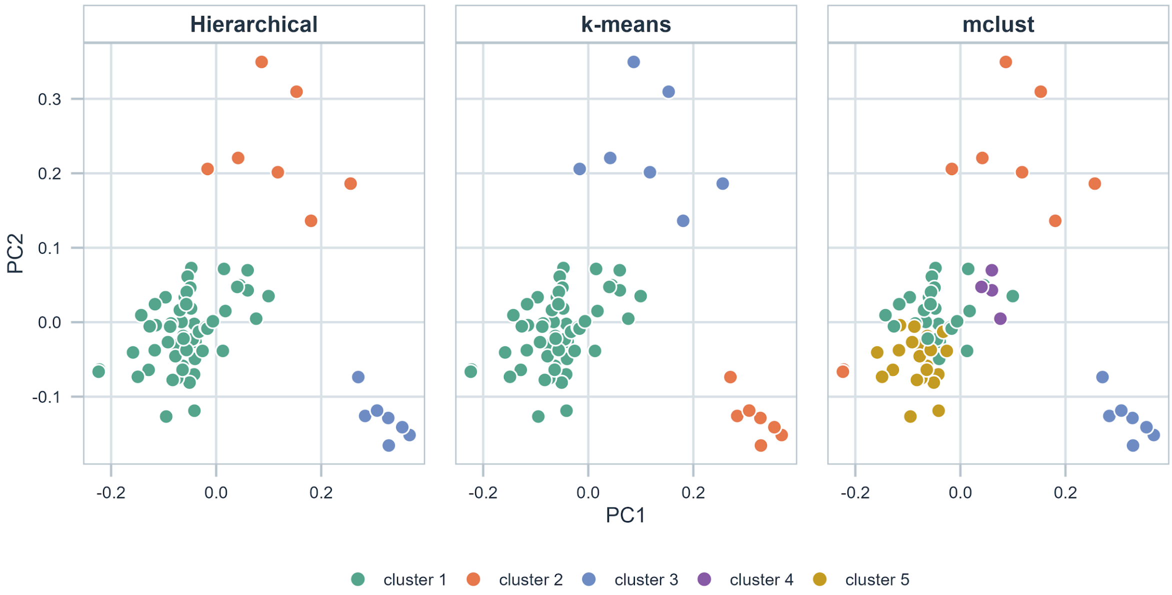


**Supplementary Figure 4. Comparison of clustering outcomes across alternative methods.** Specimens are plotted in PC1-PC2 morphospace with cluster assignments inferred separately from standardized PC1-PC5 scores by hierarchical clustering, k-means clustering and model-based clustering (mclust). Colours denote method-specific cluster assignments and should not be interpreted as corresponding clusters across panels. Hierarchical clustering and k-means each recovered three clusters, whereas mclust recovered five clusters. This method dependence does not support a single robust discrete partition of the morphospace.

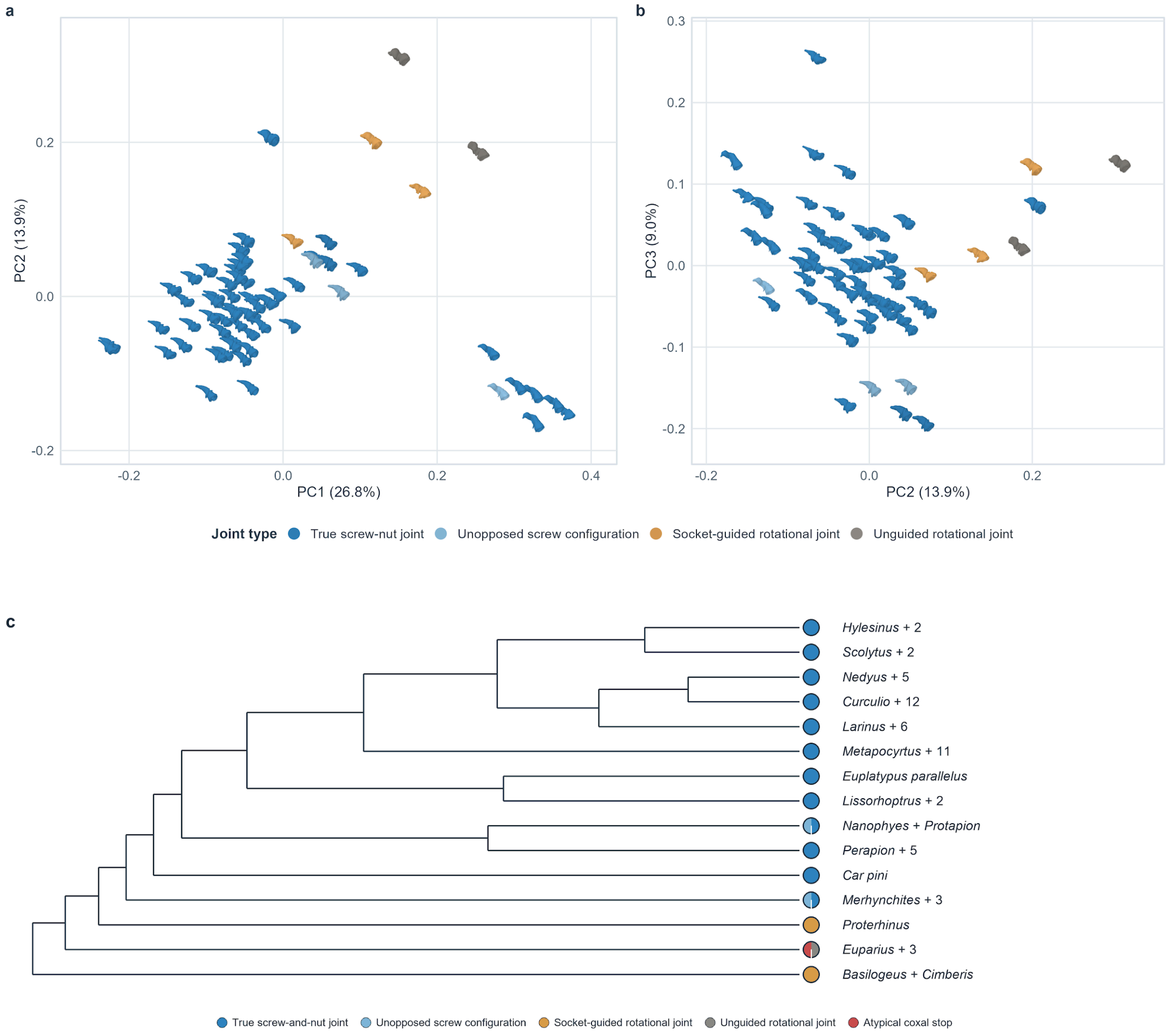


**Supplementary Figure 5. Joint types across morphospace and phylogeny.**
**(a,b)** Two-dimensional projections of trochanteral morphospace, with specimens coloured by joint type. **(c)** Phylogenetic distribution of joint type assignments after specimens were assigned to the tree tips used for comparative analyses. Pie charts at terminal branches show the composition of sampled specimens assigned to each tree tip; solid circles indicate tips represented by a single joint type. The atypical coxal-stop category denotes *Anthribus nebulosus* and *Urodon rufipes*, which were retained as descriptive observations but not included in the four recurrent joint-type categories used for group-wise statistics.

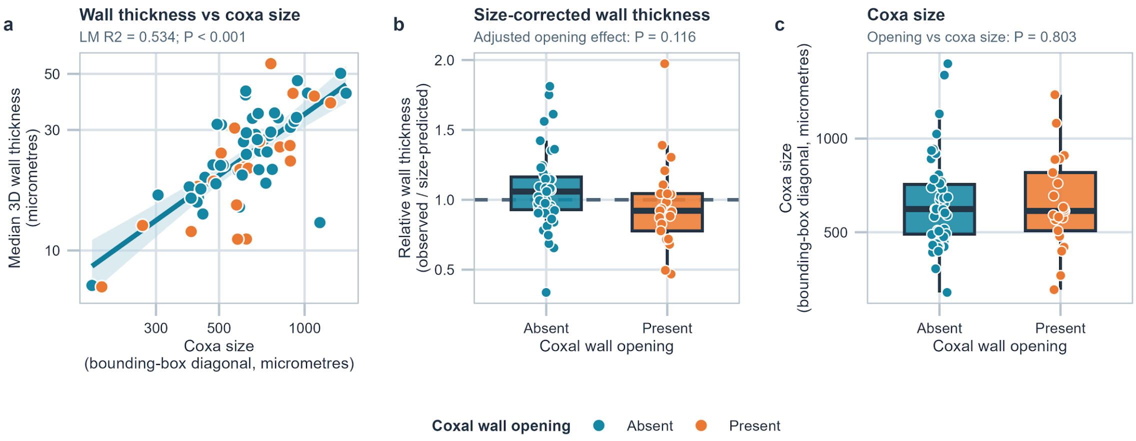


**Supplementary Figure 6. Whole-volume three-dimensional coxal wall thickness, coxa size and coxal wall opening.**

**(a)** Median local wall thickness across the complete 3D coxa mask increased with coxa bounding-box diagonal (log-log LM, $R^2 = 0.534$, $P < 0.001$). Local thickness was defined at each foreground voxel as the diameter of the largest sphere fully contained within the coxa mask and containing that voxel. **(b)** After accounting for coxa size, opening-bearing coxae had 11.5% lower median whole-volume wall thickness than coxae without an opening, but this difference was not statistically significant ($P = 0.116$). **(c)** Coxa size did not differ between opening states (logistic regression, $P = 0.803$). Points represent individual specimens ($n = 68$).

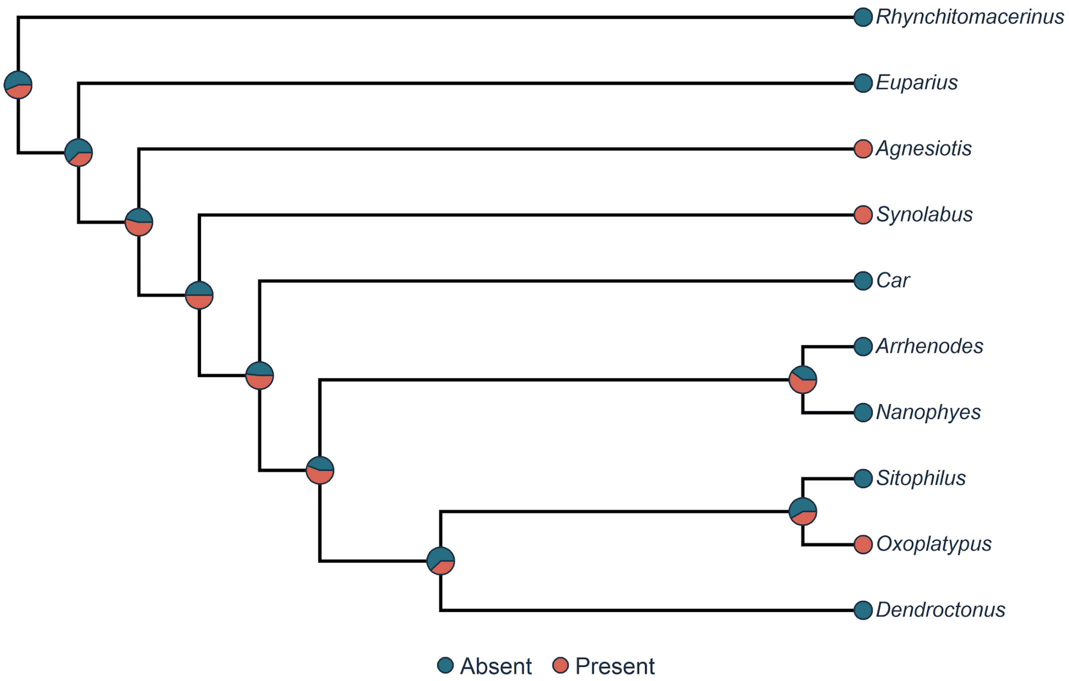


**Supplementary Figure 7. Stochastic character map of coxal wall opening states.** Stochastic character map of coxal wall opening across the 10 taxonomic proxy tips with unambiguous states on the topology-based Grafen primary tree. Tip colours show the observed states; pie charts at internal nodes summarize the proportions of 50 equal-rates stochastic maps assigning each state. The reconstruction is conditional on proxy assignment, within-tip aggregation and the selected topology and is not a genus-level ancestral reconstruction.

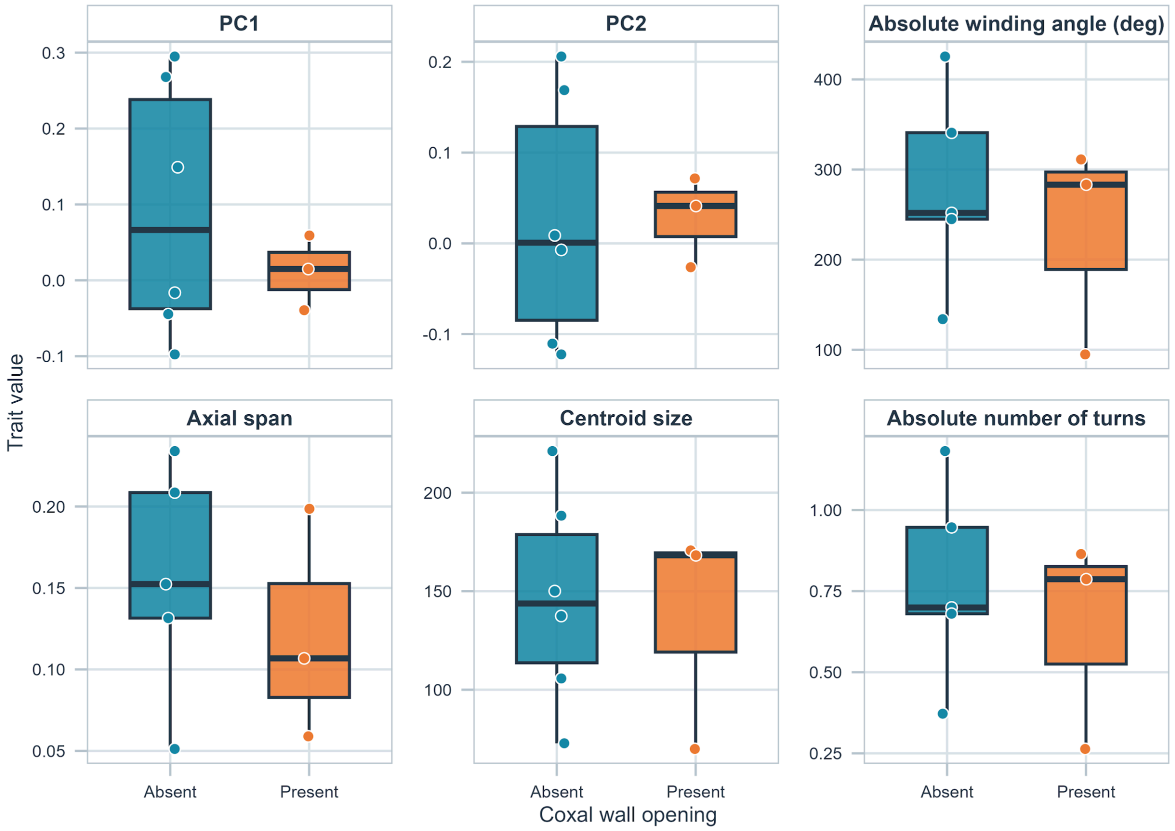


**Supplementary Figure 8. Trait contrasts associated with the coxal opening character.** Descriptive boxplots of taxonomic proxy-tip means from the main dataset for PC1, PC2, robust fitted winding angle, axial span, centroid size and absolute turn number across the two coxal-wall-opening states. The panels visualize the sparse groups only; no geometry association with opening state remained significant after correction, and the proxy-tip design does not establish a functional effect.

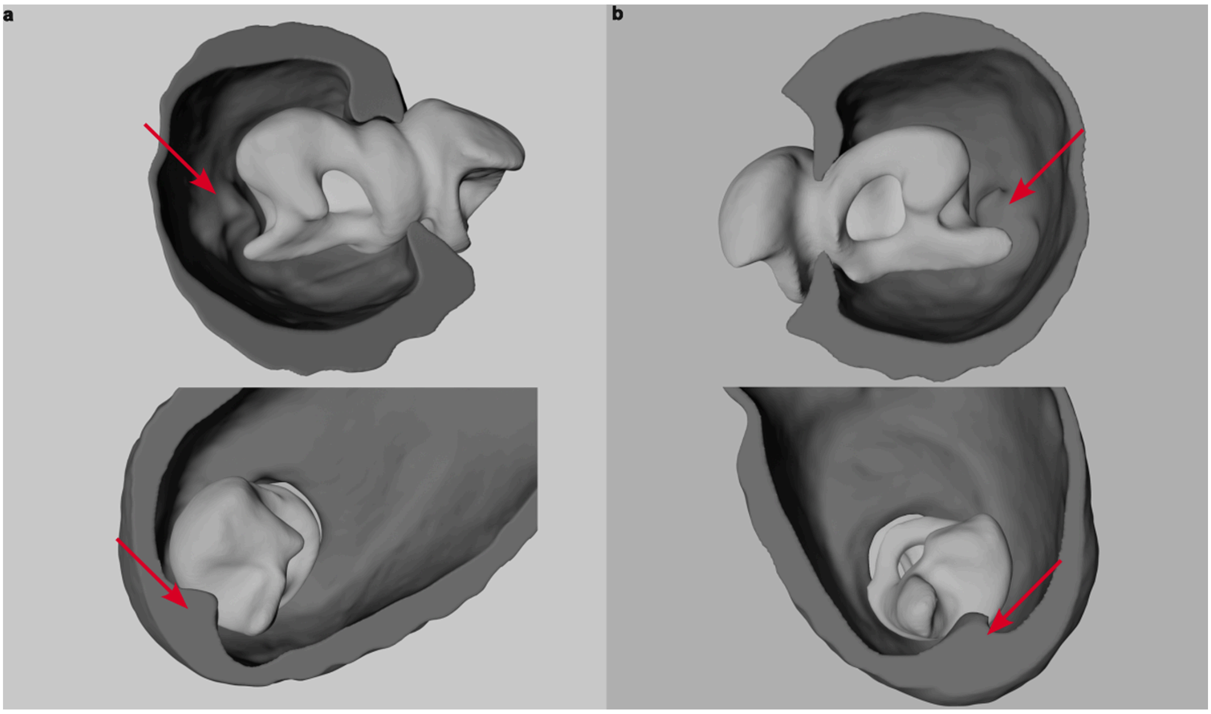


**Supplementary Figure 9. Unilateral coxal stop-like structures in two anthribids.**
Cross-sectional three-dimensional views of the coxa-trochanteral joint in **(a)** *Anthribus nebulosus* and **(b)** *Urodon rufipes*. Within each panel, the upper and lower images show the same joint from different perspectives. Red arrows indicate the stop-like structure on the coxal wall. The trochanter is shown in light grey and the coxa in dark grey.

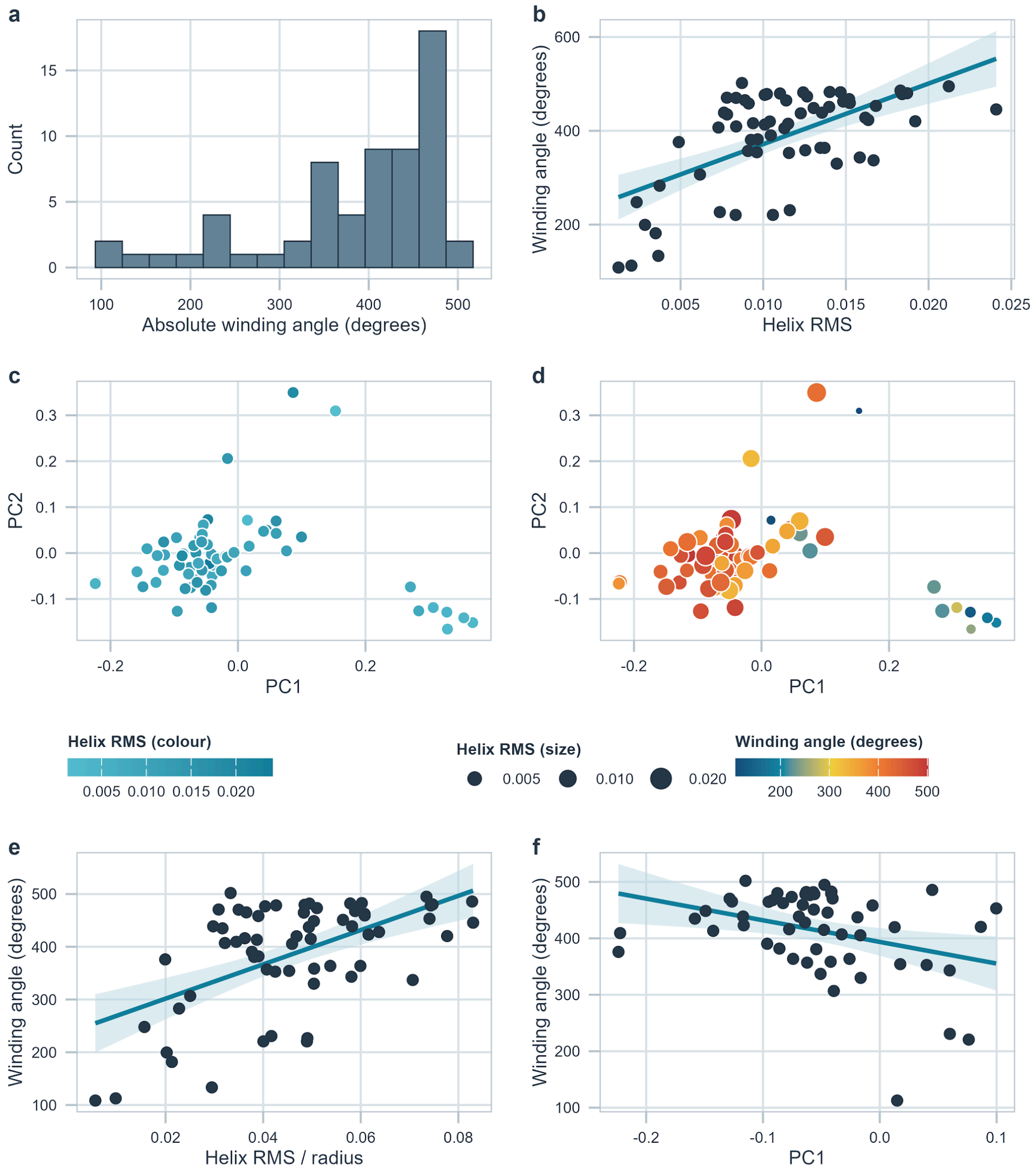


**Supplementary Figure 10. Shape-geometry association and fit-quality diagnostics in the main dataset. (a)** Distribution of absolute winding angle. **(b)** Relationship between winding angle and helix RMS. **(c)** PC1-PC2 morphospace coloured by helix RMS. **(d)** PC1-PC2 morphospace coloured by winding angle, with point size scaled by helix RMS. **(e)** Relationship between winding angle and helix RMS normalised by fitted radius. **(f)** Relationship between winding angle and PC1 within the main morphospace region.

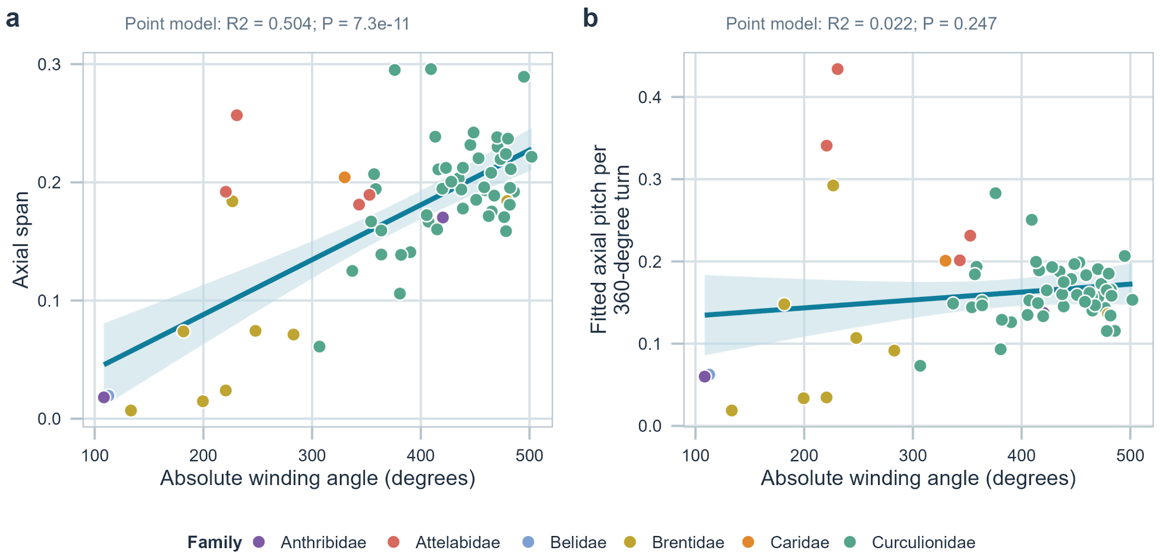


**Supplementary Figure 11. Axial relationships among screw joint geometry variables.** Relationships of absolute winding angle with axial span (a) and fitted axial pitch per 360° (b) across the 63 trajectories in the main dataset. Both panels use the untransformed measurement scales; the fitted lines and reported statistics are from the same ordinary least-squares point models. The angle-span relationship was strong ($R^2 = 0.504$, $P = 7.34 \times 10^{-11}$), whereas the angle-pitch relationship was unsupported ($R^2 = 0.022$, $P = 0.247$).

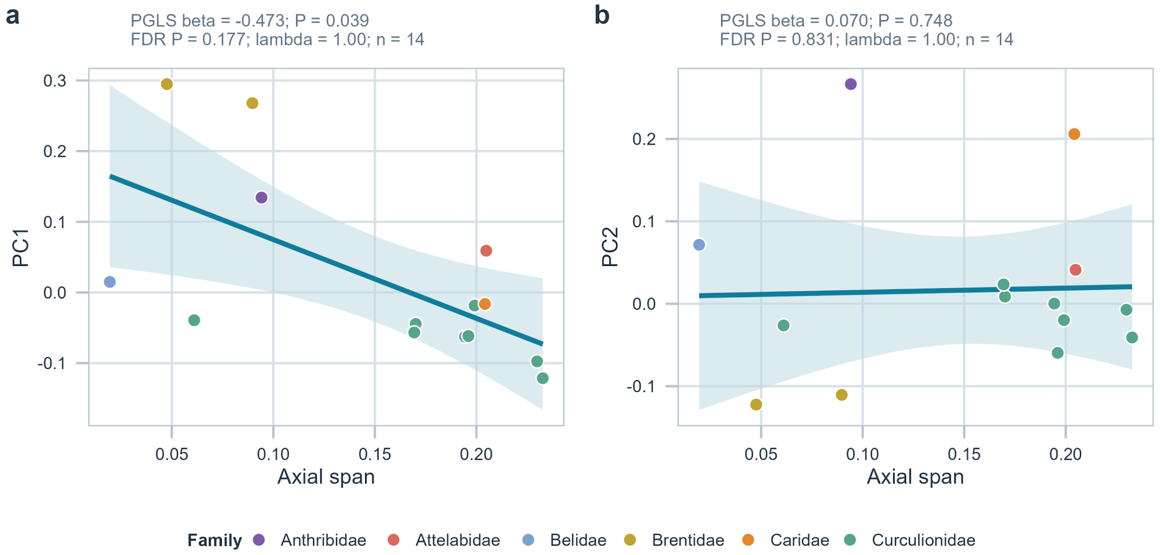


**Supplementary Figure 12. Phylogenetically informed relationships between trochanteral shape and axial span.**

PGLS relationships under the topology-based Grafen primary tree linking PC1 and PC2 to axial span at 14 matched proxy tips in the main dataset. The PC1 slope (a) was negative and nominally significant (β = -0.473, raw P = 0.039) but not after false-discovery-rate correction (FDR-adjusted P = 0.177). PC2 (b) was unsupported. Analyses across 15 tree representations and leave-one-out refits show that the PC1 result is tree- and influence-sensitive.

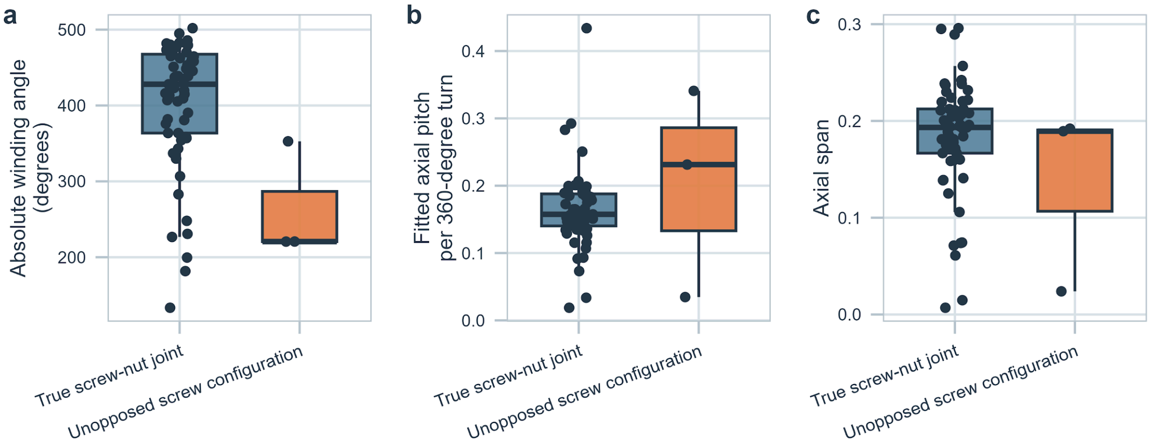


**Supplementary Figure 13. Screw joint geometry across structural joint types.**
Comparison of absolute winding angle, fitted axial pitch and axial span between 57 true screw-and-nut joints and three unopposed screw configurations in the main dataset. Winding angle (a) differed nominally ($P = 0.0202$) but not after correction (adjusted $P = 0.0605$); pitch (b) ($P = 0.387$) and span (c) ($P = 0.317$) were unsupported. Because group sizes were highly unequal, the boxplots and comparisons are exploratory.

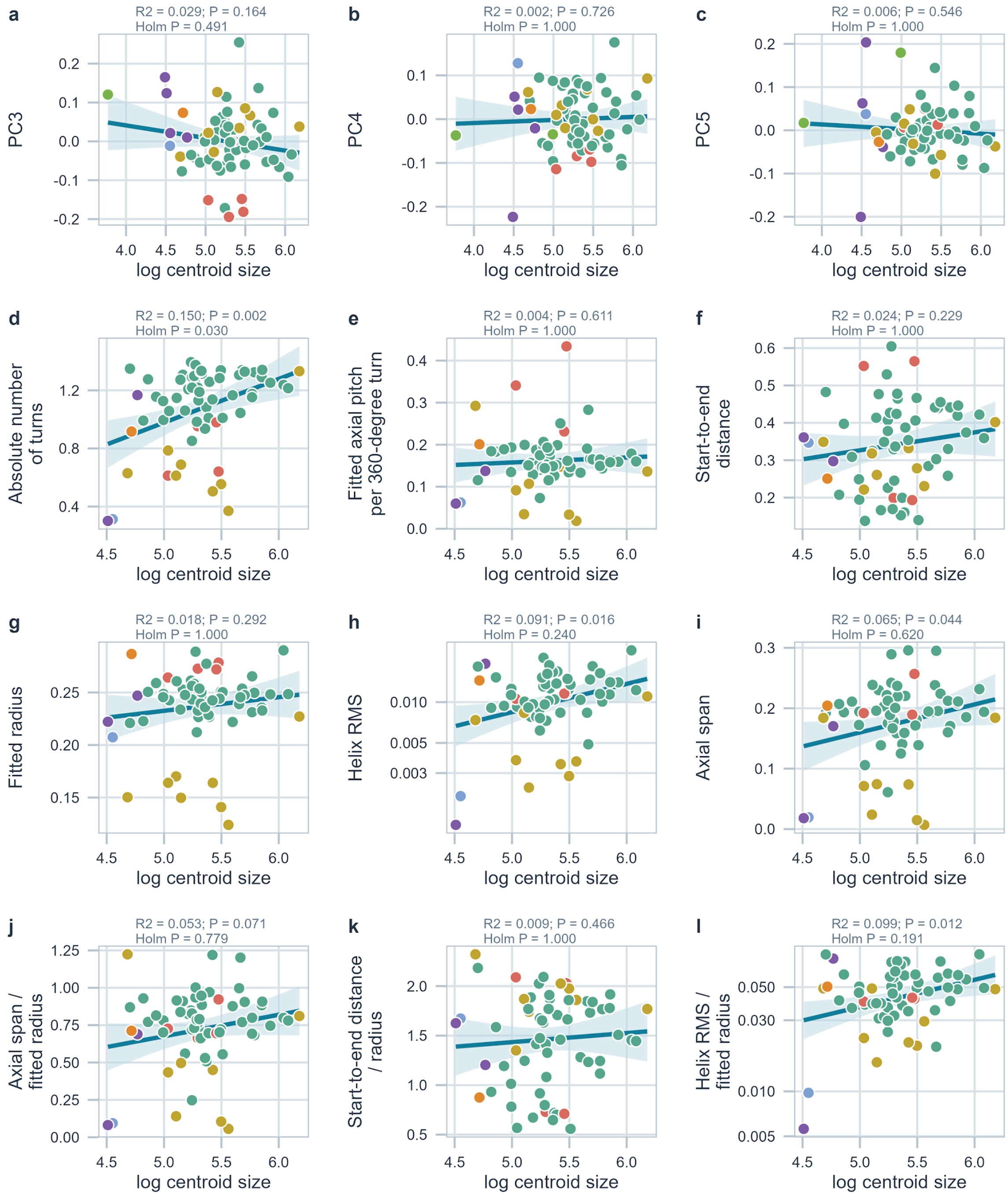


**Supplementary Figure 14. Additional specimen-level allometry plots.**

Additional allometry plots for PC3-PC5 in the full 68-specimen shape dataset and for robust geometry variables in the 63-specimen main geometry dataset, including absolute turn number, axial span and pitch, start-to-end distance, fitted radius, helix-fit error and ratio diagnostics. Among the plotted geometry variables, absolute turn number retained a Holm-corrected association with size; the other plotted relationships did not survive correction.

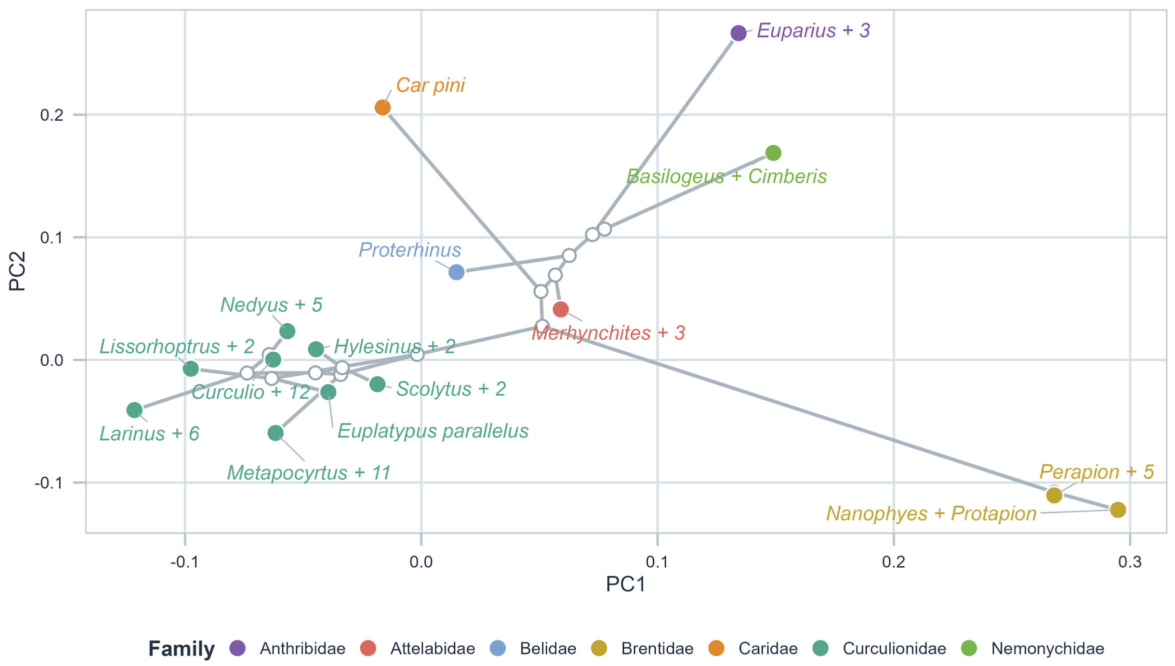


**Supplementary Figure 15. Phylomorphospace of trochanteral shape in PC1-PC2 space.**

Phylomorphospace projection of the topology-based Grafen 15-tip taxonomic proxy tree onto PC1-PC2. Terminal points are proxy-tip means and connecting branches are conditional reconstructions under the selected assignments and topology. The figure does not represent densely sampled family-level trajectories.

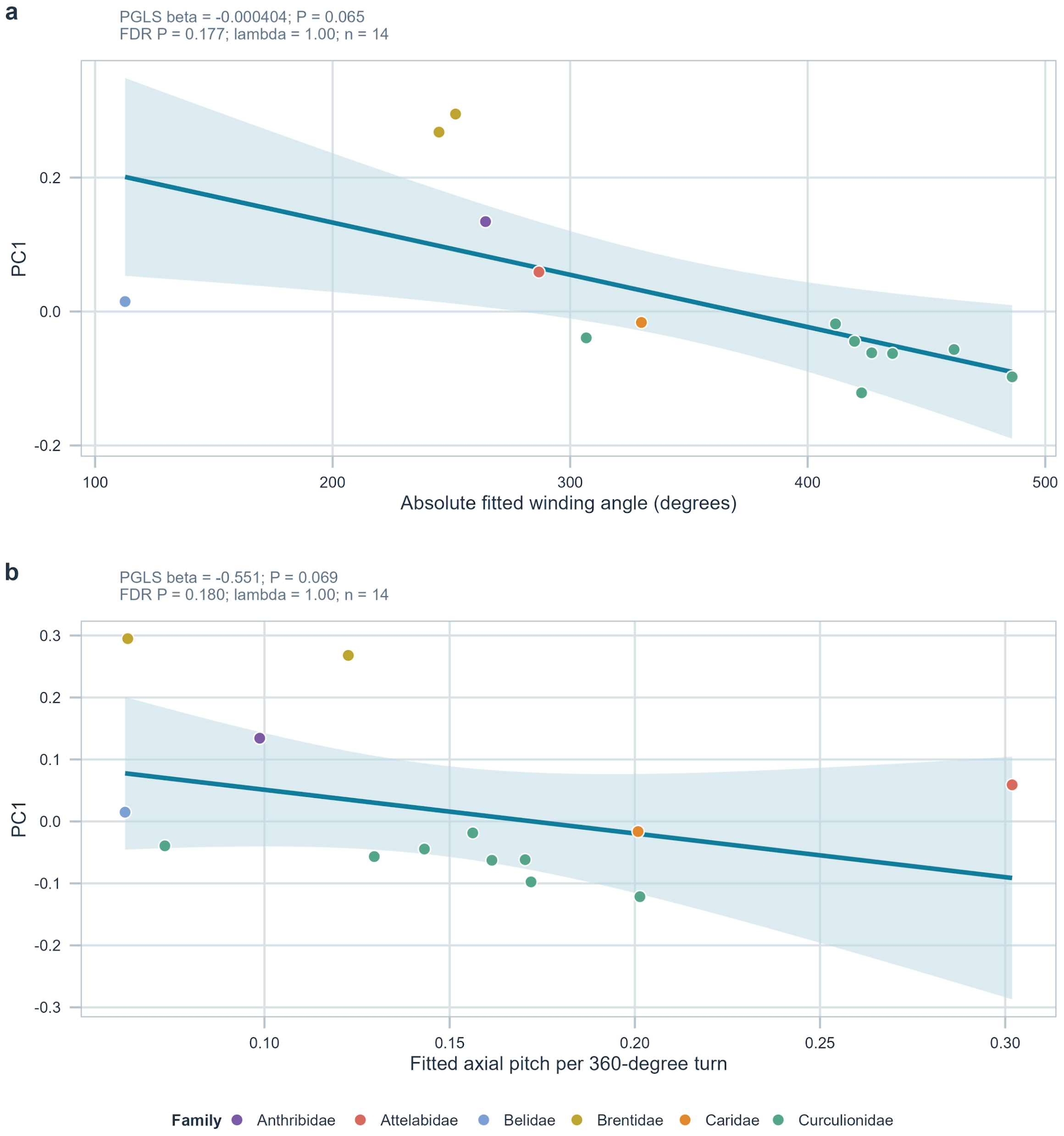


**Supplementary Figure 16. PGLS relationships in the main dataset between PC1 and screw joint geometry.**

The 14 taxonomic proxy-tip means retained in the main dataset are shown under the topology-based Grafen primary tree for (a) PC1 and absolute fitted winding angle and (b) PC1 and fitted axial pitch per 360-degree turn. Lines and 95% confidence bands are ordinary least-squares visual guides only. Both exploratory PGLS associations were negative but unsupported after false-discovery-rate correction (winding angle: $\beta$ = -0.000404, raw P = 0.065, FDR-adjusted P = 0.177; axial pitch: $\beta$ = -0.551, raw P = 0.069, FDR-adjusted P = 0.180).

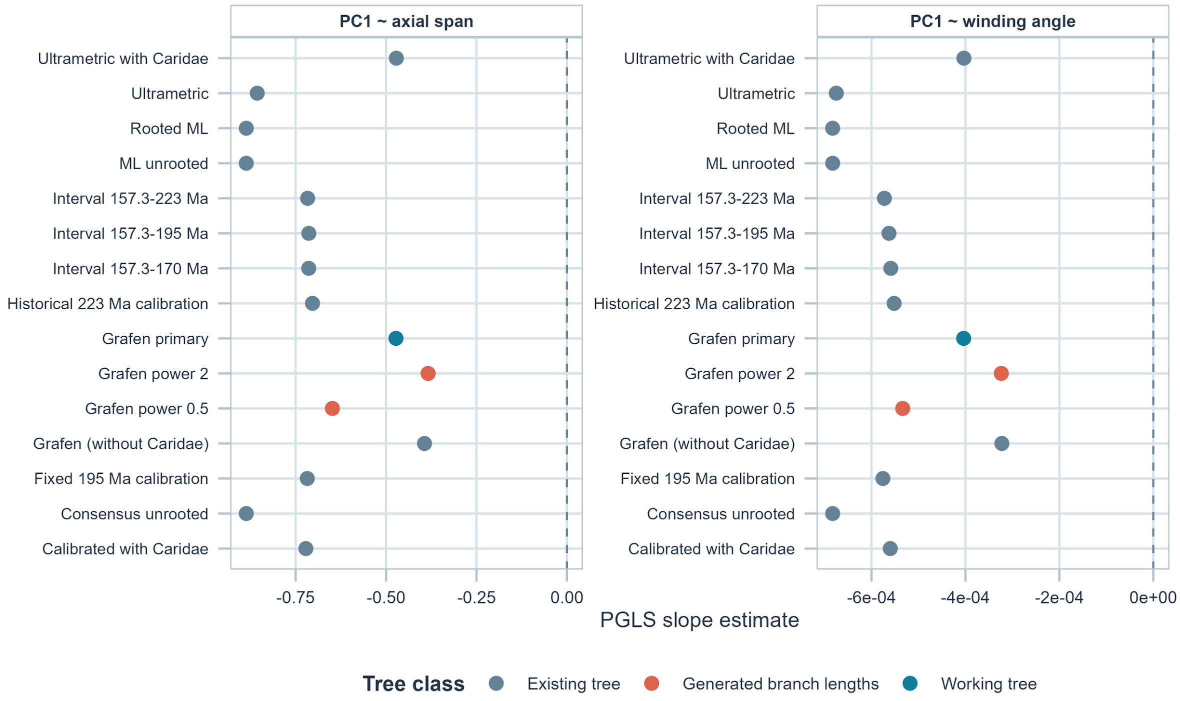


**Supplementary Figure 17. Sensitivity of PGLS slope estimates across alternative tree variants.**

PGLS slope estimates from the main dataset recalculated across 15 topology and branch-length representations. PC1 slopes for winding angle and axial span remained negative across all representations, but no focal relationship was significant after false-discovery-rate correction across the full sensitivity set. Point colours indicate tree class.

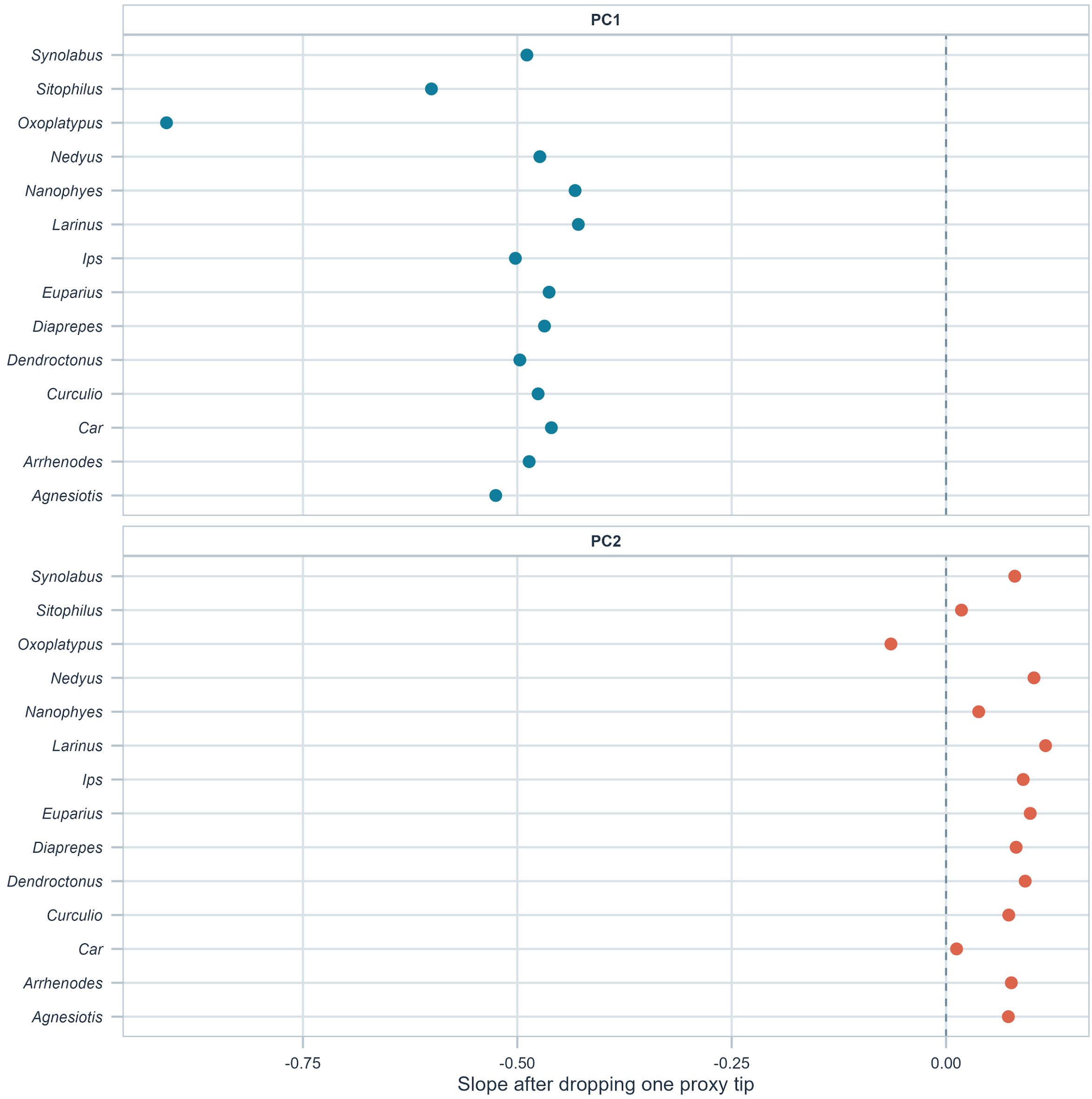


**Supplementary Figure 18. Leave-one-out robustness of PGLS slope estimates.** Leave-one-out sensitivity of the main-dataset PGLS slopes linking axial span to PC1 (upper panel) and PC2 (lower panel). Each point is a model refitted after excluding one of the 14 matched proxy tips. PC1 slopes remained negative across all exclusions, whereas PC2 slopes varied around zero.

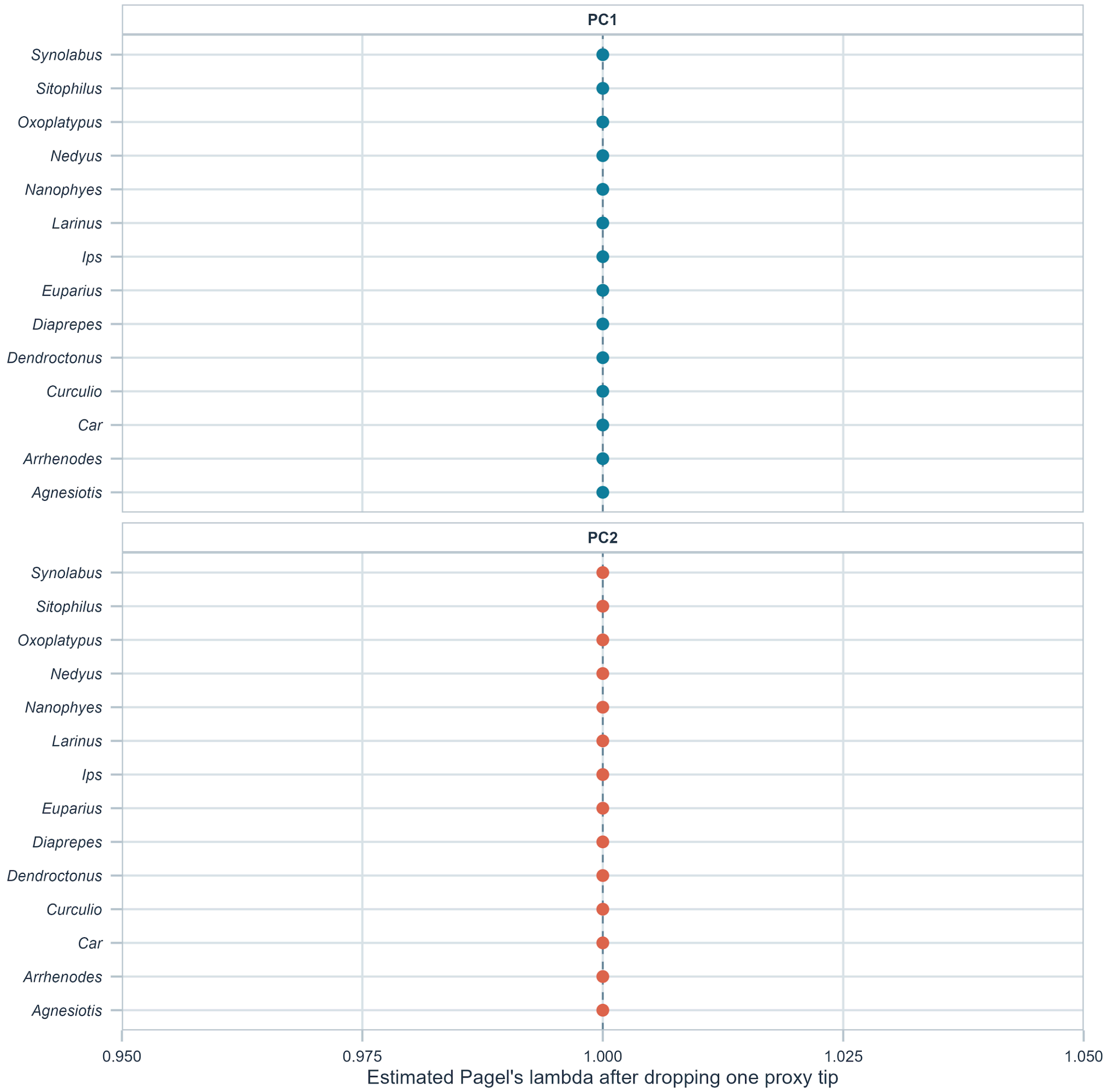


**Supplementary Figure 19. Leave-one-out robustness of estimated Pagel's lambda.** Leave-one-out estimates of Pagel's λ for the same main-dataset PGLS models linking axial span to PC1 (upper panel) and PC2 (lower panel). All leave-one-out fits reached the upper boundary estimate λ = 1.00, indicating that no individual proxy tip shifted the maximum-likelihood estimate away from the boundary.

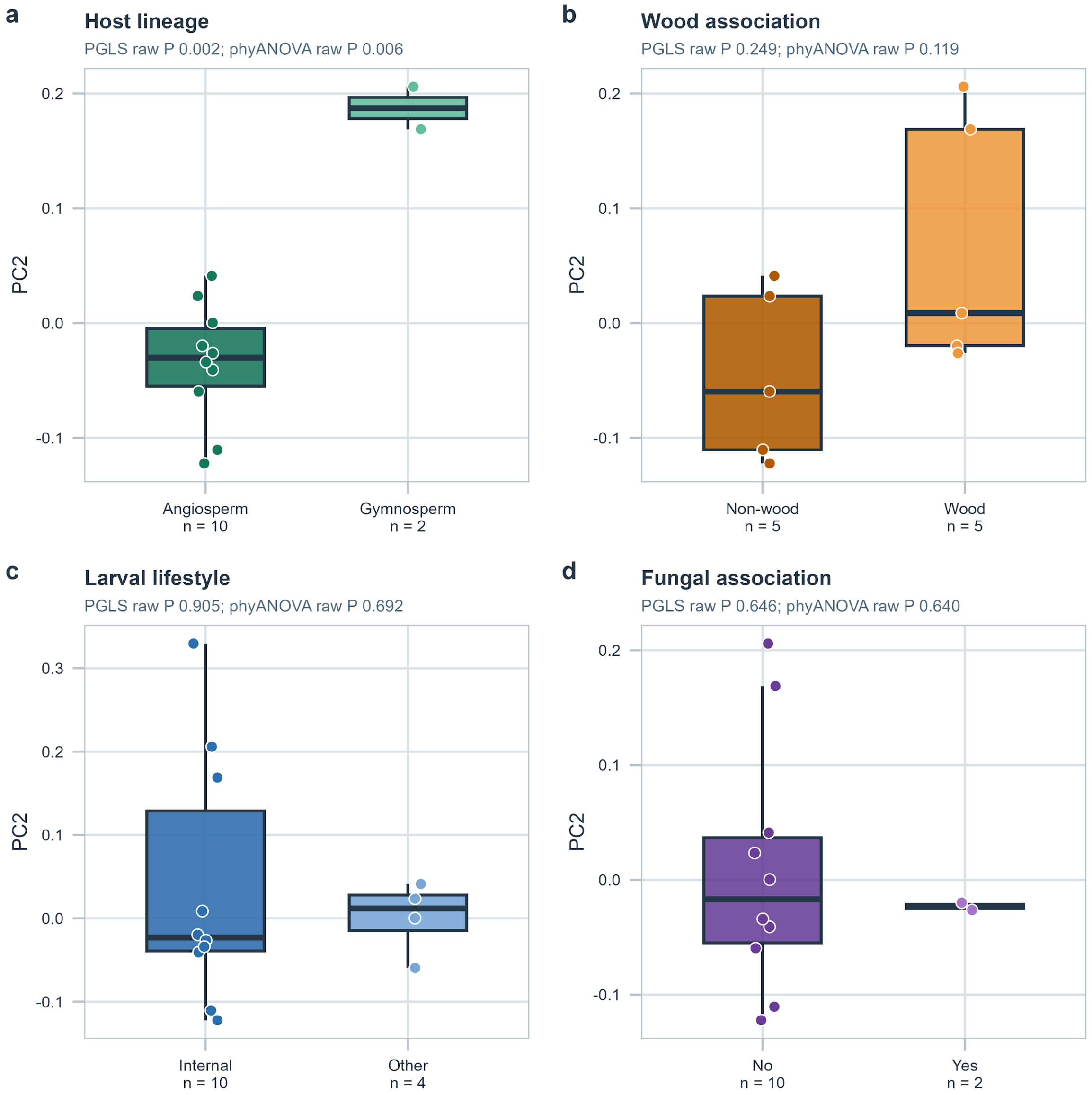


**Supplementary Figure 20. Ecological contrasts for PC2.**

Boxplots showing the distribution of PC2 across the four broad ecological predictors used in the comparative analyses: host lineage, wood association, larval lifestyle, and fungal association. Points indicate individual taxonomic proxy tips. For host lineage, the inferential comparison contains ten angiosperm-coded and two gymnosperm-coded proxy tips; mixed tips are descriptive only. Other group sizes are reported in Supplementary Tables 35 and 36. Panel annotations report unadjusted (raw) P values from the Grafen-primary PGLS and phylogenetic ANOVA; Benjamini-Hochberg FDR-adjusted values are given in Supplementary Tables 35 and 36.

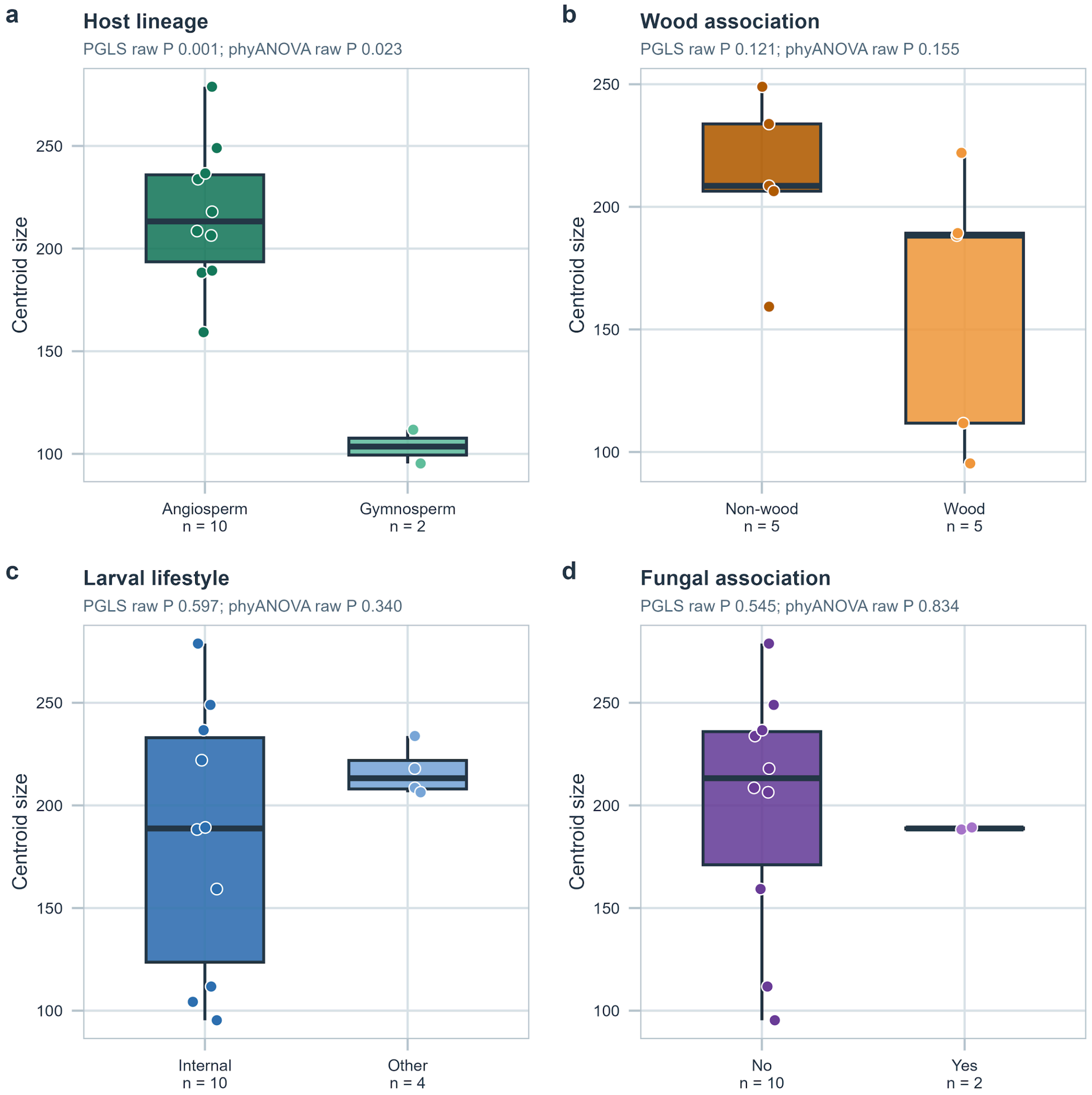


**Supplementary Figure 21. Ecological contrasts for centroid size.**
Boxplots showing the distribution of centroid size across the four broad ecological predictors used in the comparative analyses: host lineage, wood association, larval lifestyle, and fungal association. Points indicate individual taxonomic proxy tips. This figure provides a descriptive comparison of size structure across the ecological categories used in the supplementary comparative analyses. Panel annotations report unadjusted (raw) P values from the Grafen-primary PGLS and phylogenetic ANOVA; Benjamini-Hochberg FDR-adjusted values are given in Supplementary Tables 35 and 36.

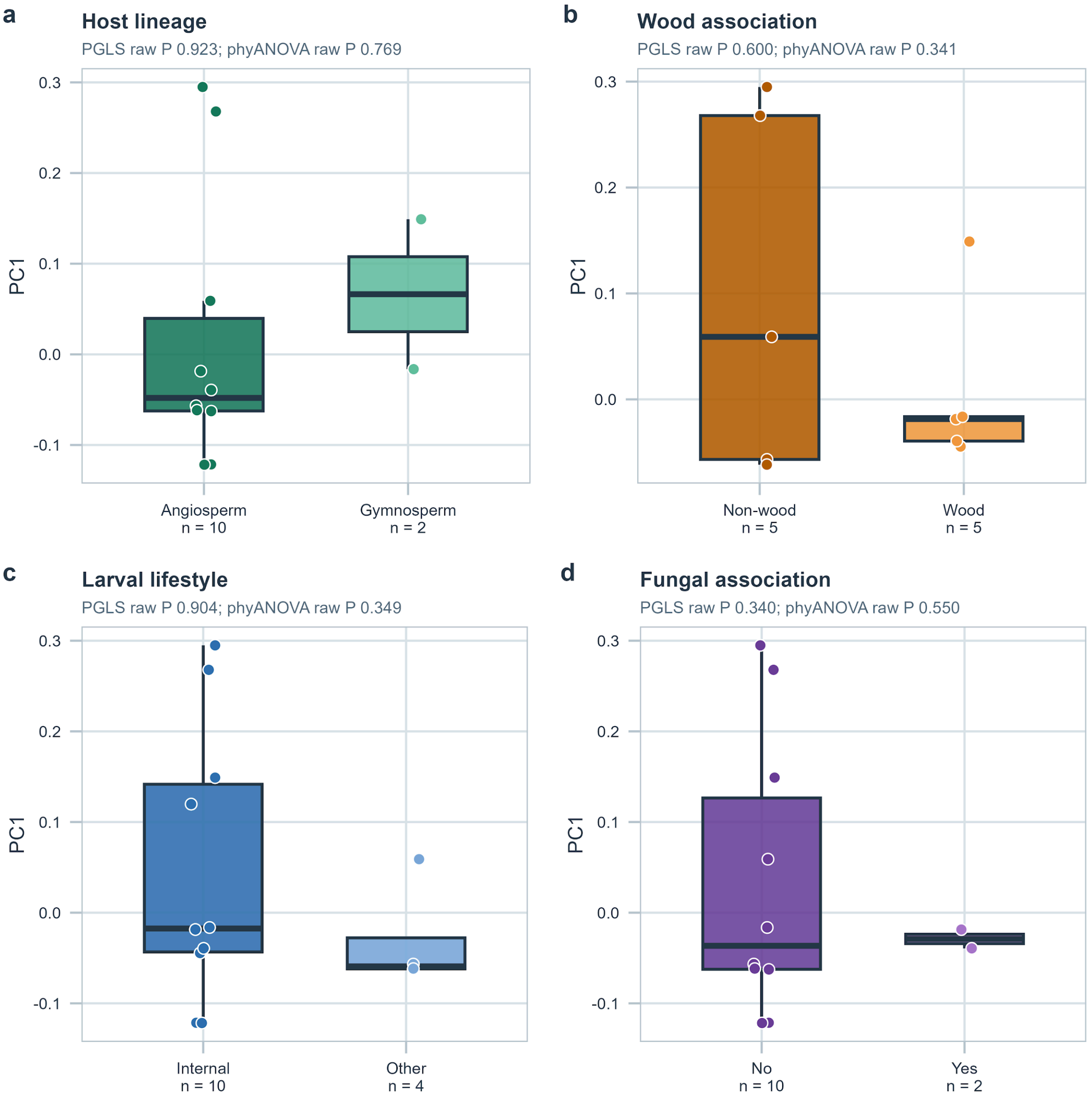


**Supplementary Figure 22. Ecological contrasts for PC1.**
Boxplots showing the distribution of PC1 across the four broad ecological predictors used in the comparative analyses: host lineage, wood association, larval lifestyle, and fungal association. Points indicate individual taxonomic proxy tips. These plots provide a descriptive overview of group overlap and effect direction for the leading axis of trochanteral shape variation. Panel annotations report unadjusted (raw) P values from the Grafen-primary PGLS and phylogenetic ANOVA; Benjamini-Hochberg FDR-adjusted values are given in Supplementary Tables 35 and 36.

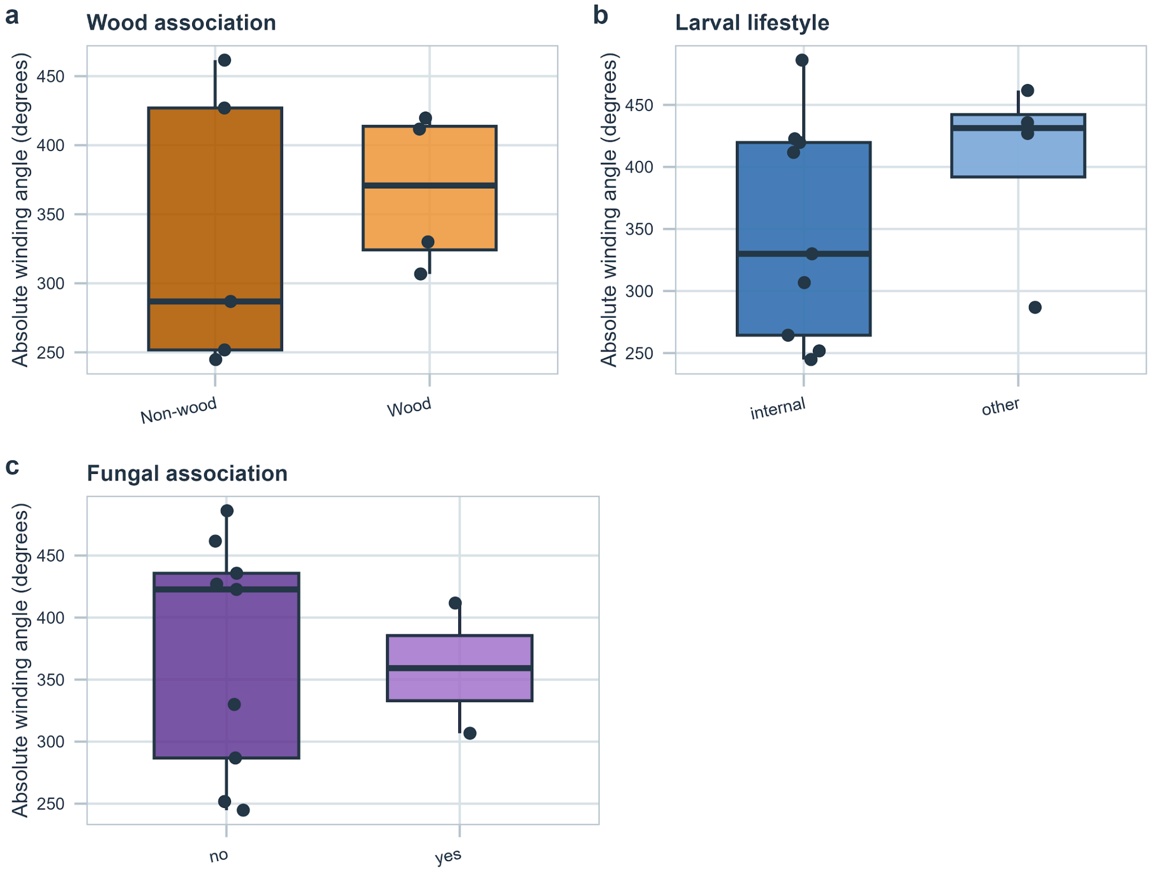


**Supplementary Figure 23. Ecological contrasts in the main dataset for fitted winding angle.**
Taxonomic proxy-tip means from the main dataset are shown across the three ecological predictors with estimable two-level contrasts: wood association, larval lifestyle and fungal association. Host lineage is omitted because only one retained gymnosperm-associated tip was available. The fungal-association contrast for absolute winding angle survived the within-framework FDR correction in PGLS (nine non-associated versus two associated proxy tips; FDR-adjusted P = 0.037) but not in phylogenetic ANOVA (raw and FDR-adjusted P = 0.853); it is therefore treated as exploratory and method-sensitive.

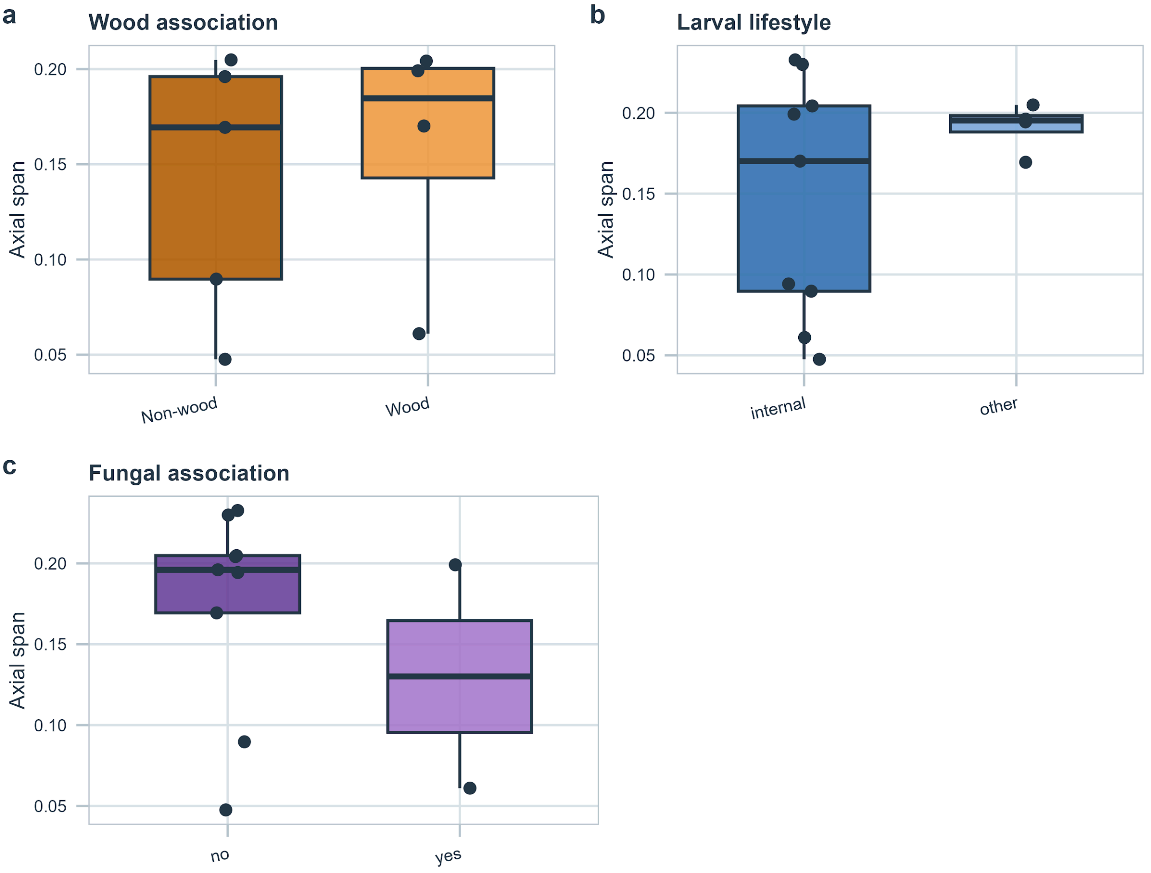


**Supplementary Figure 24. Ecological contrasts in the main dataset for axial span.** Taxonomic proxy-tip means from the main dataset are shown for axial span across wood association, larval lifestyle and fungal association. Host lineage is omitted because only one retained gymnosperm-associated tip was available. These panels are descriptive; no geometry association remained significant after FDR correction.

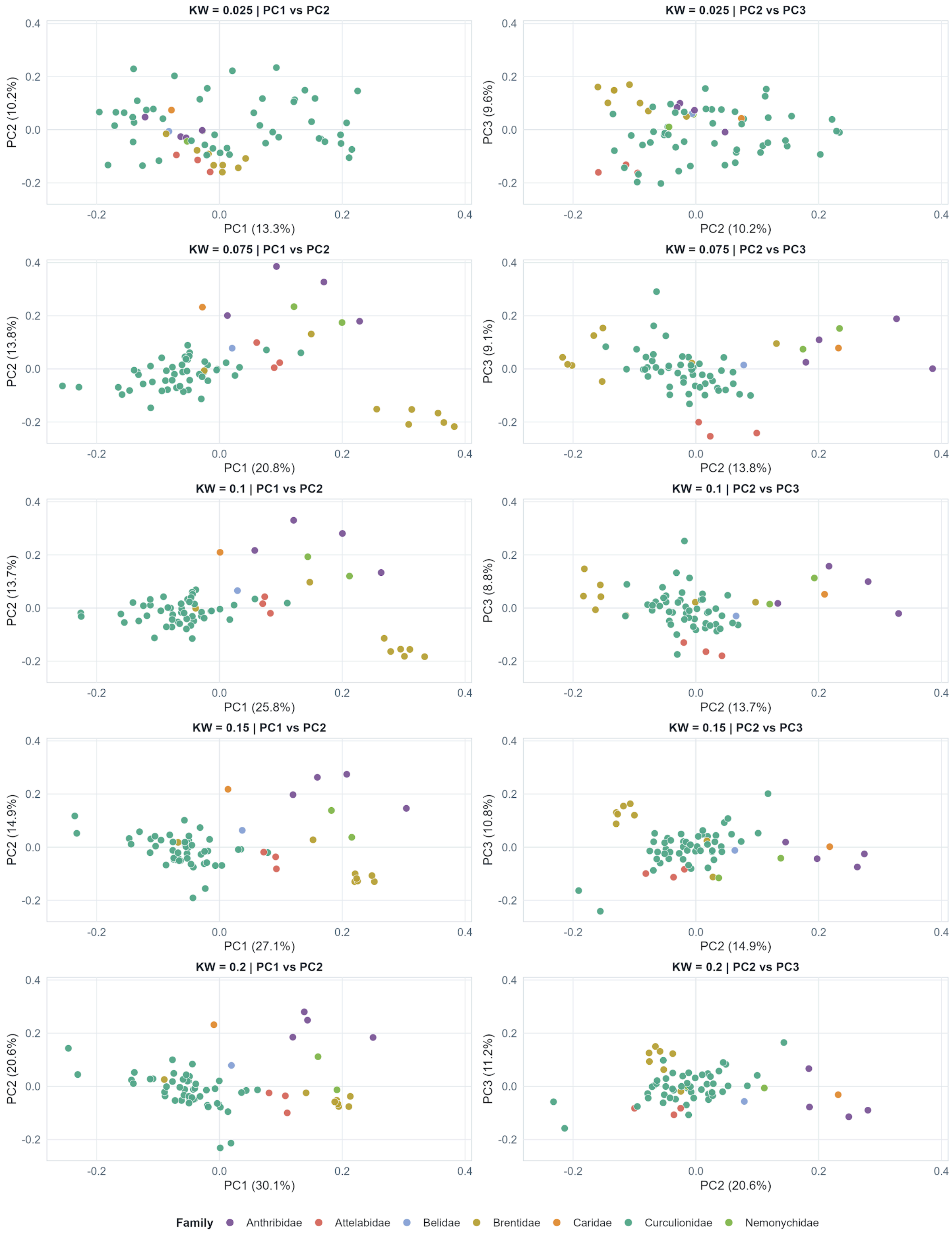


**Supplementary Figure 25. Sensitivity of morphospace structure to atlas kernel width.** Five archived 67-specimen atlas runs from the development-stage parameter comparison (kernel widths 0.025 to 0.2), shown in PC1-PC2 and PC2-PC3 space. Width 0.1 had first been selected visually; the quantitative comparison shown here was performed retrospectively. At width 0.1, full-space distance concordance was high with 0.075 and 0.15 (Spearman's $\rho$ = 0.986 and 0.973) but lower with 0.025 ($\rho$ = 0.355). These development-stage runs support a stable intermediate region, not a unique optimum or a final-dataset sensitivity result. Width 0.1 was retained unchanged for the final 68-specimen atlas after *Rhynchites cupreus* was added.

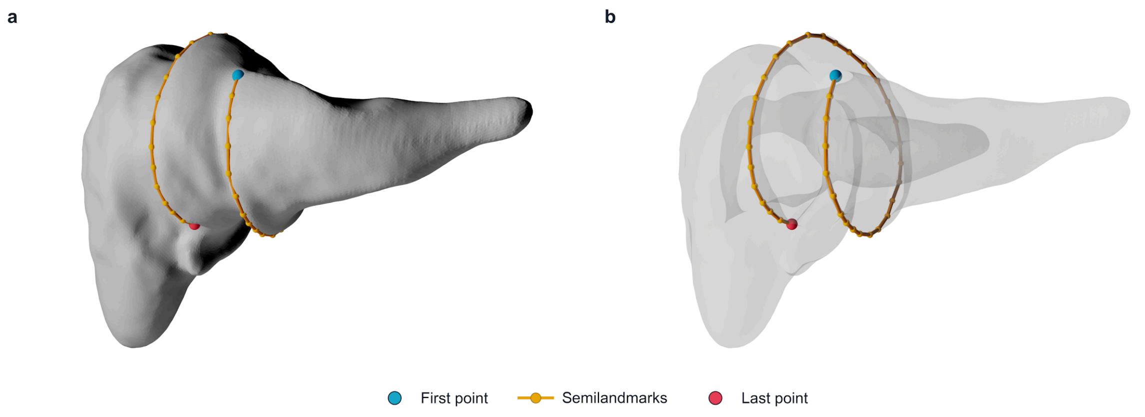


**Supplementary Figure 26. Semilandmark placement for screw joint geometry quantification.** Representative trochanter showing the ordered manually placed semilandmarks tracing the screw-like articular relief. The complete three-dimensional point series was fitted with a robust circular helix that continuously estimates axis position and orientation, radius, angular trajectory and axial rise. The blue and red markers denote the first and last points; orange markers and connecting lines show the intervening tracing sequence. Placement was performed once per specimen, so the conditional fit bootstrap does not quantify manual tracing repeatability.

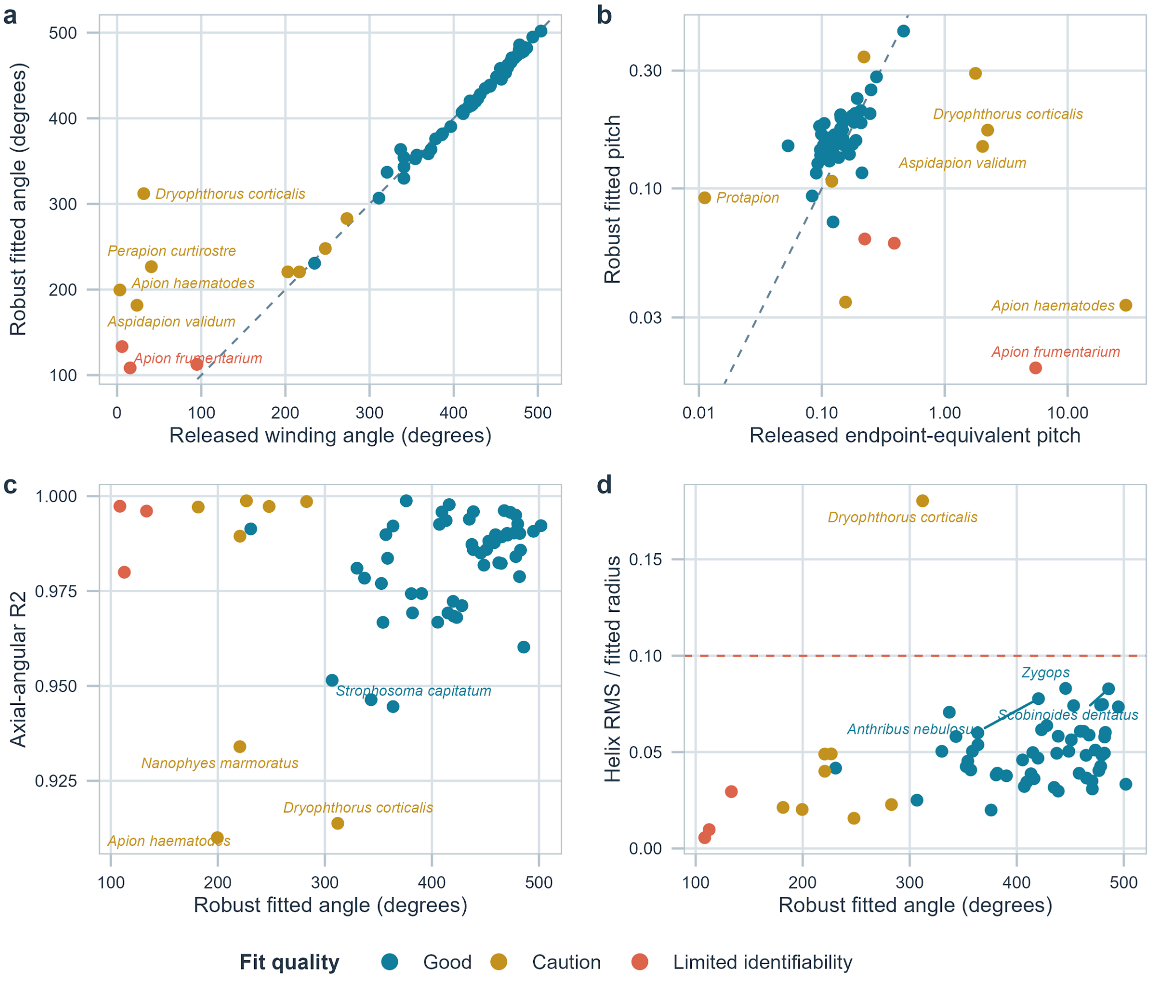


**Supplementary Figure 27. Quality control and sensitivity plots for screw joint geometry.**

Audit of the robust three-dimensional helix fit. Panels compare released and robust winding angle, released endpoint-equivalent and robust fitted pitch, axial-angular $R^2$, and helix RMS relative to fitted radius. The legacy angle estimates were closely reproduced overall, whereas fitted pitch differed strongly for several trajectories. *Dryophthorus corticalis* exceeded the relative RMS threshold used for the main dataset and was excluded. Point labels identify the largest discrepancies or diagnostic extremes.

### Supplementary Tables

**Supplementary Table 1. Specimen identifiers and phylogenetic tree-tip assignments.**
Specimen identifiers, corrected taxonomic assignments, family membership and mapping of 68 specimens to the 15 taxonomic proxy tips used in comparative analyses. The explicit proxy assignments, within-tip sample sizes and exact-versus-broader match status are provided in taxonomic_proxy_mapping.csv.

**Supplementary Table 2. PCA scores for all specimens.**
Specimen-level principal component scores derived from the landmark-free morphometric analysis of trochanteral shape. PC68 is retained only as a numerical-rank diagnostic and is not used in inferential analyses. This table corresponds to PCA_scores_with_specimen_id.csv.

**Supplementary Table 3. Variance explained by principal components.**
Variance explained by successive principal components of trochanteral shape. PC68 is retained as a numerical-rank diagnostic and explains effectively zero variance; inferential analyses use the 67 non-zero axes. This table corresponds to PCA_variance.csv.

**Supplementary Table 4. Sensitivity of morphospace and downstream analyses to linear and nonlinear ordination.**
Comparison of ordinary PCA and two RBF-kPCA parameterizations for ordination structure, family and joint-type organization, family dispersion and allometry. The kPCA family-dispersion calculation is a parallel exploratory diagnostic rather than an exact rerun of the primary bias-adjusted four-family PERMDISP. This table corresponds to kPCA_sensitivity_summary.csv.

**Supplementary Table 5. Summary statistics for the Hopkins clusterability analysis.**
Summary statistics for 100 randomized Hopkins evaluations of standardized PC1-PC5 scores ($n = 68$, $m = 6$, seed = 1), together with descriptive sensitivity across $m = 4$-$10$ using 1,000 evaluations and a documented independent seed per m. The files are hopkins_summary.csv, hopkins_randomized_values.csv and hopkins_m_sensitivity.csv. Repeated evaluations do not resample specimens and are not treated as formal inference.

**Supplementary Table 6. Clustering assignments across alternative methods.**
Specimen-level cluster assignments inferred from standardized PC1-PC5 scores by hierarchical clustering, k-means clustering and model-based clustering (mclust), together with the PC1-PC2 coordinates used for visualization. Hierarchical clustering and k-means recovered three clusters, whereas mclust selected five. This table corresponds to clusters_all_methods.csv.

**Supplementary Table 7. Family-level disparity in morphospace.**
Descriptive mean-squared-distance estimates from PC1-PC5 for families represented by at least three specimens, with bootstrap confidence intervals and $n = 4$ rarefaction. Global and pairwise inferential results are reported separately from bias-adjusted PERMDISP analyses of unsquared Euclidean distances to family centroids; pairwise P values were adjusted for the false discovery rate. Files: family_disparity_summary_pc1_pc5.csv,

family_disparity_global_test_pc1_pc5.csv, family_disparity_pairwise_tests_pc1_pc5.csv and family_disparity_rarefied_n4_pc1_pc5.csv.

**Supplementary Table 8. Observed and missing joint-character combinations among recurrent binary configurations.**
Frequencies of theoretically possible joint-character combinations among the 66 specimens assigned to recurrent binary configurations. The two atypical anthribid specimens are excluded from this combinatorial summary and described separately in the main text. This table corresponds to combos_overall_observed_vs_missing.csv.

**Supplementary Table 9. Whole-volume three-dimensional coxal wall-thickness metrics.**
Specimen-level coxa size, whole-volume local-thickness summaries, coxal-wall-opening state and mask-quality information used in the analysis. Files: coxa_3d_wall_thickness_metrics.csv and coxa_3d_wall_thickness_analysis_dataset.csv.

**Supplementary Table 10. Scaling relationship between whole-volume wall thickness and coxa size.**
Linear-model statistics and coefficients testing the relationship between median whole-volume coxal wall thickness and overall coxa size. Files: coxa_3d_wall_thickness_model_stats.csv and coxa_3d_wall_thickness_coefficients.csv.

**Supplementary Table 11. Coxal wall opening, whole-volume wall thickness and sensitivity analyses.**
Model summaries, group descriptives and sensitivity tests for the association between coxal-wall-opening state and whole-volume wall thickness after accounting for coxa size. Files: coxa_3d_wall_thickness_group_descriptives.csv, coxa_3d_wall_thickness_sensitivity_tests.csv and coxa_size_association_summary_stats.csv.

**Supplementary Table 12. Family-level distribution of recurrent joint-character combinations.**
Counts and proportional representation of recurrent joint-character combinations within families among the same 66 specimens. The two atypical anthribid specimens are excluded from this combinatorial summary and described separately in the main text. This table corresponds to combos_by_family_counts_and_props.csv.

**Supplementary Table 13. Specimen-level screw joint geometry measurements.**
Per-specimen robust helix measurements for the 63 trajectories in the main dataset, with all 64 successful fits supplied in a companion audit file. Variables include signed and absolute winding angle, axial span, fitted pitch, start-to-end distance, a separate model-derived axial-rise audit value, radius, residual metrics, descriptive diagnostic warnings and conditional-bootstrap intervals. Joint-type comparisons in the main dataset use 57 true screw-and-nut and three unopposed configurations.

**Supplementary Table 14. Main shape-geometry analysis results.**
Model-level and PC1/PC2 effect statistics for the principal robust shape-geometry analyses in the 63-specimen main dataset, with conditional-bootstrap summaries. The all-fit set is

retained only as an audit comparison in shape_model_uncertainty_summary.csv. Corresponding files are main_results.csv and shape_model_uncertainty_summary.csv.

**Supplementary Table 15. Regression summary for shape-geometry relationships.**
Regression statistics for robust shape-geometry and geometry-geometry analyses in the 63-specimen main dataset, including the axial-span model with PC1 and PC2. The tables also include descriptive regional sensitivity fits and geometry-geometry correlations. The corresponding file is regression_summary.csv; all 64 successful fits, main-dataset membership and descriptive diagnostics are documented in robust_helix_metrics.csv and the sample-flow tables.

**Supplementary Table 16. Core PGLS relationships between shape and screw joint geometry.**
Point estimates and conditional-bootstrap summaries for phylogenetic generalized least-squares relationships linking PC1 and PC2 to winding angle, axial span and fitted axial pitch. The principal axial-span models use 14 matched proxy tips in the main dataset; the all-fit set is retained as an audit comparison. Tree-variant results are reported separately in Supplementary Tables 29 and 30.

**Supplementary Table 17. Screw joint geometry statistics by joint type.**
Raw and Benjamini-Hochberg-adjusted univariate comparisons, dispersion test, fixed-seed PERMANOVA and conditional-bootstrap summaries for 57 true screw-and-nut and three unopposed configurations in the main dataset. Conditional-bootstrap summaries also retain the all-fit set as an audit comparison. The severe group-size imbalance makes all joint-type comparisons exploratory.

**Supplementary Table 18. Univariate allometry across all atlas-derived principal components.**
Axis-specific linear-model results testing relationships between log-transformed centroid size and all 67 non-zero atlas principal components, with Holm-adjusted P values. This table corresponds to allometry_univariate_all_PC_results.csv.

**Supplementary Table 19. Allometry of continuous geometry variables.**
Linear-model results for robust screw joint geometry as a function of log centroid size in the 63-specimen main dataset. Only absolute winding angle and its algebraically equivalent turn-number representation survived Holm correction; axial span and fitted pitch were unsupported.

**Supplementary Table 20. Full-shape and PC1-PC5 sensitivity RRPP allometry results.**
Permutation-based multivariate allometry results for the complete 67-axis atlas-derived shape space, together with the PC1-PC5 sensitivity analysis. These tables correspond to allometry_rrpp_multivariate_results.csv and allometry_rrpp_PC1_to_PC5_sensitivity_results.csv.

**Supplementary Table 21. Procrustes ANOVA results for multivariate allometry.**
Complementary Procrustes-based multivariate allometry results for the complete 67-axis atlas-derived shape space. This table corresponds to allometry_procD_lm_results.csv.

**Supplementary Table 22. Phylogenetically informed allometry results.**
Axis-specific PGLS for PC1–PC5 and geometry traits used log proxy-tip mean centroid size and maximum-likelihood Pagel's λ in [0, 1], without a fixed-λ fallback. Shape responses used all 15 tips; geometry responses used 14 tips after matching the 63 main-dataset specimens before aggregation. Tree-sensitivity and conditional-bootstrap results are supplied separately.

**Supplementary Table 23. Phylogenetic signal in continuous traits.**
Blomberg's K and Pagel's λ estimates for PC1-PC5 at all 15 shape-only proxy tips and for robust geometry traits at the 14 geometry-matched proxy tips in the main dataset, with FDR correction. Winding angle showed corrected signal, whereas axial span and pitch did not.

**Supplementary Table 24. Brownian-motion-conditional continuous ancestral state reconstructions.**
Raw-scale phytools::fastAnc node estimates, variances and 95% intervals under Brownian motion (BM) for PC1-PC5 and centroid size on the 15-tip shape-only dataset, and for absolute winding angle and axial span on the 14-tip main geometry dataset. The reconstruction_model column records BM for every row; analysis_tip_set and n_tips identify the trait-specific sample, and every trait displayed in Extended Data Fig. 5 is included. BM is a common reconstruction assumption rather than a model-selection result. This distinction is especially relevant to axial span, for which OU was top-ranked in the exploratory primary-tree univariate comparison. Geometry reconstructions are additionally treated as conditional summaries because of sparse proxy sampling, within-tip averaging and broader taxonomic assignments.

**Supplementary Table 25. Univariate evolutionary model fits.**
AICc comparisons of Brownian-motion, Ornstein-Uhlenbeck and early-burst models for principal traits in the main dataset. Rankings are exploratory and vary with tree choice.

**Supplementary Table 26. Multivariate evolutionary model fits.**
Diagnostics for multivariate evolutionary-model fits to PC1-PC5 and PC1-PC4 at 15 shape-only proxy tips in the main dataset. Optimizer convergence, Hessian reliability, raw AIC summaries and reliable-fit rankings are reported separately. Brownian-motion and early-burst fits were reliable in both comparisons, with Brownian motion top-ranked among reliable fits (early-burst ΔAIC = 2.79 for PC1-PC5 and 3.24 for PC1-PC4). The PC1-PC5 OU optimizer converged and had the lowest raw AIC, but its Hessian solution was unreliable; the PC1-PC4 OU fit failed. Raw OU support is retained for transparency but excluded from reliable-fit rankings. These incomplete comparisons were not used for biological inference.

**Supplementary Table 27. Full PGLS results for continuous trait-trait relationships.**
Results of phylogenetic generalized least-squares analyses across the current robust matrix of continuous predictors and responses. Geometry-dependent rows use 14 proxy tips after specimen-level matching to the 63-trajectory main dataset; shape-only allometry uses 15 tips. This table corresponds to pgls_continuous_vs_continuous.csv.

**Supplementary Table 28. ANOVA summaries for continuous PGLS models.**
One-degree-of-freedom coefficient-test summaries associated with the continuous trait-trait PGLS analyses. This table corresponds to pgls_continuous_vs_continuous_anova.csv.

**Supplementary Table 29. Summary of PGLS robustness across alternative trees.**
Summary statistics for key robust-geometry PGLS relationships in the main dataset across 15 topology and branch-length representations.

**Supplementary Table 30. Full PGLS robustness results across alternative trees.**
Complete PGLS results for the main dataset across 15 topology and branch-length representations.

**Supplementary Table 31. Summary of phylogenetic signal robustness across trees.**
Summaries of main-dataset phylogenetic-signal estimates across 15 topology and branch-length representations.

**Supplementary Table 32. Best-fit evolutionary model frequencies across trees.**
Frequencies of best-supported univariate evolutionary models across 15 topology and branch-length representations for the main dataset. Each n_trees value counts distinct tree representations, and the counts within every trait sum to 15.

**Supplementary Table 33. Brownian-motion-conditional root-state robustness in continuous ancestral state reconstruction.**
Summary of Brownian-motion (BM)-conditional reconstructed shape and robust-geometry root states for the main dataset across 15 topology and branch-length representations. Every tree-specific row and summary is labelled reconstruction_model = BM. These estimates test tree sensitivity under a fixed BM reconstruction assumption and do not compare evolutionary models.

**Supplementary Table 34. Leave-one-out robustness of PGLS models.**
Leave-one-out summaries for key robust-geometry PGLS models at the 14 matched proxy tips in the main dataset.

**Supplementary Table 35. Phylogenetic ANOVA results for ecological predictors.**
Phylogenetic ANOVA results for shape-only ecological contrasts of PC1, PC2 and centroid size and for geometry traits in the 14-tip matched main dataset. Host lineage was estimable for shape at 12 unambiguous tips (10 angiosperm and two gymnosperm) but not for geometry because only one gymnosperm-coded tip retained geometry. This table corresponds to ecology_phylogenetic_anova_results.csv.

**Supplementary Table 36. PGLS results for ecological predictors.**
Phylogenetic generalized least-squares results from caper::pgls for shape-only ecological contrasts of PC1, PC2 and centroid size and for geometry traits in the 14-tip matched main dataset; maximum-likelihood $\lambda$ fits were evaluated together with fixed $\lambda = 0$ and $\lambda = 1$ boundary candidates, and the valid fit with the highest likelihood was retained. Host lineage was estimable for shape at 12 unambiguous tips (10 angiosperm and two gymnosperm) but not for geometry because only one gymnosperm-coded tip retained geometry. This table corresponds to ecology_pgls_factor_results.csv.

**Supplementary Table 37. Descriptive ecological summaries by trait and factor.**

Group-wise descriptive statistics for PC1, PC2 and centroid size in the shape-only dataset and for geometry traits in the 14-tip matched main dataset. Host-lineage geometry rows are descriptive only because the inferential geometry contrast was not estimable. This table corresponds to ecology_trait_group_summary.csv.

**Supplementary Table 38. Non-phylogenetic ecological group tests.**
Non-phylogenetic group comparisons for shape-only ecological contrasts of PC1, PC2 and centroid size and for geometry traits in the 14-tip matched main dataset. Host lineage was estimable for shape but not for geometry. This table corresponds to ecology_nonphylo_group_tests.csv.

**Supplementary Table 39. Geometry-matched ecology matrix used for comparative analyses.**
Geometry-matched ecology matrix containing the analytical predictors and response variables for the 14 proxy tips in the main dataset. The table contains analytical variables only. Shape-only ecology tests use the 15-tip shape dataset, with 12 unambiguous tips in the host-lineage contrast, and are reported in Supplementary Tables 35-38. This table corresponds to ecology_analysis_input_merged.csv.

**Supplementary Table 40. Discrete stochastic character mapping summary.**
Summary statistics from 50 stochastic character maps of the coxal wall opening character across the 10 proxy tips with unambiguous states and their nine internal nodes. This table corresponds to asr_discrete_simmap_summary.csv.

### Supplementary Data

Supplementary Data 1**. Interactive three-dimensional trochanteral morphospace.**
Interactive 3D PDF showing the trochanteral morphospace with specimen meshes positioned according to their PC1, PC2 and PC3 scores and coloured by family. The file complements the two-dimensional projections shown in Fig. 2 and allows individual specimen models, family distributions and spatial relationships in the first three principal-component dimensions to be inspected interactively. A compatible desktop PDF reader with 3D annotation support is required; browser previews may show only the poster view. A plain-text accessibility and family-color key is included with release v1.3.2.

Supplementary Data 2**. Interactive three-dimensional heatmap reconstructions of principal-component extremes.**
Interactive 3D PDF showing the atlas mean shape and reconstructed negative and positive extremes of PC1-PC5. Surface colours indicate vertex-wise displacement from the atlas mean, with warmer colours marking regions of greater local deviation and cooler colours marking lower deviation. The file complements Fig. 3 by allowing the principal regions of trochanteral shape variation to be inspected from arbitrary viewing angles. A compatible desktop PDF reader with 3D annotation support is required; browser previews may show only the poster view. A plain-text accessibility and family-color key is included with release v1.3.2.

**Supplementary Data 3. Topology-based Grafen primary tree used for comparative analyses.**
Newick-format 15-tip taxonomic proxy tree used for the primary comparative analyses. Branch lengths were assigned from topology with Grafen's method and represent relative node depth rather than geological time. Results are conditional on proxy assignment and topology. The file is 01_primary_tree_grafen.tre.

**Supplementary Data 4. Additional tree variants used in phylogenetic sensitivity analyses.**
Collection of 14 alternative topology and branch-length representations used with the Grafen primary tree in a 15-tree sensitivity analysis. The alternatives include rooted, unrooted, ultrametric and Grafen transformations; trees with and without manually inserted Caridae; the archived historical 223 Ma calibration; a fixed 195 Ma calibration; and three interval-calibrated trees. File names, transformations, calibration provenance and fit diagnostics are documented in P01_Trees and S11_Robustness/Calibration_Sensitivity.

### Supplementary Video

**Supplementary Video 1. Continuous morphing sequence through the major axes of trochanteral shape variation.**
Animation of atlas-based shape deformation across the first five principal components. The sequence moves from the negative extreme of each PC through the atlas mean to the positive extreme, before returning through the mean and continuing to the next PC. The video visualizes the continuous shape changes underlying PC1-PC5 and is intended as an anatomical aid rather than as a temporal or evolutionary transformation sequence.
