## Supplementary figures and images for "Evolutionary diversification of the biological screw joint in weevils"

### Interactive 3D Heatmaps

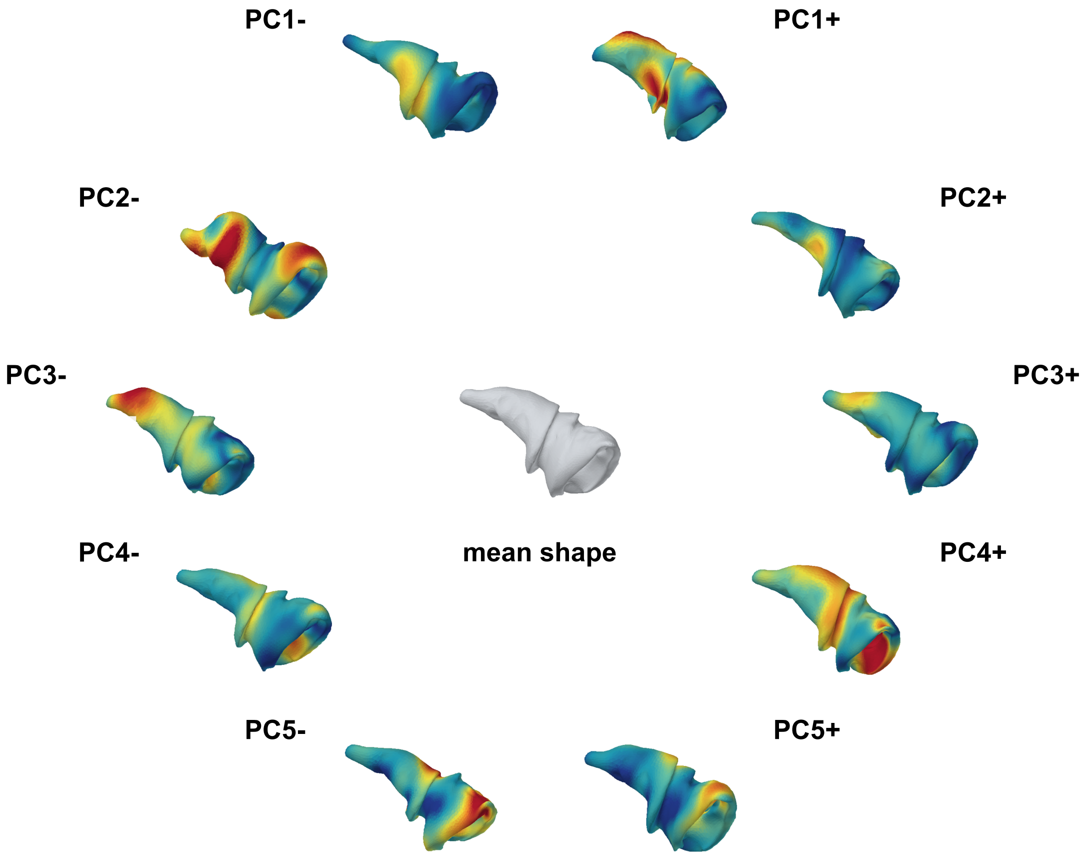

### Interactive 3D Morphospace

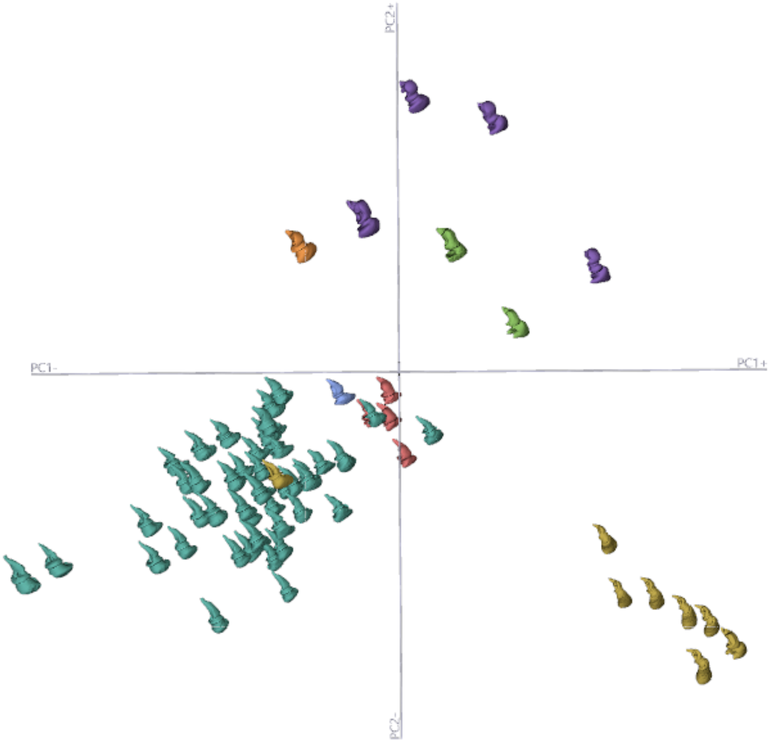
